# Apollo: A comprehensive GPU-powered within-host simulator for viral evolution and infection dynamics across population, tissue, and cell

**DOI:** 10.1101/2024.10.07.617101

**Authors:** Deshan Perera, Evan Li, Frank van der Meer, Tarah Lynch, John Gill, Deirdre L. Church, Christian D. Huber, Guido van Marle, Alexander Platt, Quan Long

## Abstract

Modern sequencing instruments bring unprecedented opportunity to study within-host viral evolution in conjunction with viral transmissions between hosts. However, no computational simulators are available to assist the characterization of within-host dynamics. This limits our ability to interpret epidemiological predictions incorporating within-host evolution and to validate computational inference tools. To fill this need we developed Apollo, a GPU-accelerated, out-of-core tool for within-host simulation of viral evolution and infection dynamics across population, tissue, and cellular levels. Apollo is scalable to hundreds of millions of viral genomes and can handle complex demographic and population genetic models. Apollo can replicate real within-host viral evolution; accurately recapturing observed viral sequences from an HIV cohort derived from initial population-genetic configurations. For practical applications, using Apollo-simulated viral genomes and transmission networks, we validated and uncovered the limitations of a widely used viral transmission inference tool.

## INTRODUCTION

Modern advancements in genomic sequencing have provided an unprecedented resolution that enables the study of viral evolution to venture into the within-host environment^1–3^. The era of epidemiological research ushered in by these technologies has revolutionized our understanding of viral evolution at host, tissue, and cellular levels^1,4–7^. However, the increasing volume and complexities of the data have outpaced current computational tools resulting in a bottleneck that inhibits our ability to fully utilize the potential of these vast new data^8,9^.

There are many simulation tools for viral evolutionary studies^10–17^. However, none natively scale to within-host, within-tissue, or within-cell resolution and thus may not accurately capture the intended evolutionary dynamics at finer resolution. Furthermore, as existing platforms are largely limited to single-core architectures, they cannot operate at scales sufficient to address the larger and more complex simulations demanded by the size and complexity of modern datasets. These limitations lead to an inability to account for transmission networks at within-host structures capturing pathogen genomic variations and phenotypic responses^11^.

To address these challenges, we developed Apollo, a simulator for studying viral evolution at scale at individual viral sequence resolution while accounting for population and within-host dynamics. We draw from the GPU-powered parallelization architecture CATE (CUDA-Accelerated Testing of Evolution) as well as conventional protocols for viral inference pipelines^18–21^.

Apollo is a forward-in-time simulator conducting evolutionary testing, analysis, and simulation at scale to bridge the gap between data and analysis. Apollo natively implements five hierarchical levels of an epidemic: network, host, tissue, cellular, and viral genome. Therefore, Apollo allows both scale and granularity in terms of epidemic configuration and simulation.

Through this paper, we present the design, implementation, and validation of Apollo. We demonstrate that Apollo is able to incorporate large pools of sequences and complex demographic and population genetic models and can replicate sequence evolution of viral sequences obtained from clinical cohort of individuals infected with HIV. Additionally, using an Apollo-generated gold-standard data set we validated and revealed the limitations of TransPhylo^16,17,22^, a popular viral transmission inference tool frequently used in amongst others in the COVID-19 pandemic^15,20,21,23^.

## RESULTS

### Software architecture spanning across five epidemiology hierarchies

Apollo’s novelty lies in its ability to span across five hierarchies of an epidemic: host contact network, individual host, tissue, cellular, and the viral genome itself (**Online Methods** and **Supplementary Note Figure S - 1**)^24–27^. Apollo’s efficiency is built on the computational framework of CATE^18^, a large-scale parallel processing architecture powered by the GPU, CPU and SSD. It is further enhanced by an out-of-core file structure supported by a novel parallelized search algorithm we refer to as Compound Interpolated Search (CIS). CIS enables identifying variants from the file space at O(log(log N)) time complexity^18,19^.

Epidemic spread is dependent on many interactions within a susceptible population. These interactions are captured via contact network graphs representing the spread of infection in the population^12,24^. Apollo supports a broad range of network models from random structures (e.g. Erdős–Rényi random graphs) to customizable networks that replicate real-world dynamics (e.g. Dynamic Caveman graphs) (**Supplementary Note Section 2.1**). Additionally, Apollo incorporates real-world scenarios such as explicit sampling schemes and their effects on a population.

Accurate epidemic modeling requires capturing within-host diversity^28^. Apollo implements support for heterogenous host populations with varying behavioral responses (such as quarantine upon diagnosis, treatment upon diagnosis and Lost-To-Follow-Up), immune and drug responses, as well as differences among within-host structures like tissues and their cellular environment (**Supplementary Note Section 2.2**).

Apollo supports detailed modelling of distinct viral populations in different tissues. It parametrizes complex tissue level dynamics using 13 parameters which govern aspects such as distinct generational phases governing viral population growth of individual tissues, cell affinity for viral attachment, and intra-host migration of viral particles (**Supplementary Note Section 2.2.3**).

At the cellular level Apollo explicitly models complex processes like viral genomic recombination, where the exchange of genetic material is dependent on the viral population occupying the same host cell (**Supplementary Note Section 2.3**). Apollo’s cell-level resolution allows users to configure characteristics representative of the individual’s infected tissues such as intra-tissue cell populations and specific roles of the tissues which can influence the infectiousness and mortality of an individual host (**Supplementary Note Section 2.2.5.1** and **2.2.5.2**). These capabilities enable support for numerous customizations including epidemiological compartment models ranging from Susceptible Infected Recovered (SIR) to Susceptible Exposed Infected Recovered Susceptible (SEIRS) and beyond^29,30^ (**Supplementary Note Section 2.2.5.6**).

The evolutionary landscape of viral evolution is modeled at the level of individual viral genomes^3,31^. Genomic variation resulting from evolutionary forces such as mutation and recombination is linked to phenotypic expression (**Supplementary Note Section 2.3**). This variation in expression introduces evolutionary pressures by affecting viral fitness, survivability, and mutation rates (**Supplementary Note Section 2.3.2.5 and 2.3.2.6**). Apollo accommodates segmented genomes allowing for multiple mutation and recombination hotspots within a single genome and each locus can be configured with its own base substitution models, mutation rates, and recombination factors (**Supplementary Note Section 2.3.2.6 and 2.3.2.7**).

Apollo navigates the complexities of simulating the five hierarchies via a three-phase architecture (**Figure 1 and Supplementary Note Section 3.3**):Parameterization, Initialization, and Simulation. In Parameterization, users configure Apollo across all five hierarchies using JSON scripting (**Figure 1A-D**). During Initialization, Apollo validates the parameters and sets up the contact network complete with heterogeneous hosts (**Figure 1E-H**). Finally, the simulation engine orchestrates the spread of the viral infection across the host population from one generation to the next. It manages the infection of the susceptible population while simulating evolutionary changes in viral genomes. Host behaviors and their characteristics are integrated, reflecting tailored host responses with the engine guiding the simulation across the modeled within host tissue and cellular environments (**Figure 1I-T**).

**Figure 1.**
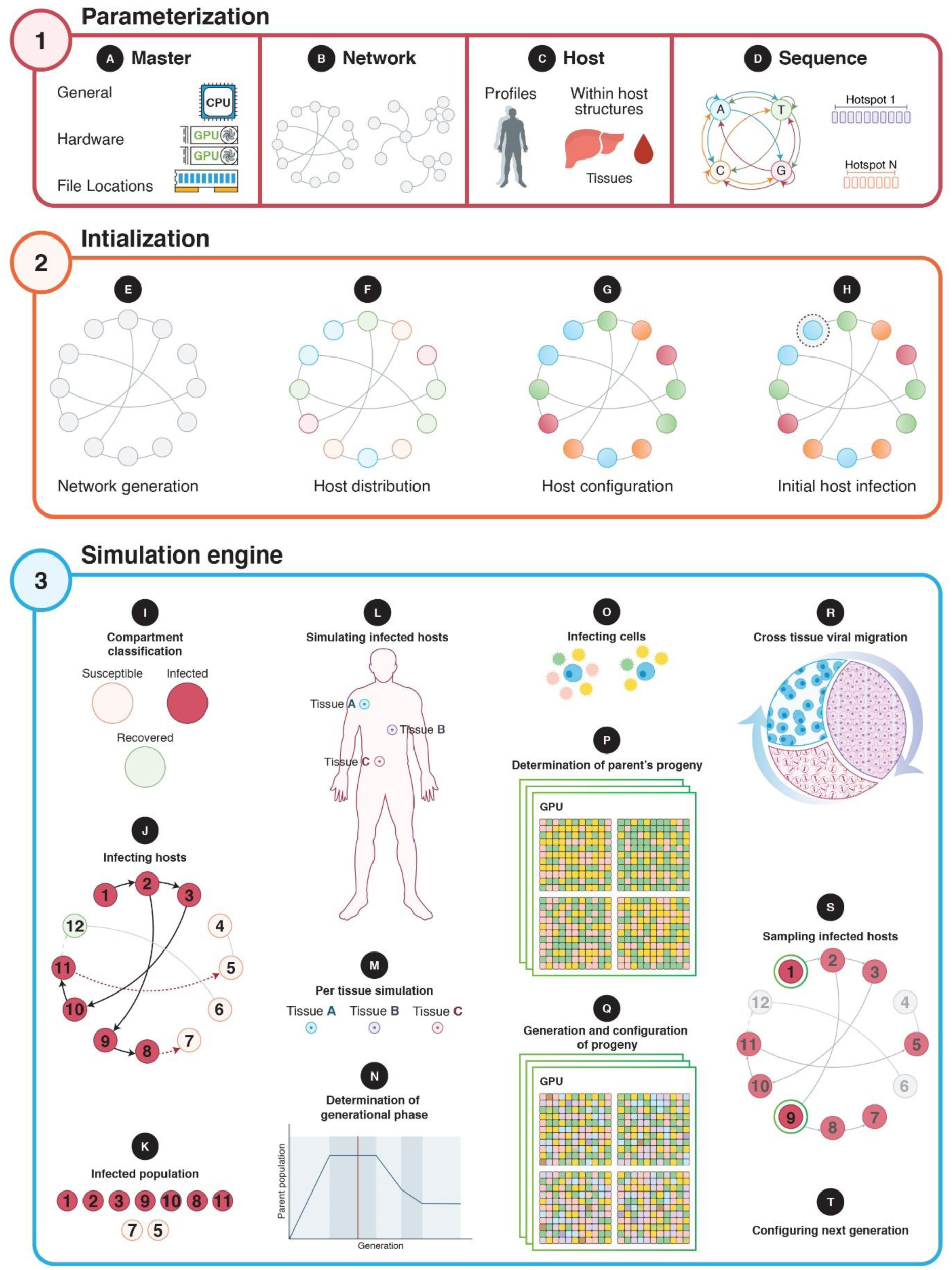
Overview of Apollo’s three phase architecture. Phase one: users configure the simulation with parameters for (**A**) computational resource allocation, (**B**) contact network, (**C**) host and within host characteristics and (**D**) viral genome. Phase two: simulation is initialized by (**E**) generating the contact network, (**F**) populating the network with hosts, (**G**) configuring individual host characteristics, and (**H**) selecting an initial host and infecting it with initial viral genome sequences. Phase three: Apollo processes the simulation one generation at a time. Beginning with (**I**) categorizing hosts to determine the infectious population who then (**J**) infect the susceptible population. From the (**K**) infected population, for (**L**) each infected host their (**M**) tissues are simulated sequentially considering (**N**) their current generation phase. (**O**) The virus infects the tissue’s cells and (**P**) initiate replication. (**Q**) Once the offspring have been mutated, assembled and configured they (**R**) migrate across tissues given the mechanics activation. After all infected individuals are simulated, (**S**) the sampling mechanism if selected is triggered, followed by (**T**) the setup of the next generation. The cycle continues until a simulation end condition such as obtaining a pre-defined number of host samples is met.

### Benchmarking Apollo’s capacity to scale

Apollo exhibited high linearity O(N) during benchmarking, where the processing time was a function of the within-host viral population size per individual host (**Online Methods** and **Supplementary Note Section 4.1**)^32^. This linearity was consistent for all test scenarios which included evolutionary mechanics of mutation, recombination and replicated across different classes of hardware resources.

Our baseline test simulations ran without evolutionary mechanics. The tests involved only viral reproduction while maintaining a constant parent population. We observed a regression gradient of 0.410 minutes per increase of 10,000 viral sequences in population size (R² = 0.995) (**Figure 2A and Supplementary Note Section 4.1.1**).

**Figure 2.**
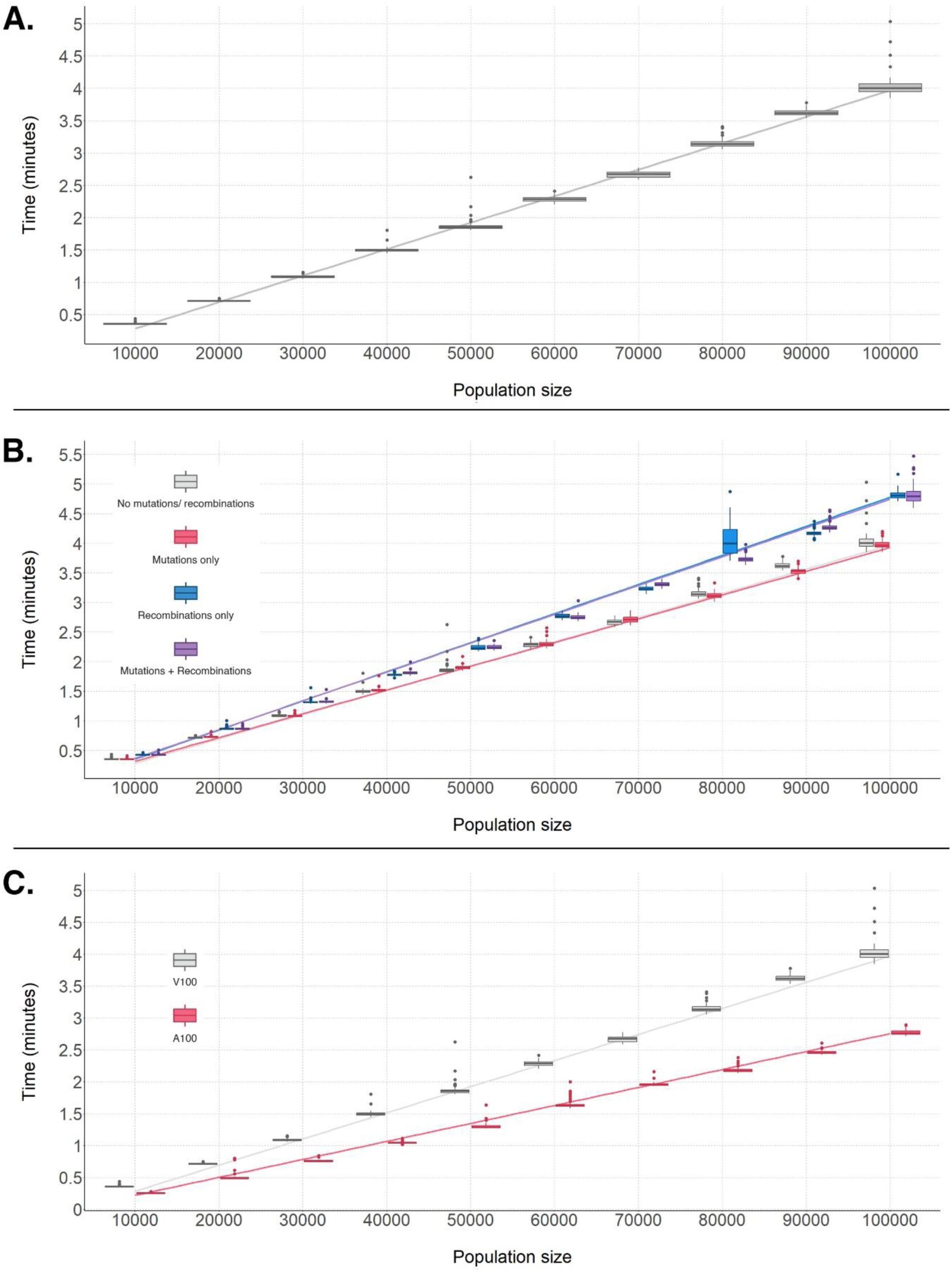
Scalability of Apollo in terms of time complexity under different test scenarios. (**A**) Evaluation of the baseline per generation processing time shows a linear increase proportional to population size. (**B**) An increase in the processing times is observed in the presence of the evolutionary mechanics of mutations (red) and recombinations (blue), or both (purple). (**C**) Apollo’s adaptability to the available hardware resources shows it was able to make use of the capabilities of the more powerful A100 GPUs (red) and increase performance above the baseline V100 (grey).

With the introduction of evolutionary mechanics, we observed slight variations in processing time in contrast to the baseline (**Figure 2B and Supplementary Note Section 4.1.2**). In the presence of only mutations, the regression gradient dropped to 0.401 (R² = 0.998). Conversely, with only recombination, the gradient increased to 0.491(R² = 0.991). When both mutation and recombination were present the gradient increased to 0.487 (R² = 0.997).

During the evaluation of Apollo’s hardware adaptability, we observed a significant decrease in processing time on the faster A100 GPUs compared to the baseline V100 GPUs (**Figure 2C and Supplementary Note Section 4.1.3**). The A100s improved the processing time by a factor of 1.454 to a reduced gradient of 0.282 (R² = 0.997).

### Evaluating integration and extension beyond the Wright-Fisher model

Apollo is built on a relaxed set of the assumptions from the Wright-Fisher (WF) model (**Supplementary Note Section 2.3.1**)^33–35^. Simulations testing the default parameterization of Apollo (**Online Methods**) saw the maintenance of the declared WF assumptions. (**Supplementary Note Section 4.2**). Apollo’s results corroborated with theoretical predictions of the rates of allele fixation due to genetic drift, including observations showcasing the increasing population size resulting in longer fixation times. Results from both tests (**Figure 3A and B**) were consistent with the predictions of standard WF model. Even though simulations started with varying haplotypes of equal frequency, the eventual fixation of a single haplotype population and the complete loss of all other haplotype populations was observed.

**Figure 3.**
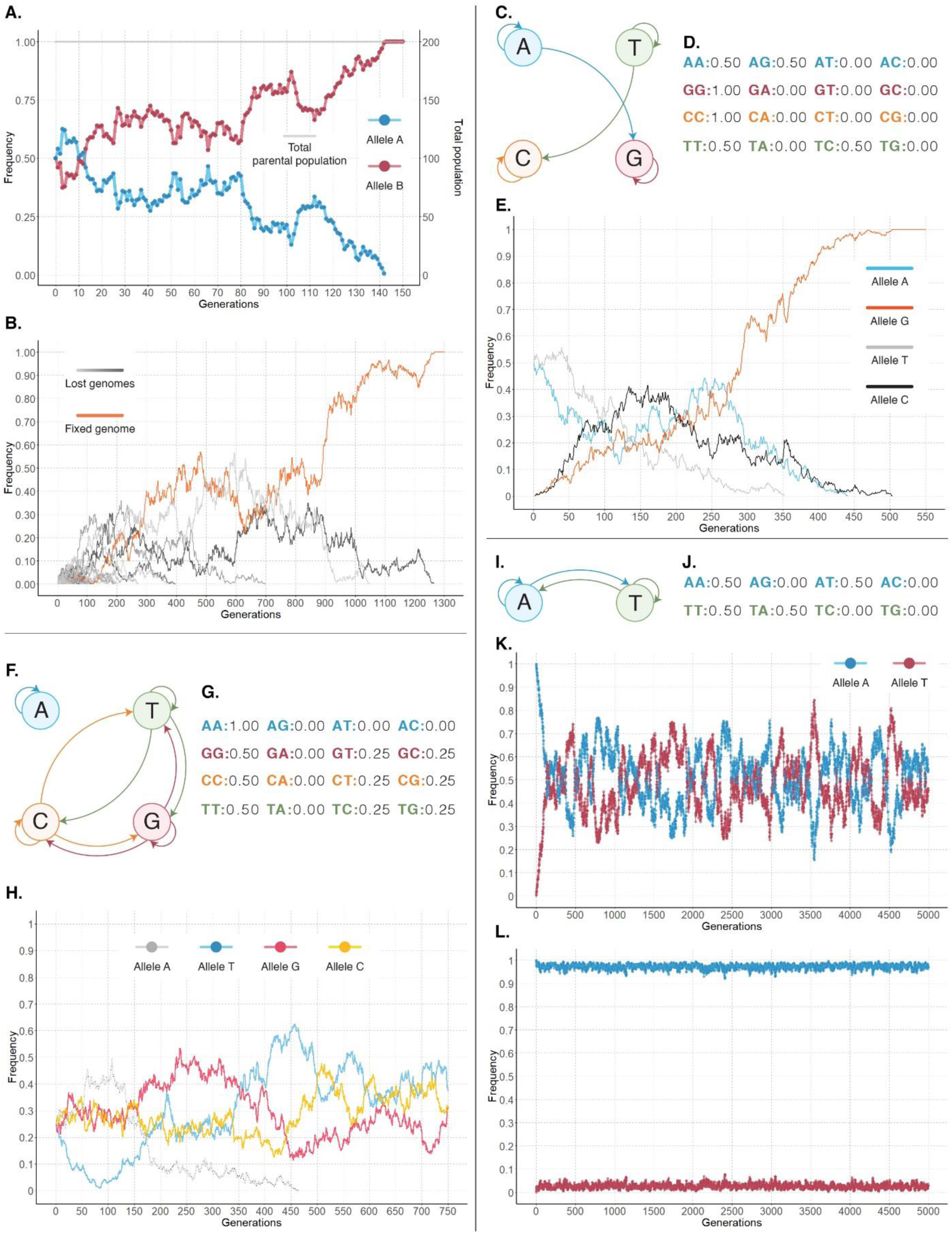
Consistency of Apollo simulations with theoretical predictions from relaxed Wright-Fisher (WF) models of population genetics. (**A**) Showcases the fixation of haplotype A (red) in the population while haplotype B (blue) becomes extinct under the forces of genetic drift while the parent population size (grey) remains constant. (**B**) Demonstrates the frequency changes of 100 haplotypes with one haplotype (orange) reaching fixation in the population while the remaining 99 haplotypes become extinct. (**C**) The Markov chain of the base substitution model used in the simulation of fixation of neutral mutations illustrates the possible transitions between nucleotides A, T, C, and G. (**D**) The corresponding transition matrix shows the probabilities of transitions between each nucleotide pair. (**E**) Showcases the changes of frequency of the alleles populations over 500 generations. Initial alleles A (blue) and T (gray) become lost, while mutated alleles G (orange) and C (black) rise in frequency, with G reaching fixation. (**F**) Represents the Markov chain used for the quasispecies simulation with (**G**) the transition matrix of the base substitution model. (**H**) Showcases the mutation-selection balance formed by the quasispecies of alleles T, G, and C (blue, red, and yellow respectively) allowing them to become fixed with allele A (grey) becoming extinct in the population. (**I**) Showcases the base substitution model’s Markov chain with the (**J**) transition matrix for the evaluation of selection forces. (**K**) Allele A (blue) viral sequences only exist at the start of the simulation and give rise to mutants of allele T (red) reaching a balance in the absence of selection forces. (**L**) Shows the frequency changes of alleles A (blue) and T (red) in the presence of the latter being deleterious. The lower frequency of the deleterious allele T is highlighted compared to the advantageous allele A.

We showcase Apollo’s capabilities to extend beyond the WF model (**Supplementary Note Section 4.3**) using mutation and selection forces. The introduction of neutral, irreversible mutations (**Figure 3C and D**) was conducted with the expectation of fixation of a mutated strain as theorized under the neutral mutation theory^36^. The simulations resulted in the rise of mutated haplotype populations followed by the subsequent fixation of one mutated haplotype. The remaining three populations including the two original populations were lost (**Figure 3E**).

Apollo’s accountability for selection forces was validated via the comparative analysis of change in population in the presence and absence of selection (**Supplementary Note Section 4.4**). We observed a balance in the mutated and original variants in the absence of selection (**Figure 3K**). Next, we expected a decline in the population of the mutated variant when a negative selection force was applied to it. As conjectured under negative selection theory Apollo’s simulations showcased the survival of the positively selected population inheriting the ancestral genome at a higher frequency while the negatively selected mutated variant existed at a lower frequency (**Figure 3L**).

Apollo’s adeptness at capturing complex evolutionary dynamics such as mutation selection balance was attested via a quasispecies simulation (**Supplementary Note Section 4.5**). Solving the fitness landscape for the eigenvectors and their subsequent eigenvalues revealed that two possible quasispecies dynamics should exist under the defined conditions. Apollo’s simulation consistent with the solution revealed the extinction of allele A while the haplotypes *T*, *G*, *C* achieved a mutation-selection balance reaching fixation of the quasispecies (**Figure 3H**).

### Simulation of HIV sequences corroborated HIV within-host status

We evaluated Apollo’s ability to simulate within-host viral dynamics, specifically the replication cycle of HIV as observed in infected individuals (**Online Methods** and **Supplementary Note Section 4.6**). Using Apollo, we modeled the progression of an HIV infection, leveraging metadata and HIV-1 viral sequences obtained from the Southern Alberta HIV Clinic, Canada as part of previous cohort studies^37,38^. Validation proved the successful recapture of sequences present in the real-world clinical samples using only the initial template sequences, base substitution models, recombination hotspots, and mutation rates.

The sequences consisted of the 701-base length clonal sequences from the viral genome’s Reverse Transcriptase (RT) pol region (GenBank: MN919177.1) obtained via Sanger sequencing (**Figure 4A and B**). The sequences collected from the Peripheral Blood Mononuclear Cells (PBMC) during the first four months of sampling contained 30 segregating sites. In total 192 segregating sites were identified to be present among all sequences spanning two years and four months collected across all five tissues (**Online Methods**).

**Figure 4.**
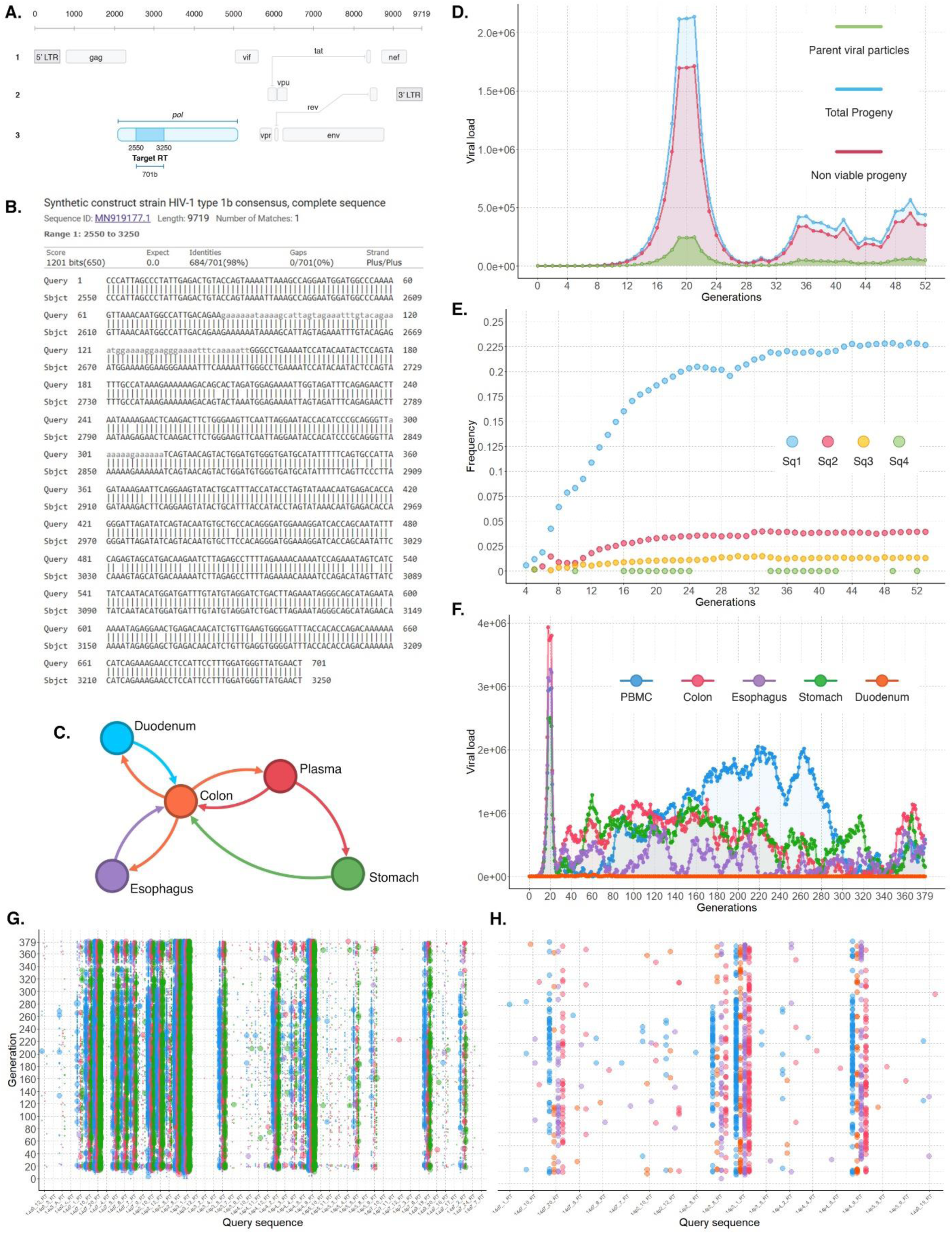
Experimental setup and Apollo’s replication of the real-world HIV infection in an individual infected with AIDS. (**A**) Genome map of HIV with the target region (dark blue) spanning 701 bases of the polymerase (pol) region (light blue) as identified by (**B**) the NCBI BLAST analysis. (**C**) Network of intra-tissue spread of the virus within the host, indicating movement between the five sampled tissues: duodenum, plasma, stomach, colon, and esophagus. (**D**) Temporal changes in the within-host viral population over 52 generations (four months) showcases the four stages of HIV infection: primary infection, acute HIV syndrome, clinical latency, and finally a slow rise in the viral load caused by the accumulation of high-fitness variants. (**E**) Frequency of four recovered sequences and their prevalence in the viral population across 52 generations. (**F**) Viral load changes in different tissues (PBMC, Colon, Esophagus, Stomach, Duodenum) over a period of two years and four months of infections. (**G**) Showcases the recaptured sequences from the simulation with two or fewer base mismatches compared to the target query sequences. Dot size represents sequence accuracy. The sequences are coloured by the tissue of occurrence (blue for duodenum, orange for colon, red for plasma, purple for oesophagus and green for stomach). (**H**) Subset of reconstructed sequences that perfectly matched clinical sequences from the HIV infected individual.

The first simulation test aimed to replicate the within-host dynamics experienced during the first four months of infection (**Supplementary Note Section 4.6.7**). The four canonical stages of HIV emerged from Apollo’s simulation (**Figure 4D**): a *primary infection* phase demonstrated an exponential increase in HIV viral load followed by *acute HIV syndrome*, then a drop in the viral load referred to as *clinical latency*, and finally a slow rise in the viral load caused by the accumulation of high-fitness variants that lead to opportunistic diseases and eventually *death*.

All four sequences present in the clinical data were recaptured during the course of the simulation (**Figure 4E**). The recaptured sequences perfectly matched those present in the clinical samples.

Next, we simulated the HIV within-host dynamics for a period of two years and four months of infection across all five tissues: PBMC, duodenum, colon, esophagus and stomach (**Supplementary Note Section 4.6.8**). This let us investigate the effect of cross-tissue migration on within-tissue viral diversity and population density (**Figure 4C**). We found that cross-tissue spread established a viral population in the Duodenum even though no viral sequences were present in the tissue at incidence of simulation (**Figure 4F**). Inspection of the simulated sequences against the clinical sampled sequences revealed that Apollo reproduced 19 sequences with 100% accuracy (**Figure 4H**), and a further 50 sequences with accuracies above 98.959% (a maximum of two base mismatches) (**Figure 4G**).

### Using Apollo to benchmark the accuracy of transmission prediction

Gold standard datasets are critical to benchmark the accuracy of predictions made by inference tools^39^. TransPhylo is a popular tool for inferring host-to-host transmission networks and predicting unsampled sources of infection and infection dates^16,17^ and has been used to evaluate epidemics including HIV AIDS and SARS-CoV-2’s COVID-19^15,20,21,23,40^. However, TransPhylo‘s inferences have not been validated against simulated datasets due to the lack of individual viral-resolution epidemic simulations with within-host dynamics. Apollo addresses this gap by providing epidemic simulations complete with the capture of within host dynamics. Apollo can generate who-infected-whom transmission networks, infection dates, and sampling information, making it ideal for benchmarking such inference pipelines (**Supplementary Note Section 4.7**).

We simulated an outbreak of a hypothetical virus in a population of 300 individuals. A total of 55 infected individuals were sampled at random via 50 sampling events. The population consisted of heterogeneous hosts of three types: non Lost to Follow-Up (LTFU) individuals (individuals who became non infectious upon sampling), complete LTFU (individuals who maintain their infectivity post sampling), and partial LTFU (individuals with reduced infectivity post sampling) (**Online Methods** and Supplementary **Note Figure S - 29**). Analysis of the transmission network revealed that hosts of the two LTFU populations remained infectious even when sampled (**Figure 5A**). We used the sampled sequences, and the host metadata produced by Apollo as the ground truth (**Figure 5B**).

**Figure 5.**
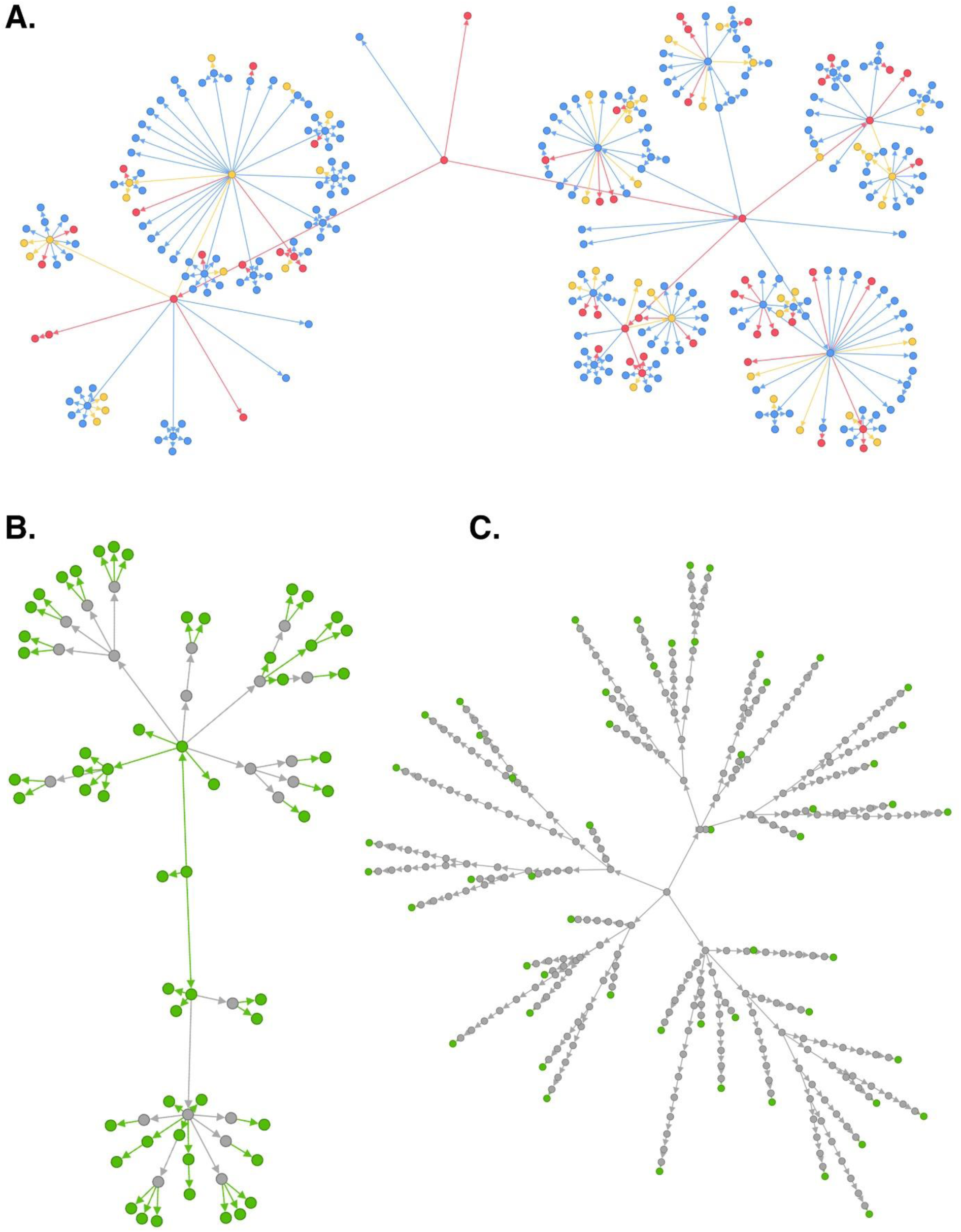
Using Apollo’s simulations as ground truth for evaluating the accuracy of TransPhylo predictions. (**A**) The complete transmission network as simulated by Apollo. Node colors represent the profile type of each host: blue for normal profiles, red for complete lost-to-follow-up (LTFU), and yellow for partial LTFU. (**B**) The abbreviated transmission network comprising of the sampled hosts (green) and the unsampled nodes (grey) between them. This is the true network to be predicted by the inference pipeline. (**C**) The predicted transmission network as generated by the inference pipeline. Green nodes represent sampled individuals, while grey nodes indicate inferred unsampled individuals. A clear overestimation of the population size along with incorrect inferences between host-to-host transmission can be observed in comparison to the ground truth.

The sampled sequences and their sampling time were then submitted to TransPhylo. It inferred the transmission network complete with unsampled sources of infection and the times of infection for the sampled individuals^16,20,21^. The successful execution of TransPhylo was evaluated using its MCMC tracer diagrams that showed convergence for the relevant parameters (**Supplementary Note Figure S - 30**). This is the standard practice for the evaluation of these tools^17,20–22,41,42^.

Despite the convergence of the MCMCs, we observed major deviations in the TransPhylo pipeline’s inferences in relation to Apollo’s ground truth (**Figure 5C**). The first inconsistency was observed in the estimation of the Most Recent Common Ancestor (MRCA) of the sampled population (**Supplementary Note Figure S – 29C and D**). The pipeline inferred the MRCA to be around the year 1990 when in fact it was around 1993.

Additionally, the pipeline’s transmission network overestimated the population that involved the sampled individuals. It predicted the presence of 329 infected individuals (**Figure 5C**) in the population while the truth was only 77 (**Figure 5B**) (including both sampled hosts and the unsampled individuals between them). This divergence was further stressed by the incorrect predictions of who infected whom. In the prediction of infection dates, we found a mean absolute error of 87.7037 days from the ground truth with 9.3 days and 194.35 days for the 5^th^ and 95^th^ percentiles.

## DISCUSSION

In this work we have introduced Apollo, a high resolution viral simulator designed to model viral infection and transmission across five hierarchical levels^43,44^. Our approach leverages the large-scale parallel processing architecture of CATE^18^, with extensions via multi-GPU support. Apollo effectively captures network-level dynamics, cross-host transmission, and host-specific behaviors, extending from tissue to cellular and genomic variations^45–47^. The robust three-phase architecture of Apollo ensures that it maintains efficiency and scalability while modeling complex epidemic scenarios.

Apollo’s observed linear increase in processing time (O(N)) with increasing within-host viral populations demonstrates that scaling simulations to larger, more complex scenarios will be both manageable and predictable. It enables accurate estimations of processing times given the complexity of the simulations at hand. For researchers working on simulations and big data projects, this linear relationship translates to reliable planning for computational needs and processing times. Additionally, Apollo’s demonstrated ability to efficiently utilize evolving hardware technologies to improve simulation speed ensures that it will remain a relevant and effective tool with its efficiency improving with advancing computational resources.

The successful integration of the Wright-Fisher model demonstrates Apollo’s ability to accurately simulate fundamental evolutionary processes such as allele fixation and extinction, aligning with theoretical predictions^34,35,48^. Apollo’s simulations were consistent with the theoretical predictions even when expanding beyond the Wright-Fisher assumptions^35,49,50^. This demonstrates that Apollo’s simulations have an accuracy and reliability that strongly reflect real-world evolutionary dynamics, including the effects of mutation and selection.

Simulation of HIV infection that was consistent with the real biological data obtained from an individual receiving ART over a period spanning two years and four months highlights its ability to model complex within-host dynamics and genomic evolution with high fidelity^51^. This capability enables the evaluation of various pathogenic scenarios, including the effects of therapeutic interventions on viral behavior. By accurately predicting sequence evolution, Apollo offers valuable insights into viral evolution, pathogenesis, resistance mechanisms, and the impact of treatment strategies.

Apollo’s proven fidelity, demonstrated through the standard Wright-Fisher model, its extensions, and consistency with real-world scenarios, makes it a reliable benchmark for evaluating inference pipelines. By serving as a ground truth, Apollo’s simulations allow for precise assessment and refinement of these tools. Comparing the inferences made by tools such as TransPhylo against Apollo’s simulations revealed their limitations. For instance, there is a need for improved compartment modeling beyond basic SIR models, including more accurate assumptions to improve the prediction of infection rates and sampling times. These comparisons highlight areas for enhancement in the inference tools and provides insights into how complex models can impact their inference accuracy^39^.

The implementation of five epidemiological hierarchies encompassed in a three-phase architecture enables Apollo to be a comprehensive tool for simulating viral dynamics at scale. We are capable of supporting this novel proposition through our large-scale parallel processing architecture and out-of-core framework^18^. Through Apollo, we bridge previous limitations of studying the effects of within-host evolution on the population scale and provide a versatile and powerful tool to explore, analyze, and anticipate various epidemic scenarios with unprecedented speed and accuracy. The insights made possible by Apollo will help drive progress in areas of epidemic inference, understanding viral evolution and behavior as well as effects of public health interventions.

## ONLINE METHODS

### Design and development of population to viral genome resolution architecture

The five epidemiological hierarchies captured by Apollo are encompassed in three main modules: network, host, and genome (**Supplementary Note Section 2**). The mechanics, assumptions, and algorithmic implementations of these modules have been extensively validated. The organization of the five hierarchies into the three modules allows the seamless integration of the required components with Apollo’s large scale parallel processing architecture (**Supplementary Note Section 3.3**).

Beginning with the network module its focus is contact network generation and recording host-host interactions. In total five stochastic network graph models designed to cater to a wide range of diseases are provided (**Supplementary Note Section 2.1**). Three of these models are standard graph models used in epidemiology: Erdős-Rényi random, Barabási Albert, and standard Caveman models. In addition, Apollo comes equipped with two more models intended to capture real-world interactions more accurately via additional layers of flexibility (**Supplementary Note Section 2.1.4 and 2.1.5**). They are named the random model and dynamic caveman model and are extensions of the Erdős-Rényi and caveman graph models.

Hosts are the unit of infection in an infection transmission chain^22,29^. The host module processes the within-host environment and the behavioral patterns of each individual in the population (**Supplementary Note Section 2.2**). The tissue structures, intra-tissue cellular environments, and the cross-tissue migration of the hosts are managed by the module. The host module is responsible for determining the duration of infection, the rate of progeny generation, and the roles of the tissue structures such as those that allow entry and exit of the disease into and out of the susceptible individual. The configuration of the host module also enables support for four epidemiological compartment models: SIR, SIRS, SEIR, and SEIRS. In addition, the host module works in tandem with the network module to determine who infects whom in the population.

The genomic module manages the viral genomes including mechanics related to its evolution and phenotypic expressions (**Supplementary Note Section 2.3**). By default, the genomic module supports the Wright-Fisher model (**Supplementary Note Section 2.3.1**). However, through parameterization of the simulation model, all WF assumptions can be relaxed except for discrete nonoverlapping generations (**Supplementary Note Section 2.3.2**).

The evolutionary forces accounted for by Apollo include mutation and recombination (**Supplementary Note Section 2.3.2.6 and 2.3.2.7**). The genomic module enables segmentation of the genome based on hotspot regions that can undergo mutations and recombinations. These regions can be overlapping. Mutational hotspots can have their own independent clock models, mutation rates, and site substitution models. The resultant effects caused by genomic variation are in relation to fitness, survivability, proofreading, and effects on recombinational factors such as the probability of region to undergo recombination and the likelihood of a recombination hotspot being the selected template.

### Evaluating scalability by benchmarking processing time relative to viral population size

To evaluate how Apollo’s per-generation processing time scales in relation to the increasing within-host viral population. Three different test types were conducted (**Supplementary Note Section 4**). The first was a baseline line test, used to obtain a comparative foundation against which the rest of the tests could be compared. The second was the assessment of the change in processing time in the presence of the different evolutionary mechanics of mutation and recombination as supported by Apollo. The third evaluated the change in processing time under different computational hardware. In all tests there is a single host, whose within host viral population was increased in increments of 10^5^ from 10^5^ to 10^6^. The processing time for each bin was averaged across 100 generations.

The baseline hardware consisted of Compute Canada’s Beluga cluster. At the time of testing, the cluster was equipped with NVIDIA V100SXM2 GPUs, Intel Gold 6148 Skylake CPUs and NVMe SSDs. We used 20 CPU cores and 50GB of RAM. For the first test the simulations were conducted under the WF assumptions with no mutations, recombination or selection.

The second test measured the change in processing times in the presence of 192 mutational hotspots, each with their own base substation models and mutation rates and 14 recombination hotspots. The mutation rates were set to follow a Poisson distribution of mean 0.3333 per generation. The hardware resources were consistent with the baseline using the Beluga cluster.

The third test was set up with the same configuration as the first with the exception being Apollo was being executed on Compute Canada’s Narval cluster. Narval comprised of improved hardware with NVIDIA A100SXM4 GPUs, AMD Milan 7413 CPUs and SSD storage^52^.

### Assessing Wright-Fisher model integration and advanced model extensions

Two simulations were configured to evaluate Apollo’s behavior on default parameters (**Supplementary Note Section 4.2.1**). At default Apollo’s simulations function under the Wright Fisher assumptions which include the absence of mutations, recombinations, selection forces and includes the maintenance of constant, within host viral populations sizes. The factors for generation time and progeny rate were the same as those used for Apollo’s baseline benchmark.

The first simulation consisted of two viral populations of haplotype A and haplotype B of equal frequency equaling to a total within host population of 200 viral sequences. The rate of progeny generation followed a negative binomial distribution of *n* = 10 and *p* = 0.55. The simulation was run for 500 generations.

In the second experiment we made the testing environment more robust by increasing the within host population size to 1000 virions and 100 unique haplotypes, each with an equal frequency. The simulation was run for 2818 generations.

To demonstrate Apollo’s capabilities to extend beyond the WF model using two experiments involving mutation and selection forces (**Supplementary Note Section 4.3.1**). To validate the mutation mechanic of Apollo we designed an experimental setup evaluating fixation under neutral mutations in absence of back mutation. The base substitution model was configured so that Haplotype A produced mutated haplotypes of base G, with base change probabilities of 0.5. Similarly, haplotype T would produce mutated haplotypes of base C. Both mutated haplotypes would not mutate further and produce only clonal progeny. Therefore, under the effects of genetic drift, one of the mutated haplotypes should reach fixation, and their frequency in the population should be affected by the parent haplotype.

For the evaluation of Apollo’s mechanisms capable of accounting for selection forces a comparative experiment was designed (**Supplementary Note Section 4.4**). First a control experiment where both the original and mutated strain had no selection advantage was implemented. The simulations were designed to start with a single loci genome of base A.

The implemented base substitution model allowed the mutation to allele T with a probability of 0.5 or it would remain unchanged with the same probability. The mutation rate followed a Poisson distribution of mean (*μ*) of 0.01. The viral population was maintained at a constant size of 1000 and the starting population comprised of only base A viral sequences. The simulation was run for 5000 generations. Subsequently, to examine the effects of selection on the above control a negative selection pressure was applied to the mutated strains containing allele T by reducing their probability of survival from 1 to 0.85.

To illustrate Apollo’s ability to capture complex evolutionary dynamics such as mutation selection balance we conducted a simulation based on Eigen’s quasispecies principle (**Supplementary Note Section 4.5**). The viral population’s genome consisted of a single base, which could be one of four bases A, T, G, or C. Genomes of allele A did not undergo mutation and produced clonal progeny. The remaining three bases T, G and C had a base substitution model where they will remain unchanged with a probability of 0.5 or mutate to either one of the other bases with a probability of 0.25 (**Figure 3F and G**). The mutation rate followed a Poisson distribution of mean (*μ*) of 0.01. Under these parameters the mutation landscape can be configured as shown by matrix *W* below.

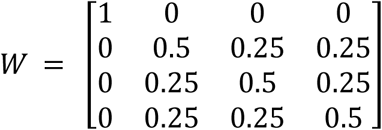

Solving for the matrix the eigenvalues and their respective eigenvectors are as depicted in **Table 1**.

**Table 1.** Solved Eigenvalues and Eigenvectors for the matrix W. In the eigen vectors corresponding with the matrix column 1 refers to base A, column 2 to base T, column 3 to base G and column 4 to base C.

| Eigenvalue | Eigenvector |
| --- | --- |
| 0.25 | [0 -1 1 0] |
| 1 | [1 0 0 0] |
| 0.25 | [0 -1 0 1] |
| 1 | [0 1 1 1] |

Based on the solution in **Table 1** either one of two quasispecies states will occur. These states are characterized by nonnegative values in their corresponding eigenvector. The expected quasispecies combinations would either be the fixation of allele A while the mutant species of T, G and C would become extinct or coexistence of alleles T, G and C forming a quasispecies through a mutation selection balance. In the latter allele A will become extinct and the quasispecies will become fixed in the population.

Following the simulations successful execution the downstream analysis of identifying the haplotypes and their frequency in each generation was conducted using Apollo’s utility function Haplotype Retriever.

### Simulation of HIV within-host infection dynamics while assessing fidelity through real world sequence data reproduction

The simulation of HIV infection within a host was conducted to demonstrate Apollo’s capabilities to capture robust epidemiological factors associated with the host, the within-host environment which spans across tissues, their cellular environment and the evolving viral genomes (**Supplementary Note Section 4.6**). Apollo’s ability to capture the complex interplay between evolutionary forces of mutation, recombination and selection was evaluated through the corroboration of simulation results using real world clinical sequence samples and meta data.

The HIV-1 sequence data used was collected as part of previous studies using the cohorts from the Southern Alberta HIV Clinic (SAC), Canada (**Supplementary Note Section 4.6.1**) (University of Calgary Conjoint Health Research Ethics Board (CHREB) approval NR# REB15-1941). The sequence data consisted of clonal HIV-1 viral sequences for the reverse transcriptase encoding region (RT) obtained over multiple time points. The samples were obtained from PBMC (Peripheral Blood Mononuclear Cells), and four gastrointestinal tissues (duodenum, colorectum, esophagus and stomach) and sequenced via Sanger sequencing^37,38^. The sample collection spanned a period of two years and three months (**Table 2**). The sequences used were obtained from on individual undergoing Anti-Retroviral Therapy (ART) monotherapy using Didanosine (DDI)^53–56^.

**Table 2.** Detailed temporal information on the sampling of the individual as well as the number of HIV RT pol sequences available and obtained from each tissue during each of the seven sampling events.

| Sampling number | Date | Sequences obtained per tissue |  |  |  |  |
| --- | --- | --- | --- | --- | --- | --- |
|  |  | PBMC | Colon | Stomach | Duodenum | Esophagus |
| 1 | 11-May-1993 | 12 | 9 | 9 | NA | 9 |
| 2 | 14-Sep-1993 | 11 | NA | NA | NA | NA |
| 3 | 11-Jan-1994 | 12 | 10 | 10 | 8 | NA |
| 4 | 21-Jun-1994 | 11 | NA | NA | NA | NA |
| 5 | 04-Oct-1994 | 12 | NA | NA | NA | NA |
| 6 | 02-May-1994 | 11 | NA | NA | NA | NA |
| 7 | 10-Oct-1995 | 12 | 10 | 10 | 10 | NA |

Evolutionary information in relation to the target gene region such as mutation rates and the positioning of recombinant hotspots was collected by referencing to existing literature. Fourteen hotspots of recombination predominant in this region were identified via a 2014 study conducted by Smyth *et al*^57^. The average Recombination Events Per Nucleotide per round of infection (REPN) was determined at 1.8 × 10^−3^. The mutation rate was configured to a Poisson distribution of mean 0.3333 per replication cycle^58,59^.

The identification of segregating sites or mutational hotspots was conducted using Multiple Sequence Alignment (MSA). We used MUSCLE (MUltiple Sequence Comparison by Log-Expectation) alignment via the MEGA (Molecular Evolutionary Genetic Analysis) software^60,61^. The base substitution transition matrix for each segregating site was then determined by analysis of the MSA data in conjunction with the time series sequence information.

The real-world analysis was conducted into two phases. Phase one was the simulation of infection spanning four months (or 126 days) and a single tissue, the PBMC. This period involves the time from the first to the second visit (**Table 2**) from May 11, 1993, to September 14, 1993. In phase two we expanded the analysis to encompass the entire sampled dataset (**Table 2**) using Apollo to simulate the entire infection, across all five tissues, from the first visit to the last visit. This period spanned 882 days or two years, four months, and 30 days. Apollo’s fidelity was evaluated by evaluating its capability to reconstruct sequences extracted at the last time point based solely on the sequences provided at the first sampling (11-May-1993).

The simulation was parameterised to encompass and reflect the within-host pressures exerted on the viral population by ART, based on documented effects. This was conducted by configuring phases of infection in each tissue. They were tailored to mirror the observed stages in HIV ART patients. Beginning with primary infection, characterized by an exponential growth or eclipse phase of the virus (0 to 4 weeks from infection), followed by acute HIV syndrome or primary infection phase (5 to 9 weeks from infection). Subsequently, clinical latency or chronic infection (9 weeks to 8 years from infection) ensues, followed by the onset of opportunistic diseases (9 to 11 years from infection), marked by a resurgence in viral load and eventual mortality^62^. As shown in **Table 3** these phases were delineated as timeframes based on the infected time of the individual.

**Table 3.** Details of the generational phases used for each tissue and the parameters used for each test. Since the Duodenum did not have a starting population, its phase configuration is different. A neutral phase is maintained till the tissue is occupied by viral particles caused by migration from other tissues.

| Test | Tissues | Phase count | Phase type | Time ratio | Distribution Parameters |
| --- | --- | --- | --- | --- | --- |
| **Test 1:**<br>4-months analysis | PBMC | 1 | Neutral | 0.35 |  |
|  |  | 2 | Stationary | 0.05 | Variance:10000 |
|  |  | 3 | Depreciation | 0.1 | Alpha:75,<br>Beta:75 |
|  |  | 4 | Stationary | 0.5 | Variance:10000 |
| **Test 2:**<br>2-years and 4 months analysis | PBMC,<br>Colon,<br>Stomach,<br>Esophagus | 1 | Neutral | 0.05 |  |
|  |  | 2 | Stationary | 0.0075 | Variance:10000 |
|  |  | 3 | Depreciation | 0.0125 | Alpha:75,<br>Beta:75 |
|  |  | 4 | Stationary | 0.93 | Variance:10000 |
|  | Duodenum | 1 | Neutral | 0.0750 |  |
|  |  | 2 | Stationary | 0.925 | Variance:1000 |

As the duodenum lacked sampled sequences from the initial sampling event (**Table 2**) its phases were adjusted to account for this by introducing a lengthened neutral phase. We introduce sequences into the duodenum by leveraging the integration of viral particle migration between tissues.

Apollo’s cross tissue spread mechanic was configured from the works by Chaillon *et al.* and Goyal *et al.* (**Supplementary Note Section 4.6.5**)^6,7^. The tissues across whom viral migration occurred were identified and the rates of spread were set using binomial distributions of *n* = 30 and *p* = 0.75. The initiation of migration was to generation 20 with the intention of providing sufficient time for the within-tissue viral population to amass. This meant that virus would start to spread from tissue to the next after 20 replication cycles had occurred in the simulation. Migration from the Duodenum was set to start after generation 40, since it only receives a viral population after cross infection from Colon.

Replication time was set to be 2.2 days with a standard deviation of 0.22. This was parameterised using a gamma distribution of shape 100 and scale 0.022^62–64^. The rate of progeny generation was set using a binomial distribution with *r* = 35 and *p* = 0.80^65,66^.

The reference survival rate for progeny was set at 0.15. The low survival rate was selected to reflect the challenges faced by the virus in the hostile host environment. Additionally it aligns with the viral particle counts observed in relation to the average progeny released by a cell and the actual viral load present in a host at any given time^62,67^. Based on the clinically sampled sequences a survival landscape was configured to favour viral sequences that were present in the real-world data. The affinity of virus to the cells of the tissues was configured using a gamma distribution of shape = 8 and scale = 6.

### Generating standard epidemiological datasets for quantifying the accuracy of inference pipelines

We begin by simulating an epidemic across a population of 300 individuals to demonstrate Apollo’s viability as a gold standard data generator. We then quantified and benchmarked the accuracy of predictions made by the inference pipeline against the simulation’s ground truth. The simulation was designed to represent a generalized viral infection without being specific to any particular disease. The hypothetical virus was 701 base pairs in length with 14 recombination hotspots and 30 mutational hotspots. The physical connections required to cause disease transmission was determined to follow that of an Erdős-Rényi contact network. The parameters were configured to capture a broad spectrum of properties associated with viral disease spreads.

The susceptible population was connected via an Erdős-Rényi contact network. The population comprised of 300 individuals with 0.75 probability of linkage between nodes. There were three types of host present. They were non lost to follow up, complete LTFU (Lost To Follow Up), and partial LTFU. Non LTFU represented individuals who upon being sampled will be removed from the infectious population. Therefore, the Lost to Follow Up (LTFU) individuals are those that remain infectious even after sampling. In contrast to non-LTFU hosts, LTFU individuals are considered to be able to have higher chance to transmit infections as they remain infectious after sampling. We segregated LTFU individuals into two categories. Those that remain completely infectious and those whose infectivity is reduced, pot sampling. They are labelled as Complete LTFU and Partial LTFU. Their percentages of distribution were 70%, 15%, and 15% respectively of the total population (**Supplementary Note Figure S - 29**). Detailed parameters for each profile type are outlined in **Table 4**. A sampling mechanic was used to conduct 50 sampling events at a rate defined using a binomial distribution of *n* = 10 and *p* = 0.05.

**Table 4.**
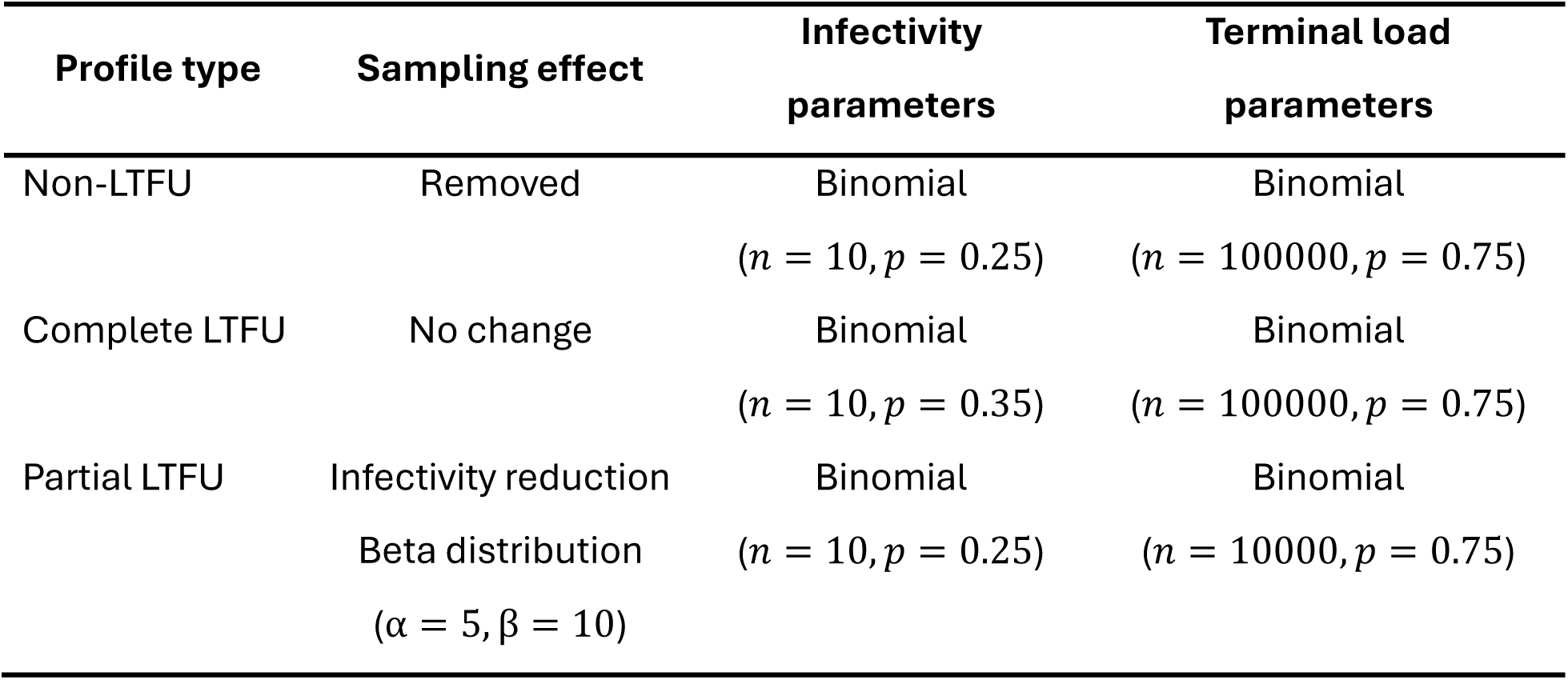
Details of the parametrization of the three profile types. The normal profile signifies individuals that are removed from the infectious population once they are sampled. Complete LTFU are individuals that continue to be infectious regardless of being sampled and identified. The partial LTFU individuals reduce their infectivity once sampled but will not be completely removed from the infectious population. The infectivity of hosts and their mortality by infection was dependent on profile type.

As depicted in **Table 4** to factor in the mortality of infected hosts we have reduced the terminal load of the partial LTFU population so that those whose viral load exceeds a particular threshold will cause the host to reach mortality.

The viral population parameters such as rate of replication, mutation, and recombination were made consistent with our previously described HIV analysis. The simulation was run with a start date of May 11^th^, 1993.

Once the simulation was completed the sequences obtained from the sampling mechanic were used to evaluate the inference pipeline. The sampled sequences came complete with host metadata, their time of being sampled, and infection time. Our inference pipeline was designed using the software TransPhylo^16,20–22^.

TransPhylo requires a time phylogenetic tree where the tip dates correspond to the sequences sampling time. We generated such a tree using the BEAST2 software. For the BEAST2 execution, we activated tip dates and used a gamma site model coupled with a GTR substitution model. The clock model was an optimized relaxed clock, and the prior was the birth-death skyline serial model. The MCMC chain was of length 10^9^. The final tracer diagram was evaluated by ensuring that each parameter had an Estimated Sample Size (ESS) greater than 200. The resultant trees were then summarised into a single tree using TreeAnnotator where the target tree type was set to maximum clade credibility and node heights to common ancestor heights. Burn-in was at the standard 10%^21^.

The resultant tree was then used by TransPhylo to predict the transmission network complete with unsampled sources of infection and the sampled hosts’ infection time. TransPhylo was parametrized according to the settings in **Table 5**.

**Table 5.** TransPhylo parameters for the generation of the MCMC tree and subsequent transmission predictions of who infected whom and infection dates.

| Parameters | Value |
| --- | --- |
| Infection rate shape | 1 |
| Infection rate scale | 0.99995 |
| Sampling rate shape | 1 |
| Infection rate scale | 0.5 |
| MCMC iterations | 100000 |
| Starting sampling probability | 0.0833 |

On evaluation of TransPhylo’s tracer diagrams and confirming their convergence (**Supplementary Note Figure S – 30B**) the predicted transmission tree and metadata were extracted. The inferences of who infected who along with the predicted infected times were evaluated against Apollo’s ground truth. Further evaluations were made in regard to the inferred population size and the Most Recent Common Ancestor (MRCA) in the sampled population.

## Supporting information

User Manual

## CODE AVAILABILITY

Apollo is freely available under the MIT license as part of the CATE software on multiple platforms including GitHub (https://github.com/theLongLab/CATE), Anaconda (https://anaconda.org/deshan_CATE/cate) and [Google Colab](https://colab.research.google.com/github/theLongLab/CATE/blob/main/Colab_CATE/CATE_on_Colab.ipynb). Apollo is complete with its own wiki (https://github.com/theLongLab/CATE/wiki/Apollo) and user manual (https://github.com/theLongLab/CATE/tree/main/Apollo_User_Manual). These documentations explain how to use Apollo and come complete with examples.

## AUTHOR CONTRIBUTIONS

Conceived the project: D.P., A.P., Q.L.

Designed the algorithm: D.P., C.D.H., A.P., Q.L.

Software Implementation: D.P.

Data analysis and benchmarking: E.L., D.P.

HIV data generation: F.vd.M., T.L., D.C., J.G., G.v.M.

Wrote the manuscript: D.P., G.v.M., A.P., with approval from all coauthors.

Supervised the project: A.P., Q.L.

Funding support: G.v.M., Q.L.

## FUNDING INORMATION

This work was funded by New Frontiers in Research Fund [Exploration NFRFE-2023-00291] (to Q.L.), NSERC Discovery Grant [RGPIN-2024-04679] (to Q.L.), Canadian Institutes of Health Research (CIHR) (to G.v.M.), National Health Research Development Program (NHRDP) (to G.v.M.), Eyes High International Doctoral Recruitment Scholarship (to D.P.), Alberta Innovates Graduate Student Scholarship 2021 (to D.P.), Graduate Faculty Council Scholarship (to D.P.), A.P. was funded by NIH grants [1R35GM134957] and [R01AR076241] to Sarah Tishkoff.

