## Supplementary Materials for "Apollo: A comprehensive GPU-powered within-host simulator for viral evolution and infection dynamics across population, tissue, and cell"

**LIST OF SUPPLEMENTARY FIGURES**

**Figure S - 1:** Depiction of the different levels of configuration that Apollo provides to enable the complete capture of the forces that govern an epidemic. (A) Generation of the susceptible populations' contact network. The colored nodes on the networks represent the different host profiles. Apollo provides a wide array of network models enabling the use of heterogeneous populations. (B) Configuration of transmission mechanics between hosts, including complete control over the disease transmission bottleneck. (C) Overview of the within-host dynamics from tissue to cell levels that govern evolutionary processes. Going beyond the host level Apollo enables the configuration of multiple tissues per host including control over tissue dynamics such as cell concentration and viral transmission between tissues. Furthermore, it accounts for the cellular level environment and the genomic variations and phenotypic responses of the viral particles. .......... 4

**Figure S - 2.** Two random graphs generated by Apollo. (A) Has a total of 50 nodes interconnected with each other and (B) forms a network with 250 nodes. As observed all nodes are connected to form a complete singular contact network. The nodes are colored by the node profile they belong to. There are three node profiles in the above networks (Blue, yellow, and green). .......... 10

**Figure S - 3.** Contact network graphs of 50 nodes generated under the Erdős-Rényi model with varying probability of node pairs being connected ($\boldsymbol{p}$). (A) Uses a $\boldsymbol{p}$ value of 0.01 and as observed a complete interconnected graph is not formed. In (B) and (C) all nodes are interconnected with each other, but the number of links progressively increase with the increasing probability ($\boldsymbol{p}$). The nodes are coloured by the node profile they belong to. There are three node profiles in the above networks (Blue, yellow and green). .......... 11

**Figure S - 4:** Contact network graphs of 50 nodes generated under the Barabási Albert model. The size of the nodes is ordered by their degree, nodes with larger degrees are depicted as larger in size. (A) The number of links ($\boldsymbol{m}$) is fixed at 1 forming the simplest form of the Barabási Albert model, the formation of Hubs can be seen. (B) Here too $\boldsymbol{m}$ is fixed but it is set to 2, and in (C) $\boldsymbol{m}$ for each newly incoming node is determined via a Poisson distribution where $\boldsymbol{\lambda = 2}$. The nodes are coloured by the node profile they belong to. There are three node profiles in the above networks (Blue, yellow, and green). .......... 13

**Figure S - 5.** Contact caveman graphs generated by Apollo with 50 nodes. (A) The nodes are distributed between five caves with ten nodes each. The nodes are colored by the cave they belong to. (B) The nodes are distributed among ten caves with five nodes in each cave. The nodes are colored by the node profile they belong to. There are three node profiles in the network (B) (Blue, yellow, and green)........................................................................................................ 15

**Figure S - 10.** Detailed depiction of the mutation process. (A) Apollo identifies the target hotspot ($m = 1$) region based on the start and stop positions. (B) Coloured coded bases with their letter and numeric index representations. (C) Coloured coded hotspot region representing its nucleotide configuration. (D) The respective hotspot region's parameters for mutation rate and base substitution. In the current generation's ($g$) mutation event, the Poisson distribution has drawn five mutations ($\gamma 1\_g$) to occur on the hotspot. (E) The five mutations occur across four base positions with base position 20 experiencing two consecutive mutations. (F) The mutated hotspot region with its new nucleotide base configuration....................................................... 41

**Figure S - 11.** The formation of recombinant progeny genomes. (A) Two parents, A and B that have coinfected a cell. Their genomes comprise of two recombination hotspots, each with their own nucleotide configurations. The resultant progeny (B) formed will be a chimera of the two parents A and B. Here the parental template is from parent B, who also contributes its genome to hotspot one, while hotspot two's region is configured using parent A's genome. ................. 43

**Figure S - 12:** Architecture comparison of the CPU and GPU. The CPU comprises of several cores. The CPU as a whole is optimised for parallel processing tasks with low latency. In contrast the GPU has smaller cores but the system as a whole is optimised with a denser core structure is for parallel processing. Therefore, GPUs with their extremely larger core count is able to outperform CPUs in tasks that require large scale parallel processing. ........................................................................ 47

**Figure S - 13.** Boxplot of the runtimes for each population size for Apollo conducted using the Compute Canada, Beluga cluster. We can see a linear increase in the average per generation processing time with the increasing bin size. ........................................................................ 57

**Figure S - 14.** Boxplot graph comparing the effects of mutation and recombination mechanisms on runtime. As observed the mutation and recombination mechanisms cause an increase in the per generation processing time. The mutation mechanism (red) appears to have a lesser impact on the baseline processing time (grey) compared to the recombination mechanism (blue). When both mechanisms are activated (purple) as expected there is an increase in the overall processing time, but only by a few seconds. ........................................................................ 58

**Figure S - 15.** Comparative analysis of the performance change of Apollo with V100 (grey) vs A100 (red) GPU architectures. We can observe a marked increase in Apollo's performance under the A100 GPU. ........................................................................ 59

**Figure S - 16.** Simulations under the standard Wright Fisher model for haploid viral genomes. The generations are abbreviated to show up to fixation (A) The changes in the two alleles' frequencies in the population across successive generations are shown. Allele A (red) reaches fixation while allele B (blue) is lost. The total parental population (grey) is constant across generations. (B) The changes in the frequency of 100 haplotypes. One genome reaches fixation (orange) while the others are lost at varying time points. ........................................................................ 61

**Figure S - 17.** Experimental setup and results of the first simulation for Wright Fisher model with neutral mutations. (A) Shows the Markov chain for the site model and (B) shows how the site model is configured within Apollo. As shown even though base G (Guanine) is not involved in the simulation it still has to be configured. (C) The results of the simulation show the variation in the frequencies of the different haplotypes. As observed the mutated haplotype containing Allele C (orange) reaches fixation eventually, while the original haplotypes (Allele A in dark grey and Allele T in light grey) cease to exist in the population. ........................................................................ 64

**Figure S - 18.** Experimental setup and results of the first simulation for Wright Fisher model with neutral mutations resulting in two mutated haplotypes. (A) Shows the Markov chain for the site model and (B) shows how the site model is configured within Apollo. (C) The results of the simulation show the variation in the frequencies of the different haplotypes. As observed the mutated haplotype containing Allele G eventually reaches fixation, while the original haplotypes A and T cease to exist in the population followed by the other mutated haplotype C. .............. 66

**Figure S - 19:** (A) Shows the Markov chain for the site model and (B) shows how the site model is configured within Apollo. As shown even though base G (Guanine) and C (Cytosine) are not involved in the simulation they still have to be configured. ...................................................... 68

**Figure S - 20:** The change in haplotype frequencies in the presence and absence of selection forces. (A) The change in haplotype frequencies in the absence of selection forces. As observed the mutated haplotype's frequency quickly rises to meet that of the original haplotype. They fluctuate around the 0.5 frequency value which matches the site model governing the simulation. (B) Shows the change in haplotype frequencies in the presence of selection. As expected even though the mutated haplotype does appear in the population it exists at a much lower frequency due to the negative selection pressure. ........................................................................ 69

**Figure S - 21.** Depiction of the experimental setup for the site model and results for the quasispecies experiment. (A) The configured site model for base changes during mutation and (B) the Apollo's site model format with the base transition probabilities for the site model. (C) A snippet of the simulation with the results of the change in frequency of each variant. As shown variants with allele A have become extinct with time, while variants with alleles T, G, and C have formed a quasispecies mutation selection balance and continue to persist in the population... 73

**Figure S - 25.** Change in viral population infecting the host as simulated by Apollo. We can see that the initial eclipse phase, followed by the acute infection phase and subsequently the period of clinical latency, and finally the cause of opportunistic infection leading to an eminent rise in the viral population once more. The viral particles that survived from the progeny to maturity to undergo cell attachment and reproduction are depicted in green. The progeny generated in each generation are shown in blue and out of them those that did not survive till reproduction are depicted in red. ............................................................................................................. 82
**Figure S - 26.** Details on the recovered four sequences via simulation. The graph depicts the frequency with which each sequence occupied the within host viral population at each generation that they were observed in. The sequence's frequency of occurrence also reflects the survivability fitness model that was applied to Apollo, which was designed using the real-world data. ........ 83
**Figure S - 27.** Change in the viral populations of each tissue with time as simulated by Apollo. The graph depicts the progeny viral population in each generation for each tissue. We can observe that the Colon and PBMC housed a majority of the viral population and the transmission between tissues helped maintain the viral populations of the other tissues as well................................ 85
**Figure S - 28.** Details on the reconstructed sequences by Apollo that match the sequences retrieved from the HIV infected subject via real world clinical data. The query sequence is shown in the x axis followed by the generation the sequence appeared in the y axis. The dots are coloured by the tissue where the sequence occurred. (A) Shows the sequences that were retrieved with two base mismatches or less in comparison to the target sequences. The size of the dot represents the accuracy of the sequence to the query sequence. A total of 50 sequences were retrieved via simulation. (B) Contains the subset of reconstructed sequences that were perfectly matched the sequences retrieved via real world clinical testing. A total of 19 such sequences were retrieved......................................................................................... 88

**Figure S - 29.** Contact network used for the simulation of the epidemic. Individuals are connected via an Erdős-Rényi graph model. The host types of non LTFU, complete and partial LTFU are distributed across the network at percentages of 70%, 15% and 15%. ...................................... 90

**Figure S - 30.** Analysis of the pipeline's BEAST2 and TransPhylo processes. (A) Depicts the generated tracer diagrams from BEAST showing the convergence of its inferences with ESS greater than 200. (B) TransPhylo generated tracer diagrams show that the algorithm has reached stabile convergence. (C) TransPhylo inferred transmission tree and (C) plot of incidences of sampled to unsampled cases over time. Observing (B) and (C) itself we can see that there is an error in the inferences as the start of infection has been predicted to be 1990 instead of 1993. Even the first occurrence of sampled individuals is placed into 1992. ............................................................ 91

**LIST OF SUPPLEMENTARY TABLES**

**Table S - 1.** Details of the 14 recombination hotspots. Their start and stop coordinates as positioned based on the complete HIV genome and the calculated rates of recombination...... 78

### SECTION 1. SOFTWARE INTRODUCTION

We present Apollo, our forward in time within-host viral simulator. Apollo is a state-of-the-art software solution capable of simulating viral infections at the individual viral sequence resolution complete with within host and population level dynamics.

Designed to extend beyond the standard Wright Fisher model Apollo encompasses evolutionary dynamics of mutation, recombination, replication, and forces of selection. It extends to the population capturing host behaviors and transmission with integration into the disease spread at population, host, tissue, and cell levels.

This is our newest addition to our established CATE (CUDA Accelerated Testing of Evolution) infrastructure[1]. Apollo is a command line software written in C/C++ and CUDA designed to run on Linux and Unix based Operating Systems (OS) equipped with NVIDIA CUDA enabled GPU hardware.

Apollo's simulations are highly configurable providing the user with the ability to parametrise a wide range of epidemic scenarios. Using large scale parallelisation technologies such as the GPU and CPU and when present the SSD, Apollo provides unprecedented resolution in a scalable, resource efficient, and accelerated manner.

#### 1.1. Software availability and documentation

Apollo is freely available as part of the CATE repo for download and use under the MIT License:

**GitHub:** https://github.com/theLongLab/CATE

**Anaconda:** https://anaconda.org/deshan_CATE/cate

Apollo comes with complete documentation that includes a user manual and GitHub Wiki. They are available at the following links:

**User manual:** https://github.com/theLongLab/CATE/tree/main/Apollo_User_Manual

**GitHub Wiki:** https://github.com/theLongLab/CATE/wiki/Apollo

#### 1.2. Defining Apollo

Apollo is a stochastic evolutionary simulator for haploid viruses, designed to study the progression of a disease across a population. Apollo's simulations move forward in time and aim to facilitate the design of highly dynamic and robust simulation models for studying within-host viral evolution and disease spread.

To achieve this level of granularity in epidemic capture Apollo is based on five main hierarchies (**Figure S - 1**). It begins at the population level with the generation of contact networks that govern the transmission of the disease across the population. Next, it factors the host model, with the flexibility to implement heterogeneity in host types[2,3], accounting from tissue to cellular level dynamics with features to implement host behavioral patterns[4]. Finally, Apollo accounts for evolutionary mechanics ranging from mutations, proofreading, recombination, and selection pressures[5–10].

Apollo by default is based on the Wright-Fisher model[8,11]. However, its power lies in its ability to go beyond the Wright-Fisher assumptions enabling the configuration of simulations that capture real world dynamics accurately[8].

Due to the complexity of Apollo's architecture, we will first begin by introducing Apollo and its capabilities (**Figure S - 1**) with the later sections diving into its architecture. Using a biological, statistical, and algorithmic approach we aim to provide the user with a detailed understanding of the different types of simulations that Apollo is capable of conducting.

As depicted in **Figure S - 1** the user is provided with capabilities to configure multiple aspects of an epidemic in great detail. Ranging from properties in the contact network, which then govern the subsequent host-to-host transmissions to the configuration of viral particle transmission between hosts and their tissues of infection[12]. It also provides granularity at the tissue and cell level enabling true simulation of an epidemic complete with within host evolutionary dynamics[13,14].

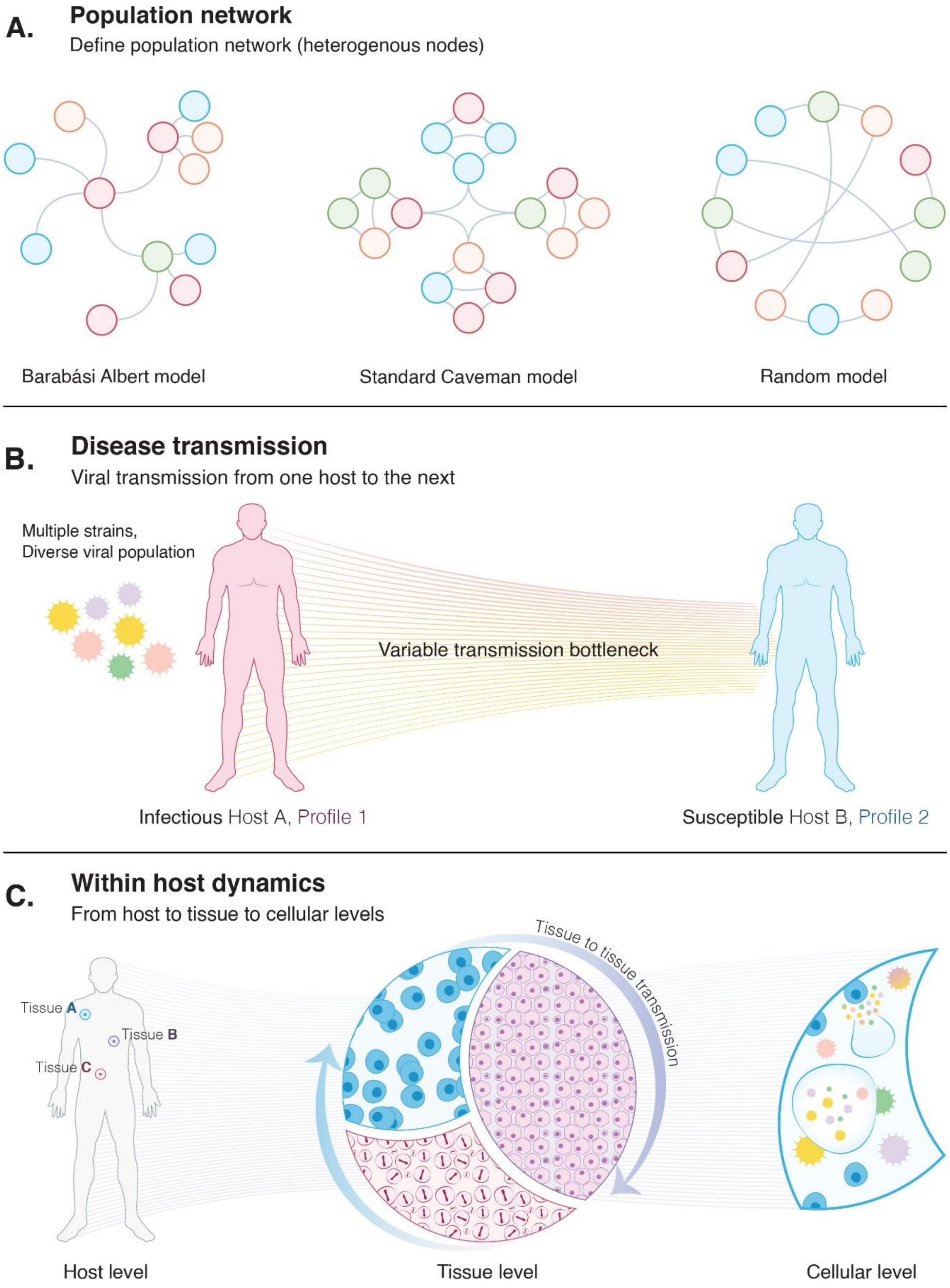
A.
Population network
Define population network (heterogenous nodes)
Barabási Albert model
Standard Caveman model
Random model
B.
Disease transmission
Viral transmission from one host to the next
Multiple strains,
Diverse viral population
Variable transmission bottleneck
Infectious Host A, Profile 1
Susceptible Host B, Profile 2
C.
Within host dynamics
From host to tissue to cellular levels
Tissue A
Tissue B
Tissue C
Tissue to tissue transmission
Host level
Tissue level
Cellular level

**Figure S - 1:** Depiction of the different levels of configuration that Apollo provides to enable the complete capture of the forces that govern an epidemic. (A) Generation of the susceptible populations' contact network. The colored nodes on the networks represent the different host profiles. Apollo provides a wide array of network models enabling the use of heterogeneous populations. (B) Configuration of transmission mechanics between hosts, including complete control over the disease transmission bottleneck. (C) Overview of the within-host dynamics from tissue to cell levels that govern evolutionary processes. Going beyond the host level Apollo enables the configuration of multiple tissues per host including control over tissue dynamics such as cell concentration and viral transmission between tissues. Furthermore, it accounts for the cellular level environment and the genomic variations and phenotypic responses of the viral particles.

When accounting for the within host evolutionary processes Apollo comes equipped with a series of parameters. At the genomic level, we have provided segmentation of the genome. These segments can then be subjected to different mutation rates and selection pressures. It also factors mechanisms such as proofreading as well as recombination between viral particles and how these mechanisms change based on the mutations that arise during the epidemic.

It should be noted that the true power of the simulator comes in its ability to generate each individual viral particle complete with its sequences. We are able to execute at this resolution due to the use of our large-scale parallel processing architecture and our partial random access file structure developed in CATE[1].

#### 1.3. Proven high performance architecture.

Apollo leverages CATE's proven large-scale parallel processing and scalable architecture to achieve the degree of intricacy promised[1,15]. This foundation enables it to achieve unprecedented speeds and hardware efficiency. Furthermore, we have taken steps to extend our architecture's parallel processing capabilities by incorporating a multi-GPU (Graphical Processing Unit) framework that surpasses our previous single GPU architecture[16]. Apollo benefits from CATE's CPU (Central Processing Unit), GPU, and SSD (Solid State Drive) parallel processing capabilities including its segmented file structure. Similar to CATE it is a C/C++ CUDA-based program designed for the Linux kernel. Simulations are parameterized via JSON scripting and software interactions are conducted via the command line.

#### 1.4. Why use Apollo?

At present, there already exists a myriad of evolutionary toolkits[7,9,17–20]. They vary from scripting-based solutions enabling the simulation of a wide range of evolutionary scenarios with granularity in configuring individual mechanics to simulators cater made to a specific organism and use case. Apollo is a hybrid solution that takes the best parts of these worlds. It provides a flexible solution that can be customized using JSON scripting to configure a wide range of evolutionary scenarios while being a cater made stochastic simulator for dynamic haploid viral genomes.

First, Apollo is specifically designed for haploid viral genomes, it is not limited by the genome length or the specific viral organism under study. It provides the user with the flexibility to segment the genome under factors such as mutation and recombination hotspots. It can account for genetic variation via factors such as fitness, survivability, and proofreading, even affecting recombination rates of individual hotspots.

Second, to the best of our knowledge, Apollo is the first simulator that comes with native integrations for the five hierarchical levels of an epidemic. Namely, these hierarchies are defined as, network, host, tissue, and cellular and viral genome. Apollo allows an unparalleled level of granularity in terms of pandemic configuration. The user is able to easily design the contact networks being simulated, structure the transmission of viral particles between hosts, and control the tissues that are available for viral shedding and, tissues that will receive the viral load. At host level, the user can configure a multitude of different host profiles. This enables Apollo to conduct simulations with heterogeneous host populations, moving away from the traditional homogenous node design adopted by most simulators[9]. Host profiles allow individual hosts to have different behavioral patterns. For instance, simulate patients who are affected by sampling. Such as where they reduce their infectious probability upon being sampled and those that do not and remain infectious[21]. The latter is reminiscent of Lost-To-Follow-Up (LTFU) individuals as observed in for instance HIV AIDS[20,22,23] including other disease scenarios. Another instance of similar host behavior is with the recent COVID-19 pandemic. Where populations resorted to quarantine upon diagnosis and individuals who still maintained social contact in spite of being diagnosed with the virus[4,24,25]. At the within host level the user is able to structure an unlimited number of tissues. Tissues can be assigned various roles such as tissues whose viral load will contribute to a host being infectious. At the cell level, the number of viral particles that inoculate a cell can be defined, these factors are highly important when configuring recombination mechanisms where the exchange of genetic material occurs between viral particles infecting the same host cell[26,27]. Finally, at the genomic level, Apollo allows control of the mutations of the viral genomes, and behavior of different strains and incorporates proofreading mechanisms. The user is also able to introduce external response mechanisms such as host immune responses, vaccination, and drug therapy.

Third, as mentioned before we adapt our proven large scale parallelization algorithm developed for CATE into Apollo[1]. It is this architecture that enables us to create a simulator that can feasibly conduct evolutionary simulations from network to individual viral genome level resolution with high levels of fidelity.

Fourth, we have provided the user with a scripting feature to configure the properties of the simulations they wish to design. Our scripting style uses the standard JSON format, and attention has been paid to ensure that the keywords and variable syntaxes are straightforward and self-explanatory. This removes the need for learning and writing complex code, which usually limits the use of certain simulation software. The scripting approach was implemented to accommodate the wide variety of flexibility that Apollo provides as well as make the implementation user-friendly.

Finally, Apollo's processing capabilities and granularity provide the ability to test a wide range of hypotheses at an accelerated pace. Most statistical solutions developed for epidemiological modeling are complex and in certain instances might even be difficult to formulate and execute. Apollo allows the user with the toolset to easily configure simulations that capture the intended dynamics of the hypothesis being tested, without the need for complex statistical intervention. Users can use Apollo to either build their null distributions across large numbers of simulations to test their hypothesis or conduct simulations to observe the behavior of their experimental models in an epidemiological setting.

### SECTION 2. MODULES

In this section, we will introduce the design language that allows the orchestration of the five epidemiological hierarchies. Apollo was designed on the foundation of three modules. They are the Network, Host and Genomic modules. Collectively they help capture the different hierarchical levels from the contact network to the genome. Each module is responsible for capturing an epidemic component and they work together to provide the user with a seamless and resource efficient simulation.

We will begin with the network module. It is tasked with generating the contact networks that capture the physical interactions responsible for the spread of the virus. Apollo's network module is equipped with a series of proven networks from graph theory including Random networks, Barabási Albert models, and more[9,28].

The hosts in the population are the units of infection in an epidemic. Their within host environment is what governs the spread of disease within the individual and subsequently the population[29,30]. Apollo attempts to provide a complete capture of the within host environment by providing the user with the capability to configure hosts complete with tissue and cell dynamics. These dynamics include viral transmission between tissues and account for external forces such as immune responses and drug resistance[31,32]. Additionally, Apollo enables users to configure multiple host profiles with capabilities to allow diversity between hosts[3].

The genome of the virus directs its phenotypic behavior[33]. A strain's phenotypic response is responsible for its virulence and success as an infectious agent[34–36]. This faction is controlled by the Genomic module. The Genomic module's infrastructure is designed to capture mechanisms such as mutation, proof reading, and recombination which create diversity in a viral population. Furthermore, it then captures the resultant phenotypic responses that will lead into selection pressures which govern the progression of the epidemic in a population.

In this section, we will take a detailed look into each of these modules with the goal of obtaining an understanding how the respective evolutionary principles and mathematical models of each module was implemented. Any deviations implemented by Apollo to the standard assumptions or principles will be discussed and validated.

#### 2.1. Network module

Apollo has five main types of Networks, equipped to provide the user with a series of options to generate validated contact networks that are adaptable to numerous diseases. The provided network types are as follows:

A. Random model

The random model is a modification to the Erdős-Rényi model. We believe that similar to the Erdős-Rényi model discussed below our random model will be ideal for simulating transmission dynamics with unknown population structures such as in the spread of HIV (Human Immunodeficiency Virus) and HCV (Hepatitis C Virus) among People Who Inject Drugs (PWIDs)[37–40].

B. Erdős-Rényi random model

The Erdős-Rényi model is commonly used in studying epidemic spreads with unknown population structures, typically lacking preferential attachment[37]. It has been used to model contact networks of individuals that represent Lost To Follow Up (LTFU) and PWID in diseases such as HCV and HIV[37,41–43].

C. Barabási Albert model

The BA model is ideal for representing contact networks that house social structures that represent preferential attachment such as hubs. This is ideal for simulating contact networks that govern the transmission of sexual diseases such as HIV due to phenomena such as homophily among infected individuals[37,44].

D. Standard Caveman model

Caveman models are suitable for simulating diseases spread via contacts caused by closely interconnected clusters or communities (referred to as caves or cliques), and how these clusters will in turn spread the disease across other such clusters in the population. Such examples lie in diseases such as COVID-19 where increased spread was seen initially among individuals of close contact in a community[28,45].

E. Dynamic Caveman model

The Dynamic Caveman is a modification of the above standard caveman model. It allows for relaxation of the parameters allowing a more real-world representation of contact networks with densely connected communities.

The Network function can be defined as $Network(network\_Type, params)$ where the $network\_Type$ will take the form of one of the network types listed above and $params$ will be the parameters that define the generated network.

In the following subsections, we will look at the mathematical principles that govern these models and their viability for certain epidemic cases. We will also discuss the modifications that have been implemented onto these models by Apollo to provide an additional layer of flexibility where relevant.

#### 2.1.1. Random model

This is the simplest network model available in Apollo. It is based on the Erdős-Rényi model of a random graph[43]. In contrast, we begin with a fixed number of nodes and iterate over each node randomly assigning it to another node ensuring that all nodes in the network are connected to at least one other node. The process of the random model generation is as follows:

1. Start with $N$ number of isolated nodes.
2. Iterate over each node $N$ and select another node ($U$) at random using a uniform distribution. The probability of two nodes being paired is $1/N$ and the number of links in the network is always $N - 1$.

$$U \sim Uniform(0,1,2..,N-1) \tag{2.1}$$

$$P(U = k) = \frac{1}{N} \text{ for } k \in \{0,1,2,..,N-1\} \tag{2.2}$$

Therefore, this graph model (**Figure S - 2**) is represented as $G(N, N-1, 1/N)$, where $N$ is the number of nodes, $N - 1$ number of links with a $1/N$ probability of two nodes being linked.

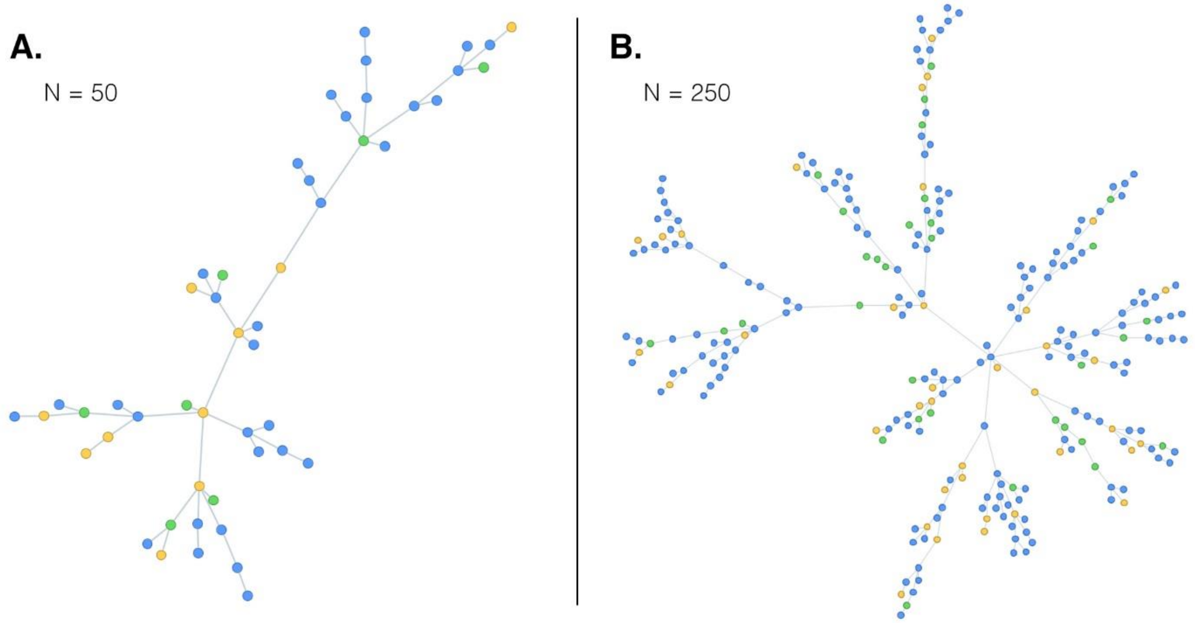

**Figure S - 2.** Two random graphs generated by Apollo. (A) Has a total of 50 nodes interconnected with each other and (B) forms a network with 250 nodes. As observed all nodes are connected to form a complete singular contact network. The nodes are colored by the node profile they belong to. There are three node profiles in the above networks (Blue, yellow, and green).

#### 2.1.2. Erdős-Rényi random model

Apollo implements the $G(N, p)$ variant of the Erdős-Rényi random model, where $N$ is the number of nodes and $p$ is the probability that two nodes would be linked[43]. The process of generating a contact network model according to the Erdős-Rényi model is as follows:

1. Start with $N$ number of isolated nodes.
2. Iterate over each node pair ($N(N-1)/2$ node pairs present). Then using a Bernoulli distribution of probability $p$ for 1 and of $(1-p)$ for 0 we determine whether the two nodes are linked or unlinked. If 1 then the nodes are linked and 0, they are unlinked.

$$X \sim Bernoulli(p) \text{ if } p \in [0,1] \tag{2.3}$$

$$P(X = k) = p^k(1-p)^{1-k} \text{ if } k \in \{0,1\} \tag{2.4}$$

It should be noted that there is a possibility that all the nodes in the network will not be interconnected with each other. In contrast, it can also be used to create complete graphs by setting $p$ to 1 (**Figure S - 3A**).

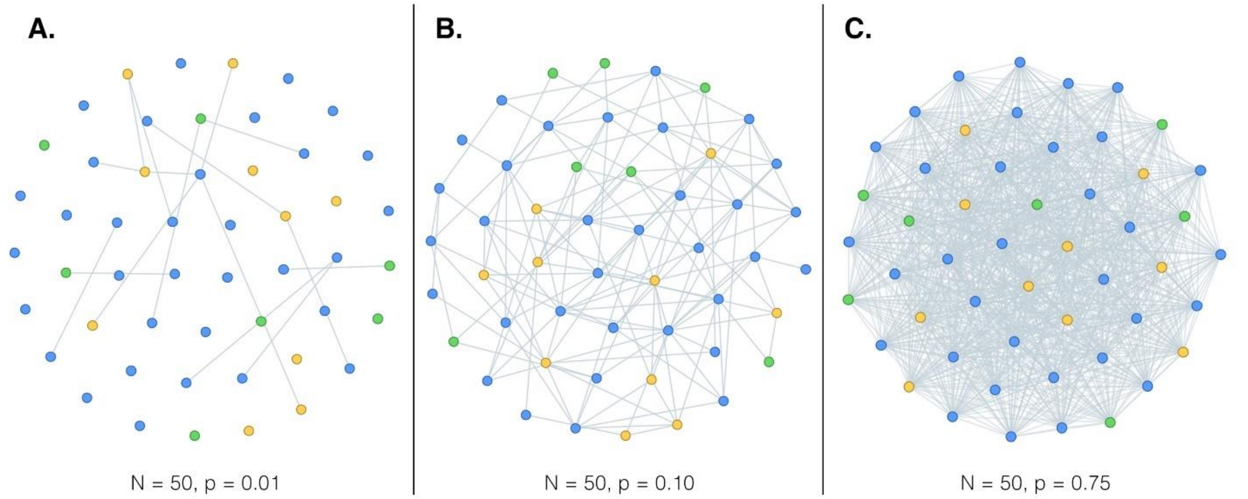

**Figure S - 3.** Contact network graphs of 50 nodes generated under the Erdős-Rényi model with varying probability of node pairs being connected ($\boldsymbol{p}$). (A) Uses a $\boldsymbol{p}$ value of 0.01 and as observed a complete interconnected graph is not formed. In (B) and (C) all nodes are interconnected with each other, but the number of links progressively increase with the increasing probability ($\boldsymbol{p}$). The nodes are coloured by the node profile they belong to. There are three node profiles in the above networks (Blue, yellow and green).

#### 2.1.3. Barabási Albert model

The Barabási Albert model enables the creation of scale-free networks governed by a power law that enables preferential attachment. The resultant networks have the presence of hubs (nodes with higher degrees) and follow a first come first serve principle, where the nodes that first entered the network tend to accumulate connections.

The Barabási Albert networks $G(N, m)$, generated by Apollo contain a pre-specified number of Nodes ($N$). Similar to the previous networks the number of links ($m$) assigned to a newly incoming node and can be determined by using three modes. These modes include a fixed notation ($M$), always having a fixed number of links, or we can determine the number of links using a Poisson or Negative Binomial distribution. Links to the newly incoming node are added one at a time and self-loops are not allowed. Apollo carries out the generation of the model via time steps ($t$) till the user-specified $N$ number of nodes are present in the network; the graph can be presented as $G_m^{(t)}$. The process of network generation is as follows:

1. Determine the $N$ number of nodes that will be present in the network.
2. Determine the distribution used to determine the $m$ number for each newly incoming node.

$$m \sim \begin{cases} Poisson(\lambda) \\ NegativeBinomial(r, p) \\ M \end{cases} \qquad (2.5)$$
$$\text{where } M \in \{1,2,3, \dots, \infty\}$$

3. Start with an empty graph, where only in the first instance it is assumed by default $m = 1$. Therefore, the graph at $t = 0$ is $G_m^{(0)}$.
4. From here onwards given $G_m^{(t-1)}$ we can generate $G_m^{(t)}$. $m$ is determined by the parametrization in (2.5). The incoming node $V_t$ is added and it's linking node $V_i$ is determined using the preferential attachment probability $\prod(k)$ where $k$ is the degree of a given node. $\prod(k)$ is defined as follows:

$$\prod(k_i) = \frac{k_i}{\sum_j k_j} \qquad (2.6)$$

where $i$ is the node being considered and the $j$ is the total number of nodes present in $G_m^{(t)}$.

The total probabilities of attachment of all nodes at time $t$ will sum to one, this includes the new node $V_t$ which is assumed to have a probability of attachment of ${}^{1}/_{\sum_j k_j}$ at first.

$$\sum_{i=1}^{j}\left(\frac{k_i}{\sum_j k_j}\right) = 1 \tag{2.7}$$

Then using a discrete probability distribution, we determine the node to which the new node $V_t$ is being linked to.

5. For $m > 1$ the $G_m^{(t)}$ is built by adding the $m$ links one at a time repeating the above step 4 for each new link. In all instances, self-loops are omitted.
6. Steps 4 and 5 are repeated till $G_m^{(t)}$ has $N$ number of nodes present (**Figure S - 4**).

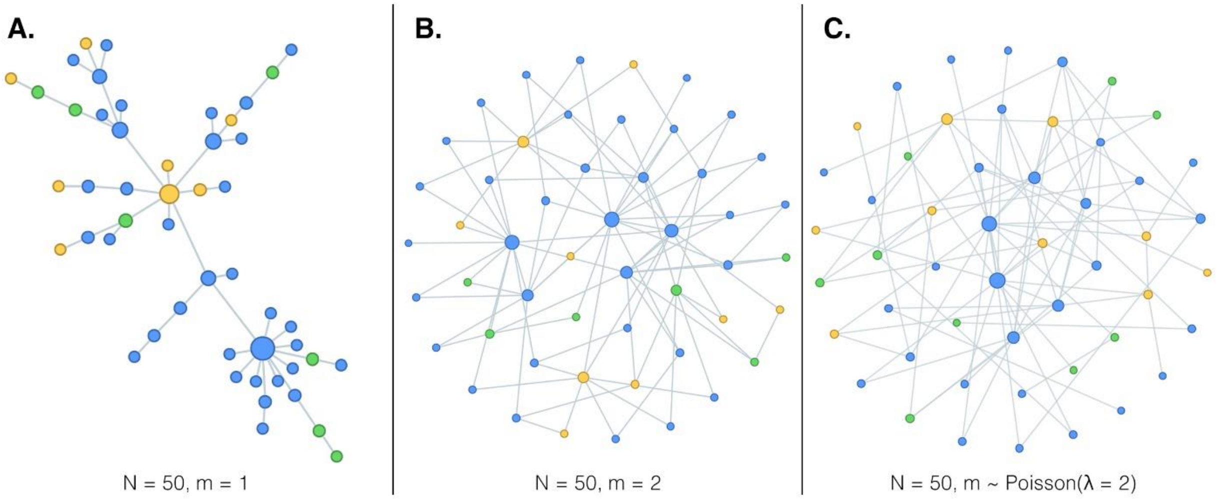

**Figure S - 4:** Contact network graphs of 50 nodes generated under the Barabási Albert model. The size of the nodes is ordered by their degree, nodes with larger degrees are depicted as larger in size. (A) The number of links ($\boldsymbol{m}$) is fixed at 1 forming the simplest form of the Barabási Albert model, the formation of Hubs can be seen. (B) Here too $\boldsymbol{m}$ is fixed but it is set to 2, and in (C) $\boldsymbol{m}$ for each newly incoming node is determined via a Poisson distribution where $\boldsymbol{\lambda = 2}$. The nodes are coloured by the node profile they belong to. There are three node profiles in the above networks (Blue, yellow, and green).

##### 2.1.4. Standard Caveman model

This is Apollo's implementation of the Connected Caveman model. In contrast to the previous models connected caveman models are used to capture networks with interconnected systems. These systems take the form of interconnected tightly knit node clusters, referred to as cliques or caves. Tightly knit clusters of nodes form caves. These caves are then interconnected by nodes in the caves (**Figure S - 5**). Such connected caveman models are defined as $G(X, C)$ where $X$ is the number of nodes present in each cave and $C$ is the number of caves. Therefore, the total number of nodes ($N$) present in the network is $N = X \times C$. The process of generation of the standard caveman model is as follows:

1. We start with $C$ number of caves and each cave has $X$ nodes.
   Caves are labeled by their number $c$ where $c \in \{0,1,2 \dots, C-1\}$.
   Nodes are labeled as $n\_c$, where $n$ is the number of the node within its cave (where $n \in \{0,1,2, \dots, X-1\}$) and $c$ is the cave, it belongs to.
2. First, we form the intra-cave connections. Each node is connected to every other node within its cave without self-loops; therefore, each node has $(X-1)$ edges. This step is repeated for each node in every cave.
3. Next the connections between caves in established. A node is selected at random from each cave to become the neighbour forming node. The neighbour forming node will be $k_{n\_c}$.

$$n \sim Uniform(0,1,2, \dots, X-1) \tag{2.8}$$

4. To form a complete graph, each cave's $k_{n\_c}$ will have to form connections with its adjacent cave. We can intuitively understand that this cave will be the immediate cave after. Therefore, node $k_{n\_c}$ will connect with nodes $k_{n\_c+1}$.
   This holds true except for the last cave ($c = C-1$). For the last cave c where ($c = C-1$) the node $k_{n\_c+1}$ will actually be $k_{n\_0}$. We can represent this as a function as follows:

$$f(c, n) = \begin{cases} k_{n\_c+1}, if\ 0 \le c < C-1 \\ k_{n\_0}, ifc = C-1 \end{cases} \tag{2.9}$$

5. Next an intra node connection from the node ($k_{n\_c}$) will be removed at random before establishing the connection to the neighboring node. This rerouting preserves the total edges in the connected caveman graph.

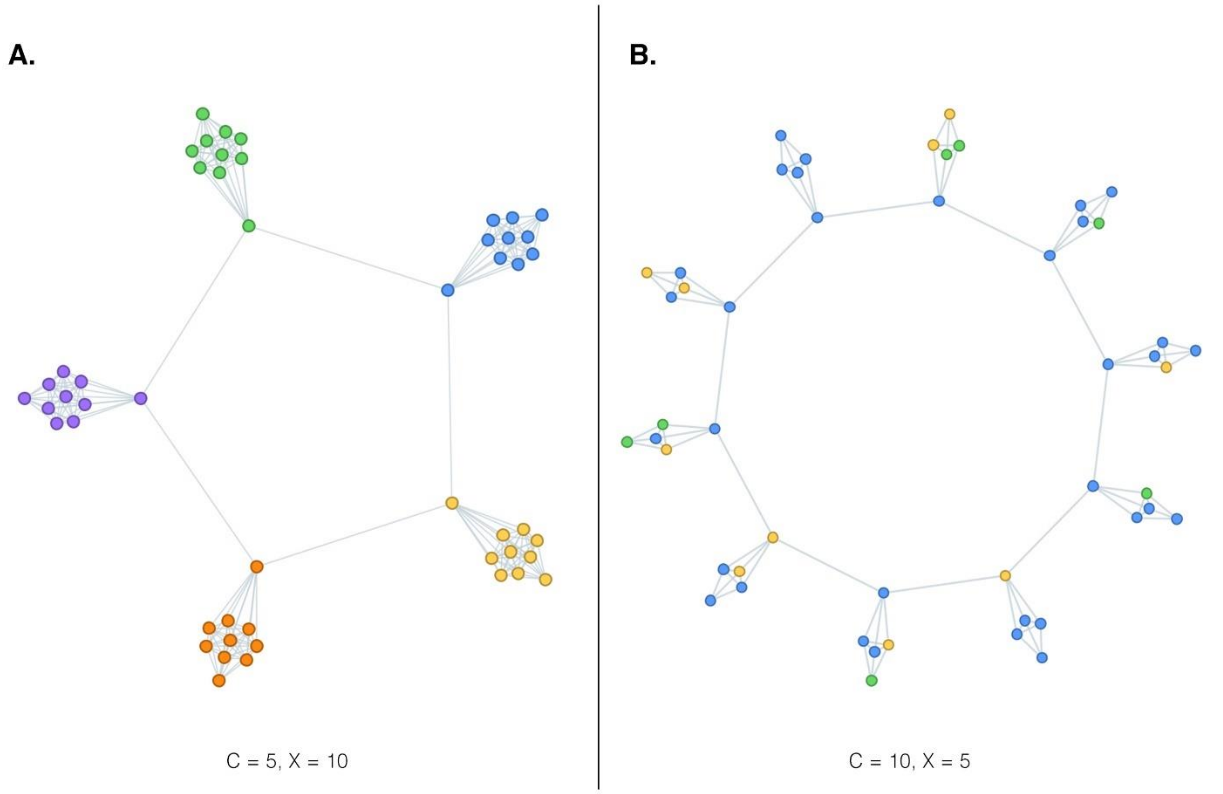

**Figure S - 5.** Contact caveman graphs generated by Apollo with 50 nodes. (A) The nodes are distributed between five caves with ten nodes each. The nodes are colored by the cave they belong to. (B) The nodes are distributed among ten caves with five nodes in each cave. The nodes are colored by the node profile they belong to. There are three node profiles in the network (B) (Blue, yellow, and green).

#### 2.1.5. Dynamic Caveman model

The Dynamic Caveman model is an extension of the standard connected caveman model (2.1.4). It is similar to connected caveman model in terms of the presence of the caves, the complete interconnectedness of the nodes in the caves, and the formation of a complete network by neighboring caves being connected. Apollo's implementation allows us to go beyond the standard connected caveman model by adding a few layers of flexibility. The goal of the dynamic caveman model is to provide the user with the flexibility to better capture real-world dynamics by removing some of the restrictions that pertain to the connected caveman model.

Similar to the connected caveman model, the dynamic caveman model uses a fixed number of caves ($C$). However, the number of nodes per cave can be variable. The number of nodes in a cave can be determined using a Poisson distribution or a negative binomial distribution.

The number of nodes in a cave is defined as $C_i$ where $i$ is the cave number and $i \in \{0,1,2 \ldots C-1\}$.

If the user has selected the Poisson distribution for node per cave generation, then the value of $C_i$ can be denoted as follows:

$$C_i \sim Poisson(\lambda) \tag{2.10}$$

And if the negative binomial distribution is selected then:

$$C_i \sim NegativeBinomial(r, p) \tag{2.11}$$

We can define this as a concise function:

$$f(C_i, D, params) = \begin{cases} Poisson(C_i; \lambda) \text{ if } D = \text{"}Poisson\text{"} \\ NegativeBinomial(C_i; r, p) \text{ if } D = Negative\ Binomial \end{cases} \tag{2.12}$$

Where $params$ is a tuple storing the parameters of the distribution selected.

Therefore, the total number of nodes ($N$) in the network will be:

$$N = \sum_{i=0}^{C-1} C_i \tag{2.13}$$

Apollo's dynamic caveman models also enable the user to determine the number of nodes in a caveman that can take part in forming connections with their neighboring caves. This is in contrast to the standard caveman model where only one node per cave took part in the forming of connections with the neighbor caves. The number of neighboring nodes ($k$) from the total number of nodes in a cave is determined using a percentage α where $\alpha \in [0,1]$.

A major contrast between the dynamic caveman model and the standard caveman model is the implementation of global neighbor nodes ($g$). These nodes can form connections with not only neighboring caves (Equation ((2.9)) but also with distant caves, hence global connections. The number of global nodes is a subset of the total number of neighboring nodes ($k$) in a cave that is determined using a percentage $\beta$ where $\beta \in [0,1]$.

We can define the dynamic caveman model as $G(C, D, params, \alpha, \beta)$ (**Figure S - 6**). We are now ready to look at the process of generation for the dynamic caveman model:

1. Starting with each cave we determine the number of nodes in each cave ($C_i$) using the function $f(C_i, D, params)$ from Equation (2.12).
2. Then intra cave connections are formed. Each node is connected to every other node within its cave without self-loops; therefore, each node has ($C_i - 1$) edges. This step is repeated for each node in every cave.
3. To form connections with the nodes from caves forward and backward in space the number of neighboring nodes is determined for each cave using a binomial distribution, a portion of these nodes will also take part in the formation of the global nodes.

$$k_i \sim Binomial(C_i, \alpha) \tag{2.14}$$

4. We then select the different $k_{n\_i}$ neighboring nodes where $n$ is the node within the cave where $n \in \{1,2,3 \dots, C_i\}$.

$$f(k_i, i) = \{x_{0\_i}, x_{1\_i}, \dots x_{(k_i-1)\_i}\} \text{ where } f(k_i, i) \mapsto X_i \tag{2.15}$$

Where $X_i$ is returned with the set of numbers selected uniformly within a range of integers from 0 to $C_i - 1$.

$$x_{y\_i} \sim Uniform(0,1,2, \dots, C_i - 1) \tag{2.16}$$

5. Similar to Equation (2.9) we form connections with the adjacent caves' neighboring node. From the collection of neighboring nodes ($X_i$) present per cave $i$ one is selected uniformly at random. No rerouting of connections is performed.
6. Then we have to determine the number of nodes available to form global connections. This too is determined using a binomial distribution.

$$g_i \sim Binomial(k_i, \beta) \tag{2.17}$$

7. Similar to Equation (2.15) we then determine which nodes will form global connections. But we will select the global nodes from the collection of neighboring nodes ($X_i$).
8. Then Apollos iterates over each cave and attempts to connect the global nodes of that cave to a randomly selected cave and its global nodes. No rerouting of connections is performed.

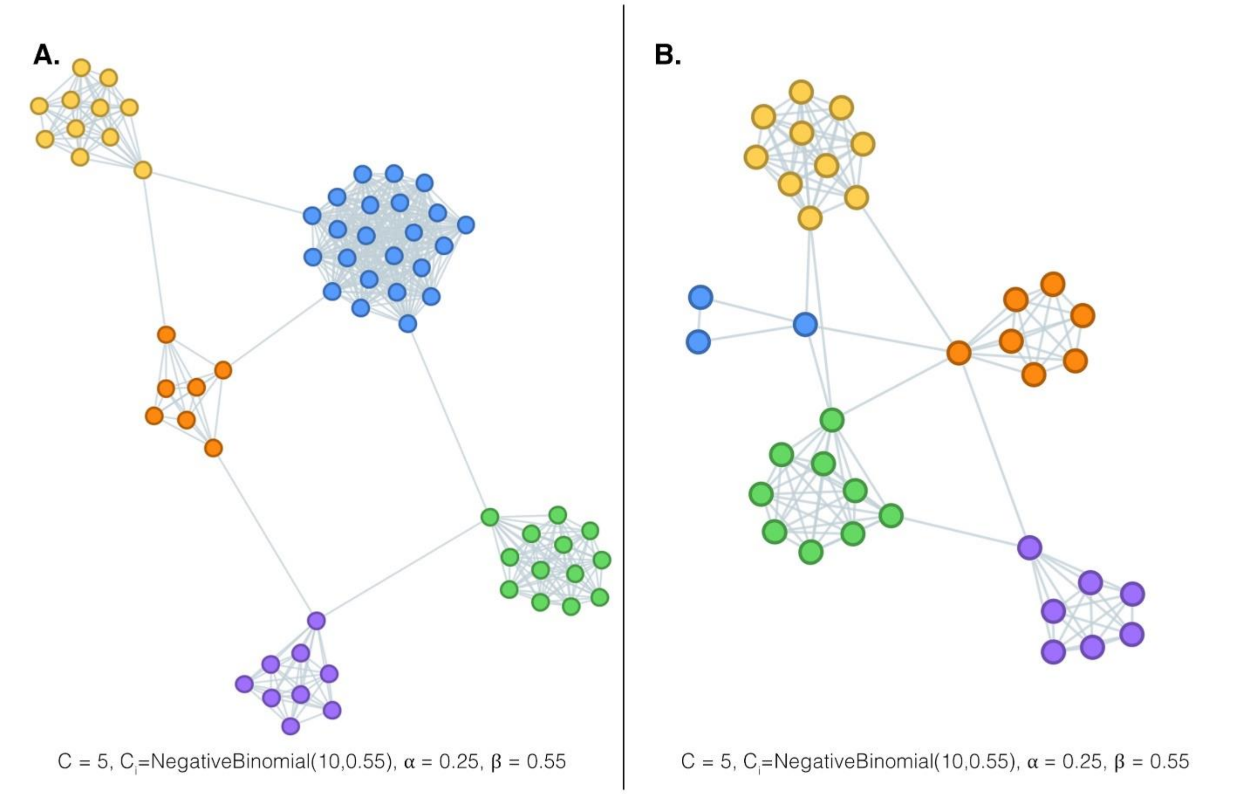

**Figure S - 6.** Dynamic caveman graphs generated by Apollos. As observed even though both graphs (A) and (B) have the same parameters, separate simulations generate different graphs. This shows the dynamic nature of the graph model. Additionally, the formation of global connections among caves that transcend beyond their immediate neighbors can also be observed.

#### 2.2. Host module

The properties of the hosts in the network in relation to an epidemic are managed by the host module. The susceptible hosts, referred to as nodes in the network are the units of infection.

To provide a complete capture of the host dynamics, Apollo moves across the hierarchical levels of an epidemic encompassing pathogen transmission between hosts and into the intricate within host dynamics at tissue and cell levels. Furthermore, it not only simulates the phenotypic responses of viral genomes on the host but also the host's responses to an infection.

Beginning at the population level the host module accounts for host-to-host transmission. The module controls the rate of viral transmission between infected and susceptible hosts. It is also responsible for accounting for the sampling of infected hosts and their behavioral responses to actions such as sampling. At the population level, the host module controls the distribution of different host profiles across the contact network.

Moving into the within host dynamics the host module accounts for both tissue and cell levels before focusing on the pathogen genome. The host module initiates the within host configuration by the structuring of the tissue structures, their roles, interactions, and affinity to the pathogen during infection in addition to the transmission of pathogens between the configured tissues.

At the pathogen level, the host module begins by parametrization of viral factors that govern the duration of infection. The module begins by configuring viral behavioral factors such as its generation time and rate of progeny generation including the potential time of the disease persistence in each host.

In this section, we will investigate how these parameters are structured inside the host module, including how they connect from the population level to within host dynamics.

#### 2.2.1. Determining the generations spent for the duration of infection

Apollo begins by determining the generation time of the viral pathogen. This is the time taken from viral cell attachment to the host cell to the release of newly assembled viral progeny. The generation time ($G$) is determined by using a gamma distribution of shape $\alpha_1$, and scale $\beta_1$.

$$G \sim Gamma(\alpha_1, \beta_1) \tag{2.18}$$

Apollo then determines the duration of infection, $D_h$ $where\ h\ \in \{0,1,...,N-1\}$) of each individual host, $h$ from the total susceptible population $N$ in the contact network. $I_h$ is determined using a gamma distribution of shape $\alpha_2$, and scale $\beta_2$.

$$D_h \sim Gamma(\alpha_2, \beta_2) \tag{2.19}$$

Therefore, the number of generations of each host $g_h$ spent in the event of infection is determined by the division of Equation (2.19) by Equation (2.18).

$$g_h = \frac{D_h}{G} \text{ where } g_h \in Z \tag{2.20}$$

Apollo keeps track of the epidemic using generations. Therefore, it requires the calculations of $g_h$. $Z$ represents all positive integers ($Z\ \in \{0,1,2, \dots \infty\}$).

#### 2.2.2. Rate of progeny generation

The rate of progeny generation ($R$) can be determined by using either a negative binomial, gamma distribution, or a Poisson distribution.

$$R = \begin{cases} NegativeBinomial(r_1, p_1) \\ Gamma(\alpha_3, \beta_3) \\ Poisson(\mu_1) \end{cases} \text{ where } R \in Z \tag{2.21}$$

$R$ captures the number of progenies that are generated on average by a parent viral particle. The number of progenies generated can change with mutations. During an event of replication $R$ is determined by a draw of a random number from the selected distribution type.

#### 2.2.3. Tissue configuration

Each host will have a configuration of tissues that the virus interacts with. The host module enables the configuration of these tissue structures, complete with their labeling, role assignments, and cross tissue viral transmission. In this subsection let us take a closer look into these factors.

Apollo enables the user to configure an unlimited number of tissue types ($T$) and to help the user keep track of them the user can assign names or labels to each individual tissue type ($t_i$ $where\ i\ \in \{0,1,..,T-1\}$). Secondly, each tissue type can be part of five roles. Based on the tissue role, the resultant effect will be executed by the cumulative number of viral particles occupying the tissues belonging to that specified role.

The user can structure each tissue's cellular environment based on each host profile. We will look into this in the following subsections, but for now, we will take a quick look into how the host module factors the inter-tissue viral transmission.

##### 2.2.3.1. Tissue roles

Tissues can have five major roles. Namely viral entry ($\pi_1$) tissues and exit tissues ($\pi_2$), respectively the tissues from which viral particles enter a host and exit a host during host-to-host infection. These are followed by two load tissues whose cumulative viral loads will trigger specific mechanisms. Namely the infectious ($\pi_3$) and terminal ($\pi_4$) tissues whose viral loads will determine a host becoming infectious or reaching mortality. Finally, we have the sampling tissues ($\pi_5$), from whom sequence samples will be obtained during a sampling event.

$$Tissues = \{0,1,\ldots,T-1\} \tag{2.22}$$

$$\text{where } \pi_1\pi_2,\pi_3,\pi_4,\pi_5 \subseteq Tissues \tag{2.23}$$

##### 2.2.3.2. Viral transmission between tissues

The viral transmission from a tissue $x$ to tissue $y$ defined as $M_{xy}$ where $x, y \in \{0,1, \dots (T-1)\}$ and $x \neq y$. The number of viral particles migrating is determined using a binomial distribution.

$$M_{xy} \sim Binomial(n_{xy}, p_{xy}) \tag{2.24}$$

This can take the form of a matrix M of size $T \times T$, since M can be quite sparse in practicality, we only store the defined tissues to tissue transition values.

$$M = \begin{bmatrix} 0 & \cdots & M_{0(T-1)} \\ \vdots & \ddots & \vdots \\ M_{(T-1)0} & \cdots & 0 \end{bmatrix} where\ M_{xx} = 0 \tag{2.25}$$

#### 2.2.4. Sampling of hosts

The sampling of hosts is an optional mechanic. Sampling can also be configured to have effects on host infectivity dependent on the profile a particular host belongs to. A factor we will look at later when discussing host profiles. Sampling is configured to sample a certain number of hosts per generation and obtain a certain number of samples from each host. The mechanisms provide capabilities to sample hosts multiple times throughout consecutive generations or sample the same host multiple times during a single sampling event.

The number of hosts ($H$) being sampled per generation is determined by a Binomial distribution and the number of sampling events ($H_h$) conducted on each sampled host ($h$) is determined by either a fixed value parameter ($\theta_1$) or another binomial distribution.

Therefore, we determine $H$ as follows:

$$H \sim Binomial(n_2, p_2) \tag{2.26}$$

and the number of sequencing events for each host:

$$H_h = \begin{cases} Binomial(n_3, p_3) \\ \theta_1 \text{ where } \theta_1 \in \{1, \dots, \infty\} \end{cases} \text{where } h \in \{0,1, \dots, (H-1)\} \tag{2.27}$$

#### 2.2.5. Host profiles

The host profiles allow a more granular configuration of the hosts, allowing the user to create a truly heterogeneous population. In the host profile, the module configures factors such as the viral load limits for particular roles such as infectiousness, mortality, transmission bottlenecks, and sampling effects. It goes beyond this to the tissue's cellular level allowing the user to configure the number of cells available for viral attachment for each tissue and control the affinity of each tissue's cell types. The host profile also provides the host module with the necessary parameters to control the eternal forces that may affect the viral load in a tissue, such as immune response or drug therapy. The user specifies the total number of host profiles that can be present ($J$ where $J \in 1, \infty$) and each host ($h$) can have a particular profile ($h_j$ where $j \in \{0,1,2,..,J-1\}$)

##### 2.2.5.1. Load tissues

The load tissues are infectious ($\pi_3$) and terminal ($\pi_4$). They trigger a response once the cumulative viral load within these tissues exceeds a user specified limit. Therefore, the response function ($R$) can be defined as follows in the form of a Heaviside step function:

$$R(\mathrm{\pi_i}, \mathrm{\omega_{i\_j}}) = \begin{cases} 1, & \text{if} \sum_{t \in \pi_i} t \geq \mathrm{\omega_{i\_j}} \\ 0, & \text{Otherwise} \end{cases} \tag{2.28}$$

Where $\pi_i$ is the load tissue type set being processed ($\pi_3$ $or$ $\pi_4$), $t$ are the elements that refer to the viral load count of each tissue in the set and $\mathrm{\omega_{i\_j}}$ is the user specified limit for specific load tissue as $i \in \{3,4\}$ based on each profile $j$.

$\omega_{i\_j}$ can be defined either by a binomial distribution or a fixed value ($\theta_{i\_j}$).

$$\omega_{i\_j} = \begin{cases} Binomial(n_{i\_j}, p_{i\_j}) \\ \theta_{i\_j} \text{ where } \theta_{i\_j} \in \{0, \dots, \infty\} \end{cases} \tag{2.29}$$

#### 2.2.5.2. Tissue structures

Apollo allows the user to configure the tissue structures at a profile level so that each host profile can have a specific tissue structure and even replication phases. The tissue structure encompasses a specific tissue type ($t_i$), has a limited number of cells for infection, and the affinity of a specific tissue's cells to the virus. These are configurable at the host profile level. Meaning that each host profile can have different values for these factors.

Looking at the number of cells available for infection per tissue ($\lambda_{i\_j}$), they can either be limited or unlimited. Unlimited means the viral particles present in the tissue will have a limitless number of cells to attach to for viral replication. Else, if limited, the number of cells present can be defined by a binomial distribution.

$$\lambda_{i\_j} = \begin{cases} Binomial\left(n\_limit_{i\_j}, p\_limit_{i\_j}\right) \\ \infty \end{cases} \tag{2.30}$$

The affinity of viral particles to the cells of a tissue can ($a_{i\_j}$) be defined using either a gamma or binomial distribution.

$$a_{i\_j} = \begin{cases} Binomial\left(n\_affinity_{i\_j}, p\_affinity_{i\_j}\right) \\ \mathrm{Gamma}(\alpha\_\mathrm{affinity}_{i\_j}, \beta\_\mathrm{affinity}_{i\_j}) \end{cases} \tag{2.31}$$

#### 2.2.5.3. Host infection rate

The number of hosts being infected by a host defined as $S_h$ belonging to host profile $j$ can be determined by either a binomial distribution or a fixed value.

$$S_h = \begin{cases} Binomial\left(n\_infection_j, p\_infection_j\right) \\ \theta\_infection_j \text{ where } \theta\_infection_j \in \{0, \dots, \infty\} \end{cases} \tag{2.32}$$

##### 2.2.5.4. Replication phases

The replication phases designed to capture a degree of external factors that may influence the within host viral population can be one of three types namely, Neutral, Stationary, and Depreciation. They mainly help control the next generation's population. The number of phases ($X \in 1,..,\infty$) can be unlimited, and they can be any combination of the three phase types. The phase type each generation belongs to is determined by the time ratio of each phase $X_x$ where $x \in \{0,1,..,X-1\}$.

If we define ($P$) which is the set of phases for each generation in a host where $P = \left\{\rho_{h_i}^{0}, \rho_{h_i}^{1}, \rho_{h_i}^{2}..., \rho_{h_i}^{g_{h-1}}\right\}$ where each ($\rho_{h_i}^{x}$ for $x = 0,1,2,\ldots, g_h - 1$ where $x$ is the generation) can be one of the three phases. The function for determining the next generations viral load population can be defined as follows:

$$f\left(G_h, V_{h_i}^{G_h}, i, h_j\right) = \begin{cases} V_h^{G_h+1}, & \text{if } \rho_{h_i}^{G_h+1} = Neutral \\ V_{h_i}^{G_h+1} \sim Normal(V_{h_i}^{G_h}, \sigma\_phase_{j\_i}), & \text{if } \rho_{h_i}^{G_h+1} = Stationary \\ V_{h_i}^{G_h+1} = V_{h_i}^{G_h} - \left(V_{h_i}^{G_h} \times Beta(\alpha\_phase_{j\_i}, \beta\_phase_{j\_i})\right), & \text{if } \rho_{h_i}^{G_h+1} = Depreciation \end{cases} \quad (2.33)$$

The function takes four input variables where the current generation $G$ of a host $h$ is defined as $G_h$ followed by the viral load $V$ present in tissue $i$ of the host at generation G as $V_{h_i}^{G_h}$ and the host's profile $j$ being $h_j$. The population at the next generation $G_h + 1$ is determined by first identifying the tissue's phase in the next generation which is $\rho_{h_i}^{G_h+1}$.

##### 2.2.5.5. Sampling effect

Dependent on a host's profile $h_j$ sampling can have an effect on its infectivity. Apollo allows the host profiles to be configured so that a sampled host can be less infectious or be altogether removed from the infectious population. They can also be configured so that there is no effect on infectivity by sampling. Therefore, if a host's probability of infecting another susceptible host is defined as $I_h$ then if host $h$ is sampled the new probability of infectivity is determined as follows.

$$I(h_j) = \begin{cases} I_h = 1, & \text{if } I_j = \text{"}No\ effect\text{"} \\ I_h \sim Beta(\alpha\_se_j, \beta\_se_j), & \text{if } I_j = \text{"}Less\ infectious\text{"} \\ I_h = 0, & \text{if } I_j = \text{"}Removed\text{"} \end{cases} \tag{2.34}$$

Here function $I$ utilises the profile type ($j$) of the host ($h$) to determine its sampling effect ($I_j$) and the host's new probability of infectivity ($I_h$) is provided. This value will then be used to determine if the host will infect another susceptible host.

#### 2.2.5.6. Epidemiological compartment models

Apollo can be parameterized to support four different types of epidemiological compartment problems namely the SIR (Susceptible Infectious Recovered), SIRS (Susceptible Infectious Recovered Susceptible), SEIR (Susceptible Exposed Infectious Recovered), and SEIRS (Susceptible Exposed Infectious Recovered Susceptible) models.

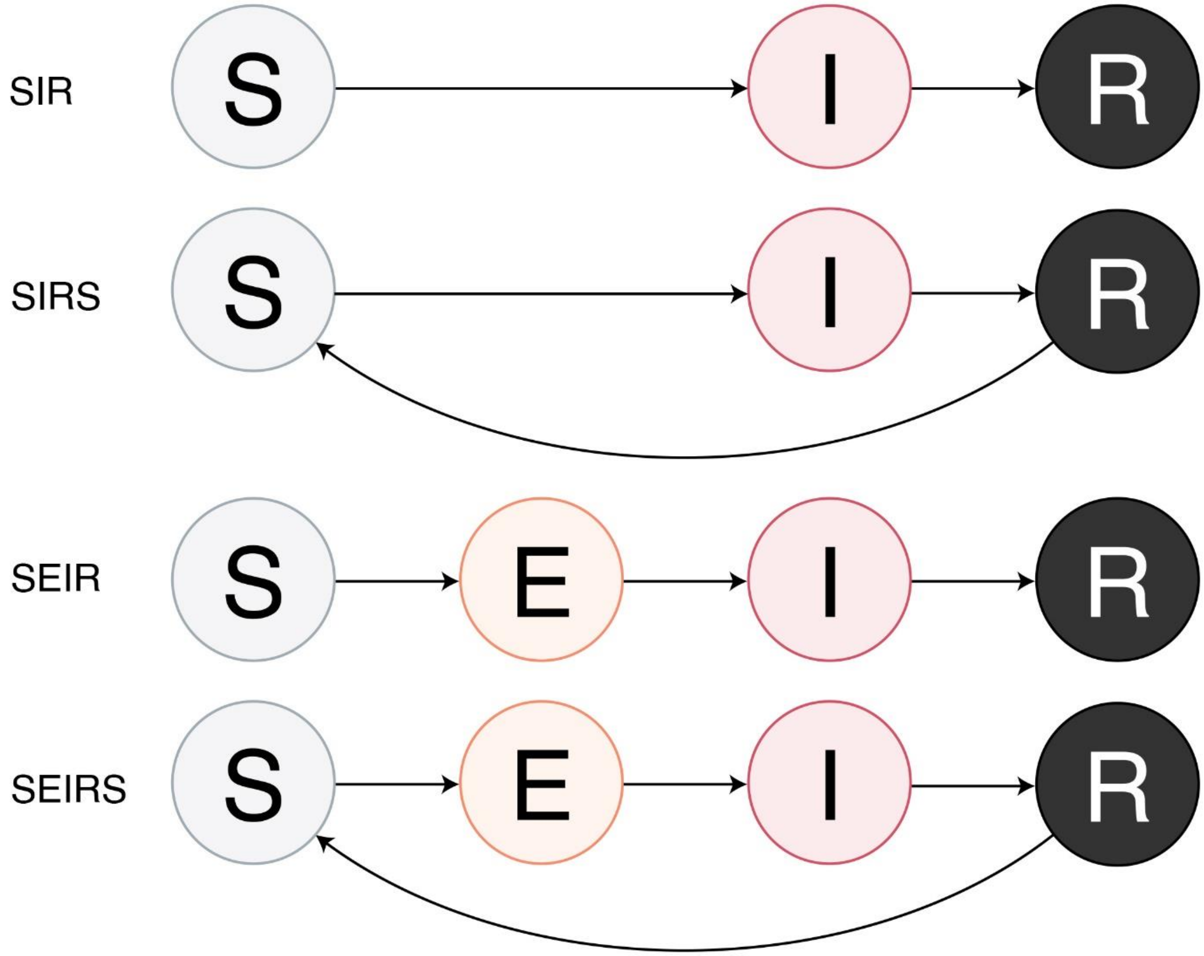

**Figure S - 7.** The four epidemiological compartment models are supported by Apollo. (S) Susceptible, (I) Infectious, (R) Recovered and (E) Exposed.

By setting the infectious tissue ($\pi_3$) load to greater than zero the model becomes a Susceptible to Infectious (SI) model and when it is greater than zero it moves from an SI model to a Susceptible Exposed (E) Infectious (SEI) model. By using the infected to recovered parameter we can switch between having a Recovered (R) and to Susceptible (S).

#### 2.3. Genomic module

The genomic module manages the final hierarchy in Apollo. It is responsible for the configuration and execution of the mechanics related to the evolution of the viral genome and its phenotypes. The genomic model in its simplest form is designed to follow the basic Wright Fisher model's assumptions and through parameterization, these assumptions can be relaxed allowing Apollo to go beyond the Wright Fisher model.

The module at its highest level is responsible for controlling the viral transmission bottleneck for the number of viral particles that will be transmitted from one host to the next. At the genomic level, it's responsible for the control of the viral genome down to the individual bases.

Apollo has dedicated mechanisms to control the evolutionary forces of mutation, proof reading, and recombination. Changes in the viral genome will subsequently have phenotypic effects that will affect the fitness of the new strains, the rate of progeny generation, their probability of surviving till parenthood, and even affect the evolutionary mechanisms themselves.

The genomic module allows the user to first establish reference parameters for factors such as fitness, survivability, and proof reading. These reference parameters also extend into the recombination hotspots in terms of a recombination region's probability of undergoing recombination and the likelihood of selection. Apollo is then able to augment a viral particle's parameters in relation to the reference parameters based on its genomic configuration.

The user is then able to configure the different mutation and recombination regions as needed. Apollo enables overlapping of these regions and their priority of mutation or recombination occurs in the order of configuration. There are a multitude of parameters that the user can use to control these mechanisms including per region, clock models, mutation rates, and site models.

The genomic module is designed to provide the user with a mechanism that allows them to intricately manage the evolution of a genome. In this subsection, we will look at the different parameters and assumptions that govern the genomic module in relation to the standard Wright Fisher model and how it goes beyond it with mathematical validations.

#### 2.3.1. Wright Fisher model

The Wright Fisher (WF) model is one of the most powerful models for capturing the changes in allele frequency by genetic drift. However, the model relies on a number of critical assumptions. The WF model can be described as having no mutation, selection, or migration in a population (**Figure S - 8**).

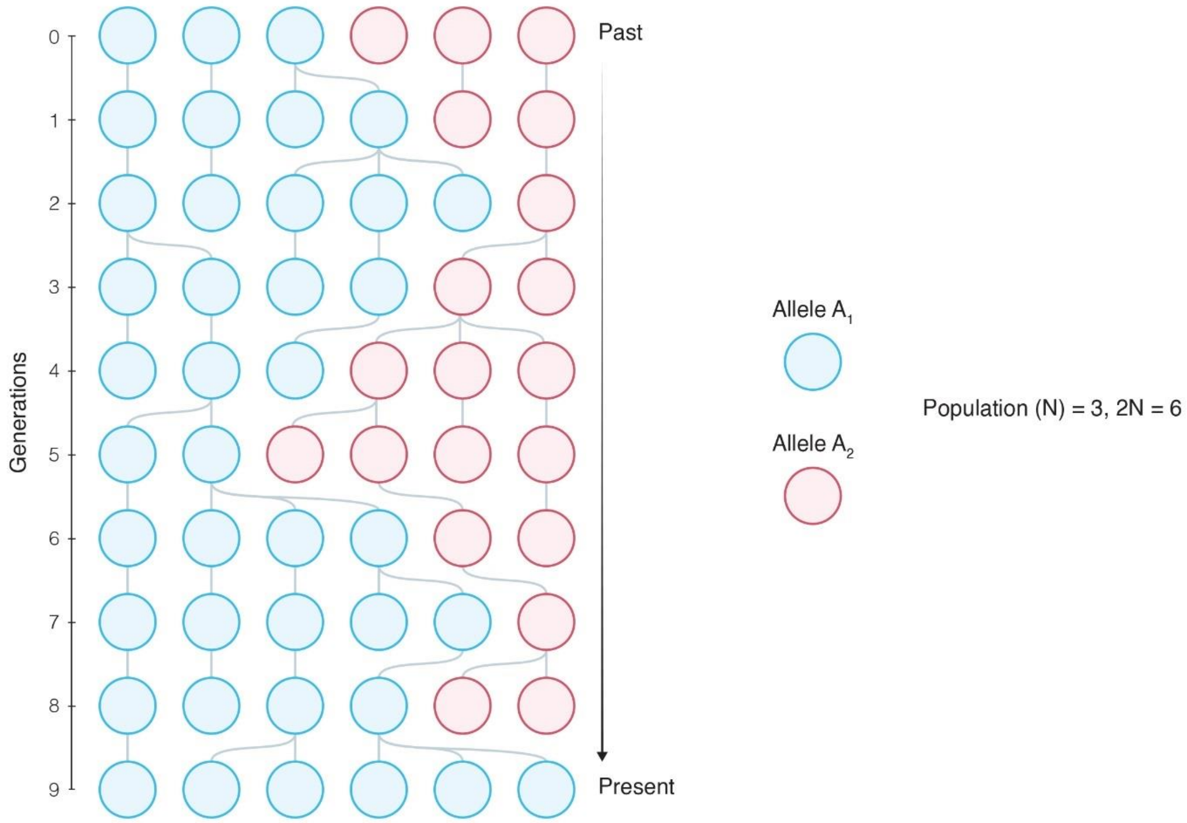

**Figure S - 8.** Simple Wright Fisher model evolution process across 10 generations. Each generation is discrete from the next without any overlapping of individuals. The population is of constant size ($N$) three and therefore the number of chromosomes is six ($2N$). There are two alleles in the population $A_1$ (blue) and $A_2$ (red).$A_1$ has reached fixation where as $A$_2 has been lost.

As depicted in **Figure S - 8** the WF model focusses on diploid populations of a constant size ($N$) where the number of chromosomes in the said population will be $2N$ for every successive generation. The population usually focusses on a single locus ($A$) that's neutral. The locus of a chromosome can be occupied by either one of the two alleles $A_1$ or $A_2$. These loci do not undergo mutation. The populations are discrete with no overlapping of parents and the progeny of the successive generation (at time $t+1$) is determined by the random pairing of an infinite number of gametes from the current population ($t$). This gives the model an inherent Markov property. The change in the allele number across generations occurs solely from genetic drift[8,11,46–48].

Under the WF model if we have $i$ number of $A_1$ alleles at time $t$ the transition probability from $i$ to $j$ at time $t+1$ in the successive generation can be given as follows:

$$P_{ij} = \binom{2N}{j}\left(\frac{i}{2N}\right)^{j}\left(1-\frac{i}{2N}\right)^{2N-j} \quad (2.35)$$

It is quite clear that this is the Probability Mass Function (PMF) of a binomial distribution of $n$ trials, for $k$ successes with a probability of success $p$. Where $k=j$, $n=2N$ and $p = {}^{i}/_{2N}$. The transition probability matrix filled by the above equation will be of size $(2N+1)\times(2N+1)$ (accounting for allele count 0)[49].

Another, interesting factor of the WF model is due to the lack of mutations or migrations $i=0$ or $i=2N$ are absorbing states where fixation occurs for allele $A_2$ and $A_1$ respectively. All other states are transient states. This is an important proposition put forward by the WF model. Under the effects of genetic drift alone as time progresses $\left(\lim_{t\to\infty}\right)$ each neutrally mutating allele will eventually either get lost ($i=0$) or fixed ($i=2N$) in the population[49].

This is the basis of the WF model. The WF model can be adapted to account for factors of mutation, selection and even migrations between populations. We will take a quick look into how this done. But it is apparent these models still adhere to most of these assumptions.

Apollo, at its core is based on the WF model to conduct its forward in time simulations. However, to adapt to the viral model we have to adapt the standard diploid WF model to suit the parameter for viral genomes. Additionally, we attempt to go beyond the WF model to provide the user with a framework that more accurately captures real world evolutionary dynamics.

##### 2.3.1.1. Mutations under the WF model

In the accumulation of mutations, it is first and foremost assumed that all newly arising mutations are neutral in nature. The model also assumes an infinite sites model where no two mutations occur on the same locus implying that every successive descendant of that individual will contain the mutation, and the site will not mutate again. Each site mutates independently of each other. Therefore, these newly arising mutations too will get fixed in the population. This probability of fixation of a neutral mutation has been solved to be equal to its initial frequency in the population which is ${}^{1}/_{2N}$, since at initial incidence, there is only chromosome (heterozygous) with the mutated base[49,50].

##### 2.3.1.2. Selection and mutation under the WF model

The WF model can be adapted to account for selection forces. Intuitively, a population can experience three types of selection on a mutation. They are neutral, beneficial, and deleterious. These three types can be captured by the selection coefficient ($s$).

$$s = \begin{cases} 0, & mutation = \text{"Neutral"} \\ > 0, & mutation = \text{"Beneficial"} \\ < 0, & mutation = \text{"Deleterious"} \end{cases} \tag{2.36}$$

Under this selection coefficient, the probability of sampling a gamete for the next generation ($t + 1$) becomes:

$$P(A_1 sampled) = \frac{k(1+s)}{k(1+s) + 2N - k} \tag{2.37}$$

Where $k$ is the original count for allele $A_1$. Solving for this with values for $s$ we can see that:

$$P(A_1 sampled) = \begin{cases} {}^{1}/_{2N}, & mutation = \text{"Neutral"}, s = 0 \\ > {}^{1}/_{2N}, & mutation = \text{"Beneficial"}, s > 0 \\ < {}^{1}/_{2N}, & mutation = \text{"Deleterious"}, s < 0 \end{cases} \tag{2.38}$$

With this, we can account for newly arising mutations within the WF model. If the proportion of $A_1$ that mutates to $A_2$ is $\mu_1$ and the proportion of $A_2$ that mutates to $A_1$ is $\mu_2$ and $A_1$ and $A_2$ have selection coefficients of $(1 + s)$ and 1 respectively then the proportion of $A_1$ offspring in generation $t + 1$ after the events of selection and mutation can be defined as:

$$\theta_{A_1} = \frac{k(1+s)(1-\mu_1)}{k(1+s)+2N-k} + \frac{(2N-k)(1)(\mu_2)}{k(1+s)+2N-k} \tag{2.39}$$

With this our binomial probability changes from $i/_{2N}$ to $\theta_{A_1}$ resulting in:

$$\text{Number of offspring with } A_1 \sim Binomial(2N, \theta_{A_1}) \tag{2.40}$$

**2.3.1.3. Overall assumptions of the WF model**

As previously stated, the WF model in its simplest form relies on a series of assumptions for both mathematical and computational simplification. These assumptions are six-fold namely, discrete, and non overlapping generations, constant diploid ($2N$ haplotypes) population sizes, equal fitness among all individuals regardless of genome, random mating with no geographical structures, no recombination, and an infinite sites model[8,49,50]. As shown, these assumptions can in fact be relaxed to a certain degree. It also quickly becomes evident in order to form real world simulations in the context of a viral epidemic most of these assumptions have to be relaxed[8]. In the following sections we will look at which assumptions we hold and which of those we relax in Apollo allowing it to go beyond the simple WF model to capture a virus's evolution more accurately.

#### 2.3.2. Beyond the Wright Fisher model

Apollo's foundational framework is designed to follow the WF model. However, through our customizable framework, we allow the user to relax the WF assumptions. This permits the creation of simulations that capture dynamic epidemiological models that could better represent real world scenarios.

##### 2.3.2.1. Discrete generations carried forward from the WF model

Of the six WF model assumptions Apollo is hard coded with one of them. This is the maintenance of discrete nonoverlapping generations. Here the individuals of one generation are not carried forward to the next generation, even if they do not take part in progeny formation in the current generational cycle.

Now we are ready to look at the remaining six assumptions and how they are relaxed in Apollo.

##### 2.3.2.2. Migration from diploid to haploid genomes

We can begin with the most straightforward deviation from the Wright Fisher model. In the context of viral evolution, the shift of ploidy from diploid (ploidy = 2) to a haploid (ploidy = 1). The solution is simply changing the instances of $2N$ in the WF model equations to $N$. Therefore, number of genomes becomes equal to the population size. This is a hard coded change fixed into Apollo's infrastructure.

##### 2.3.2.3. Dynamic population size

Apollo allows the fluctuation of population size across successive generations. This is dependent on several factors, the simplest being the number of progenies being produced in the previous generation. By default, the generation phase of a tissue (compartments within the host where the life cycles occur) is set to stationary. This causes the simulation to determine the next generation's population around a normal distribution based on the current generation's population. If the variance is set to zero, then the simulation will behave as a WF model by maintaining a fixed population size for each successive generation.

However, if the generation phase is neutral, Apollo is free to have a dynamic population size, or if its depreciation the population size will fall based on a Beta distribution.

##### 2.3.2.4. Genetic variability shapes fitness

Apollo is designed to account for the effects of mutations on a genome and allow the mutational effects to collectively influence the evolution of the virus. A factor affected by the genome is fitness. Therefore, the variability of the genome will change the fitness of a virus.

Under the WF model fitness is defined as the capability of an individual to produce subsequent progeny of the same genotype[50–52]. In the WF model, this is captured by the selection coefficient $s$. The definition of fitness can account for two factors. The fitness of the individual, namely the number of progenies it leaves behind, and the fitness of the genome which is the number of copies of itself it leaves behind[50].

Apollo accounts for these two fitness definitions using a combination of two parameters. Namely the similarly named fitness parameter ($f$) and the survivability ($z$) parameter. The fitness parameter affects the number of progenies that will be produced by a parent whereas the survivability parameter is the probability that a progeny will survive till reproduction. To allow for better capture of the changes in these parameters Apollo uses reference parameters for them. This is the value that a non mutated standard genome will have.

Given a reference parameter ($F \in [0, \infty]$) we can show that fitness ($f \in [0, \infty]$) being a multiplicative parameter captures selection as:

$$f = \begin{cases} f = 1, & mutation = \text{"Neutral"} \\ f > 1, & mutation = \text{"Beneficial"} \\ f < 1, & mutation = \text{"Deleterious"} \end{cases} \tag{2.41}$$

and for survivability ($z \in [-\infty, +\infty]$) being an additive parameter given a reference parameter ($Z \in [0,1]$):

$$z = \begin{cases} z = 0, & mutation = \text{"Neutral"} \\ z > 0, & mutation = \text{"Beneficial"} \\ z < 0, & mutation = \text{"Deleterious"} \end{cases} \tag{2.42}$$

With this let us see how Equation (2.39) can be adapted to fit Apollo's framework.

If the number of progeny ($P$) produced by a viral genotype strain $v$ is $P_v$ ($v \in \{0,1,\ldots.V-1\}$ where $V$ is the total number of genotypes in the population) then $P_v$ can be determined as follows.

Using the function $S(d\_type, parameters)$ we first determine the standard number of progenies that will be produced:

$$S(d\_type, parameters) = \begin{cases} P_{v\_standard} \sim NegativeBinomial(n_p, p_p), \text{if } d\_type = \text{"}Negative\ binomial\text{"} \\ P_{v\_standard} \sim Poisson(\mu_p), \text{if } d\_type = \text{"}Poisson\text{"} \\ P_{v\_standard} \sim Gamma(\alpha_p, \beta_p), \text{if } d\_type = \text{"}Gamma\text{"} \end{cases} \quad (2.43)$$

Where $d\_type$ is the distribution selected to determine the progeny generation and the $parameters$ are the selected distributions relevant parameters.

With that if the fitness of that genotype is given by $f_v$ then we can determine the total progeny produced as:

$$P_v = P_{v\_standard} \times F \times f_v \text{ where } P_v = \{n \in A: n \geq 0\} \quad (2.44)$$
$$\text{where } A \text{ represents all integers}$$

Similarly, as before letting us have $k$ number of parents from a total parent population $N$ with the genotype of $v$ and they mutate with a rate of $\mu_1$ to form other variants $x$ where ($\forall x \in \{0,1,\ldots.V-1\}, x \neq v$) and all $x$ mutate at a rate of $\mu_2$ to form $v$. Then we can modify Equation (2.39) as follows:

$$\theta_v = \frac{k(P_v)(1-\mu_1)}{k(P_v)+(N-k)(P_x)} + \frac{(N-k)(P_x)(\mu_2)}{k(P_v)+(N-k)(P_x)} \quad (2.45)$$

We can take this further to incorporate the proportion of progeny of genotype $v$ that survive till reproduction ($\theta_{v\_survive}$) by factoring in our survivability parameter ($z$) as follows:

$$Z_v = \begin{cases} 0, & Z+z_v < 0 \\ 1, & Z+z_v > 1 \\ Z+z_v, & if\ 0 \leq Z+z_v \leq 1 \end{cases} \quad (2.46)$$

$$\theta_{v\_survive} = \frac{[k(P_v)(1-\mu_1)]Z_v}{k(P_v)+(N-k)(P_x)} + \frac{[(N-k)(P_x)(\mu_2)]Z_v}{k(P_v)+(N-k)(P_x)} \quad (2.47)$$

Certain viruses such as SARS CoV-2 are equipped with proofreading machinery. These mechanisms help the viral replication mechanism to prevent base mismatches reducing mutations. Apollo incorporates the proofreading machinery into its genome module.

To see how the proof-reading mechanism works let's take a closer look at the mutation rate parameter $\mu$. We can see that $\mu_v$ is the probability that a genome $v$ mutates to genome $x$. This value can be determined as follows:

$$\mu_v = \frac{n_{v\_x}}{n_v} \tag{2.48}$$

where $n_{v\_x}$ is the number of individuals of genome $v$ mutated to genome $x$ given an original total of individuals with genome $v$ being $n_v$.

We can then use $\mu_v$ to determine the number of $x$ genome individuals produced by $v$ using a binomial distribution:

$$n_{v\_x} \sim Binomial(n_v, \mu_v) \tag{2.49}$$

The proof-reading mechanism in Apollo is defined by the probability of catching a base mismatch ($\sigma \in [0,1]$). For here we can simply take it as the probability of catching a mutated genome. Similar to survivability this is an additive parameter and given a reference parameter ($R \in [0,1]$). With this we can determine the overall proof-reading accuracy ($R_v$) of a genome $v$ as:

$$R_v = \begin{cases} 0, & R + \sigma_v < 0 \\ 1, & R + \sigma_v > 1 \\ R + \sigma_v, & if\ 0 \leq R + \sigma_v \leq 1 \end{cases} \tag{2.50}$$

We can then determine how many of the mutated genomes are rectified using another binomial distribution:

$$\grave{n}_{v\_x} \sim Binomial(n_{v\_x}, \mathrm{R_v}) \tag{2.51}$$

Therefore, the actual number of mutated genomes would then become:

$$N_{v\_x} = n_{v\_x} - \grave{n}_{v\_x} \tag{2.52}$$

With that we can see that the mutation rate ($\mathrm{\mu_v}$) decreases with the proof-reading mechanism as the new mutation rate with proofreading ($\grave{\mu}_\mathrm{v}$) would be:

$$\grave{\mu_v} = \frac{n_{v\_x} - \grave{n}_{v\_x}}{n_v} \quad (2.53)$$

Apollo only accounts for the proofreading machinery if it is activated (α).

$$\grave{\mu_v} = \frac{n_{v\_x} - \left[\left(\grave{n}_{v\_x}\right) \times \alpha\right]}{n_v} \quad (2.54)$$

$$\text{Where } \alpha = \begin{cases} 1, & \text{activated} \\ 0, & \text{deactivated} \end{cases} \quad (2.55)$$

This is a simplified form of what happens internally in Apollo's architecture. Due to the GPU acceleration, Apollo accounts for each individual's genome separately in parallel, therefore providing an accurate exact count of progeny formed, their mutations, population frequency, and subsequently the progeny survived till reproduction.

##### 2.3.2.5. Accounting for genotype variability

Given a viral particle's genome, Apollo calculates its phenotypic factors such as overall fitness survivability, and proofreading accuracy. The user can configure different loci and the contributions made by the occupying allele. The overall influence (E.g.: $f_v$, $z_v$) of the genotype is determined by accounting for all contributing loci.

First, let's determine the different bases that can occupy a locus. Apollo identifies the four bases Adenine ($A = 0$), Thymine ($T = 1$), Guanine ($G = 2$), and Cytosine ($C = 3$).

If set $C$ is our sorted collection of contributing positions in the genome (E.g.: $C \in \{0,20,100\}$) for our contributing loci. We can then iterate through these positions to determine the total effect on the factor under study.

The genetic makeup of these positions can be arranged via $|C| \times 4$ matrix ($G$), where the columns represent the base occupied in the order of $A, T, G, C$, and the occupied base will hold the value 1 while the rest hold 0.

Similarly, we can use a matrix notation to represent the contribution of these bases to a particular factor. Let's have this matrix be $E$ and it too will be of size $|C| \times 4$. However, here the contribution of a particular base at a given loci to the factor will occupy each cell.

Let's take a look at a quick example to understand the above statements better. Consider our factor as fitness ($f$) and fitness is affected by five loci. Therefore, our set $C$ with the positions would be $C \in \{7,100,110,200,301\}$ and the cardinality of the set is $|C| = 5$. Let's define our genome matrix ($G$) and the effect matrix ($E$) for fitness as:

$$\mathrm{G} = \begin{bmatrix} 1 & 0 & 0 & 0 \\ 0 & 0 & 0 & 1 \\ 0 & 1 & 0 & 0 \\ 0 & 0 & 1 & 0 \\ 0 & 1 & 0 & 0 \end{bmatrix}, E = \begin{bmatrix} 0.5 & 1 & 1.5 & 1 \\ 1 & 3 & 0.5 & 1 \\ 1 & 2 & 1 & 0.2 \\ 1 & 1 & 0.9 & 1 \\ 1 & 1.7 & 1 & 1 \end{bmatrix}$$

From the above matrix $G$ (rows = positions and columns = bases) we can see that the positions in $C$ are respectively A, C, T, G, and T. We can get the overall fitness matrix ($\hat{f}$) for this genome by performing an element wise multiplication of the two matrixes:

$$\hat{f} = \mathrm{G}.* \mathrm{E} \tag{2.56}$$

For this example, above $\hat{f}$ will be:

$$\hat{f} = \begin{bmatrix} 0.5 & 0 & 0 & 0 \\ 0 & 0 & 0 & 1 \\ 0 & 2 & 0 & 0 \\ 0 & 0 & 0.9 & 0 \\ 0 & 1.7 & 0 & 0 \end{bmatrix}$$

Then to get the total overall fitness value ($f$) we can multiply all the values in the matrix ($\hat{f}$) together. We multiply the values since $f$ is a multiplicative factor. If it was an additive factor like survivability ($z$) we can add all the values together.

$$f = \prod_{i=1}^{|C|} \prod_{j=1}^{4} \widehat{v_{ij}} \text{ or } z = \sum_{i=1}^{|C|} \sum_{j=1}^{4} \widehat{z_{ij}} \tag{2.57}$$

The above values can then be plugged into their respective equations such as Equation (2.44), Equation (2.46) or Equation (2.50). In this manner, Apollo can account for each genome's phenotypic response. The actual algorithmic implementations are designed to be more efficient than their mathematical counterparts using large scale parallel processing techniques and algorithmic optimizations.

##### 2.3.2.6. Base mutations

Now that we have seen how Apollo accounts for genomic diversity, it is important to understand how this diversity occurs. The simplest and straightforward mechanism is mutations. Apollo accounts for point mutations and substitutions[53–55]. Mutations typically occur spontaneously during DNA replication, these mutations can then become fixed in the population and become inherited by the subsequent progeny produced by the parent genome. Mutations guide the evolution of a species in a population[51,56].

Apollo uses the molecular clock hypothesis to induce mutations into a genome. Here different regions of the genome can be configured with different mutation rates and site models[57,58]. Each site in the genome mutates independently of one another. A mutating genome is a dynamic entity, and different areas can undergo different rates of mutation based on factors such as the selection pressures acting on these regions. Apollo identifies a mutating region as a hotspot. The user can configure multiple hotspots on a single genome, with overlaps (**Figure S - 9**).

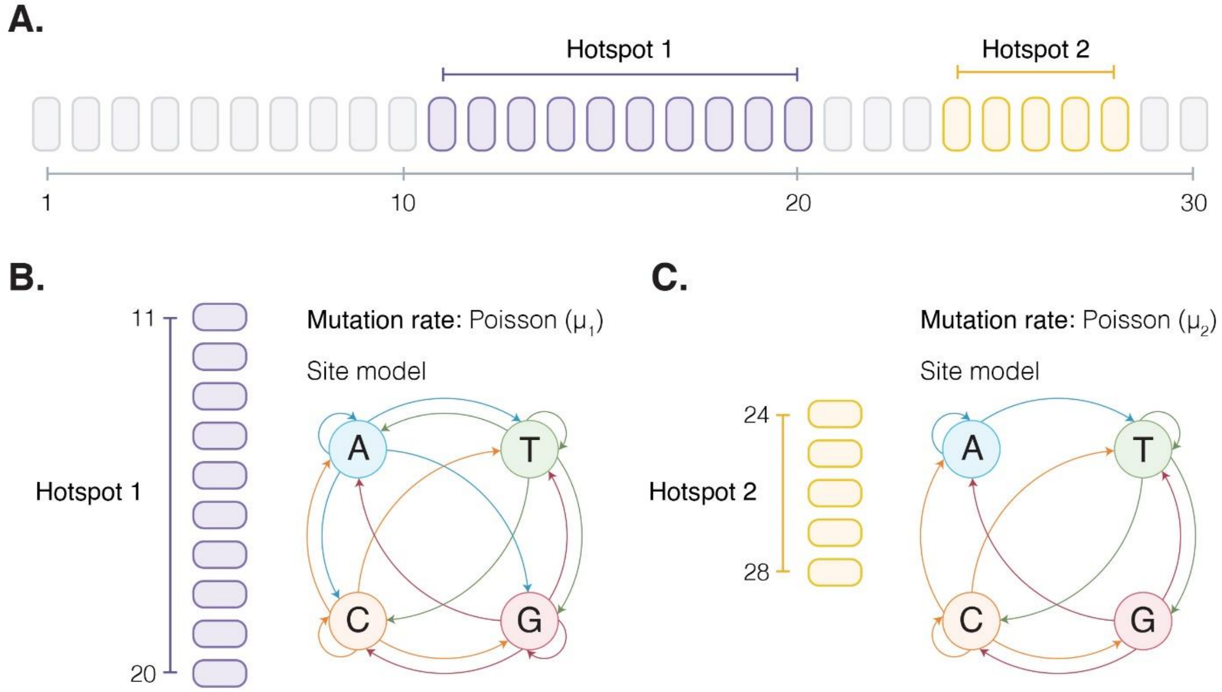

**Figure S - 9.** Configuration of a genome with two mutating hotspot regions (Hotspot 1 in purple and Hotspot 2 in gold). (A) Depicts the position of the hotspot regions in the genome. (B and C) Each region has its own mutation rates and site models that govern base substitution.

The molecular clock hypothesis states that DNA sequences evolve at a relatively constant rate. A direct consequence of this phenomenon is that the genetic difference between two species is observed to be proportional to the time since their Most Recent Common Ancestor (MRCA)[59,60]. Therefore, the molecular clock model governs the mutation rate, and it typically follows a Poisson distribution[61]. But with Apollo, we have provided the user with an additional two methods of a negative binomial distribution and fixed mutations per generation. At first glance, it might appear that Apollo follows a strict molecular clock paradigm, where a consistent mutation rate is maintained across all species. However, due to the presence of our proofreading mechanic, Apollo can be modeled to simulate relaxed molecular clock behavior. Since the proofreading mechanism controls the mutation rate and mutations in turn influence the proofreading accuracy; a relaxed molecular clock model can be adapted, where certain strains experience faster mutation rates than others.

The site model is used to define the path of nucleotide evolution from one base to the next. This is a nucleotide substitution model and helps determine the replacement of one base with another during a mutation event. Apollo does not formally recognize transition (purine (A and G) to purine or pyrimidine (T and C) to pyrimidine) and transversion (purine to pyrimidine or pyrimidine to purine) events, but it allows control of all 16 possible base substitution events by implementing a complete substitution model (**Figure S - 9B**). Using this complete model the user is free to parametrize these events. Even taking into account transitions and transversions. It should be noted that Apollo does not follow an infinite sites model for mutations. This means that a locus has the potential to undergo multiple rounds of mutations.

Let's now look at how Apollo configures its mutational hotspots and induces base changes. Apollo supports an unlimited number of mutational hotspots ($\mathrm{M} \in \{1,2, \dots \infty\}$). These hotspots ($m \in \{1,2, \dots M\}$) will be defined by the start and stop base positions they span across with each having their own clock model ($C$) and transitional matrix ($S$ of size $4 \times 4$ allowing 16 possible base changes) representing the site model. Therefore, the number of mutations ($\gamma$) that a site experiences during a given generation ($g$) can be obtained via the following function $\mathrm{f}\left(C_m, params, \gamma_{m\_g}\right)$:

$$
\mathrm{f}(C_m, params, \gamma_{m\_g}) = \begin{cases} \gamma_{m\_g} \sim Poisson(\mu_m), & C_m = "Poisson" \\ \gamma_{m\_g} \sim NegativeBinomial(n_m, p_m), & C_m = "Negative\ Binomial" \\ \gamma_{m\_g} = \tau \text{ where } (\tau \in \{0 \ldots \infty\}), & C_m = "Fixed" \end{cases} \tag{2.58}
$$

Once the number of mutation events ($\gamma_{m\_g}$) has been obtained we can select the bases that undergo mutation (**Figure S - 10**). The mutation events are processed sequentially (**Figure S - 10E**). The position undergoing mutation is selected using a uniform distribution, the range being the span of the hotspot. These bases would then change based on the hotspot's ($m$) transition matrix ($S_m$) **Figure S - 10D** and **E**.

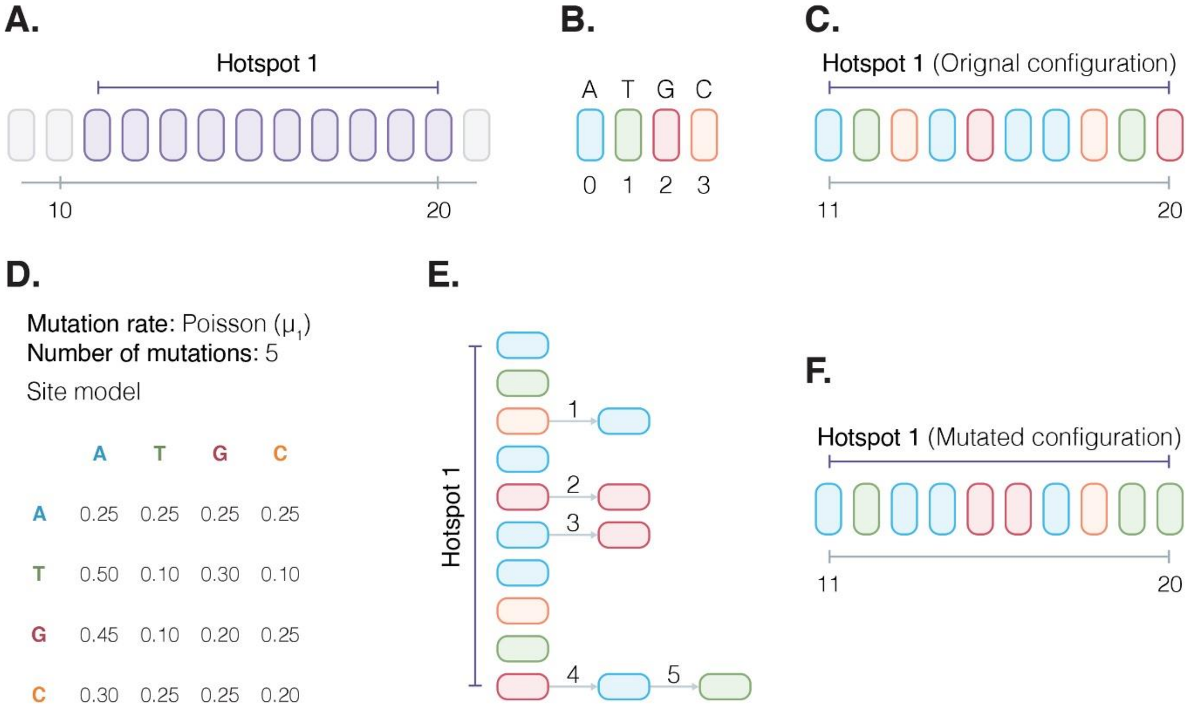

**Figure S - 10.** Detailed depiction of the mutation process. (A) Apollo identifies the target hotspot ($m = 1$) region based on the start and stop positions. (B) Coloured coded bases with their letter and numeric index representations. (C) Coloured coded hotspot region representing its nucleotide configuration. (D) The respective hotspot region's parameters for mutation rate and base substitution. In the current generation's ($g$) mutation event, the Poisson distribution has drawn five mutations ($\gamma_{1\_g}$) to occur on the hotspot. (E) The five mutations occur across four base positions with base position 20 experiencing two consecutive mutations. (F) The mutated hotspot region with its new nucleotide base configuration.

The determination of the mutated base given the current base is a transition matrix ($S_m$) multiplication with the vector of the position's base configuration ($v_{p\_g}$). For instance, let's take the mutation of position ($p$) 13 in **Figure S - 10** to see this worked out assuming it's at generation ($g$) 0.

$$S_1 = \begin{bmatrix} 0.25 & 0.25 & 0.25 & 0.25 \\ 0.50 & 0.10 & 0.30 & 0.10 \\ 0.45 & 0.10 & 0.20 & 0.25 \\ 0.30 & 0.25 & 0.25 & 0.20 \end{bmatrix},\ v_{13\_0} = [0 \quad 0 \quad 0 \quad 1]$$

We can then determine the probability of the next base by:

$$v_{p\_(g+1)} = v_{p\_g} \times S_m \tag{2.59}$$

$$\therefore v_{13\_1} = v_{13\_0} S_1 = [0 \quad 0 \quad 0 \quad 1] \begin{bmatrix} 0.25 & 0.25 & 0.25 & 0.25 \\ 0.50 & 0.10 & 0.30 & 0.10 \\ 0.45 & 0.10 & 0.20 & 0.25 \\ 0.30 & 0.25 & 0.25 & 0.20 \end{bmatrix} = [0.30 \quad 0.25 \quad 0.25 \quad 0.20]$$

As observed in **Figure S - 10E** the base mutated to an A (Adenosine) as it had the highest probability of 0.30 ($v_{13\_1} = [1 \quad 0 \quad 0 \quad 0]$ as shown in **Figure S - 10F**). This is achieved computationally using a cumulative probability technique. In this manner Apollo is capable of simulating the induction of mutations governed by clock and site models.

#### 2.3.2.7. Recombination

Recombination is the exchange of pieces of genetic material between molecules of DNA to produce genetic variants with new combinations. Mutations generally create subtle genetic changes while recombination results in major changes. It has important implications for the evolution of organisms. Haploid viral genomes have displayed adaptions to accommodate recombination in their replication cycles during events of coinfection of cells. Coinfection is when more than one viral particle infects a host cell.

Apollo implements a template switching by recombination model (**Figure S - 11**). This is where the polymerase pauses during replication at specific sequences and then switches to a different parent's template strand for that target sequence before resuming the replication. These strands from different parents will then recombine to form the new progeny's chimeric genome[26,62,63].

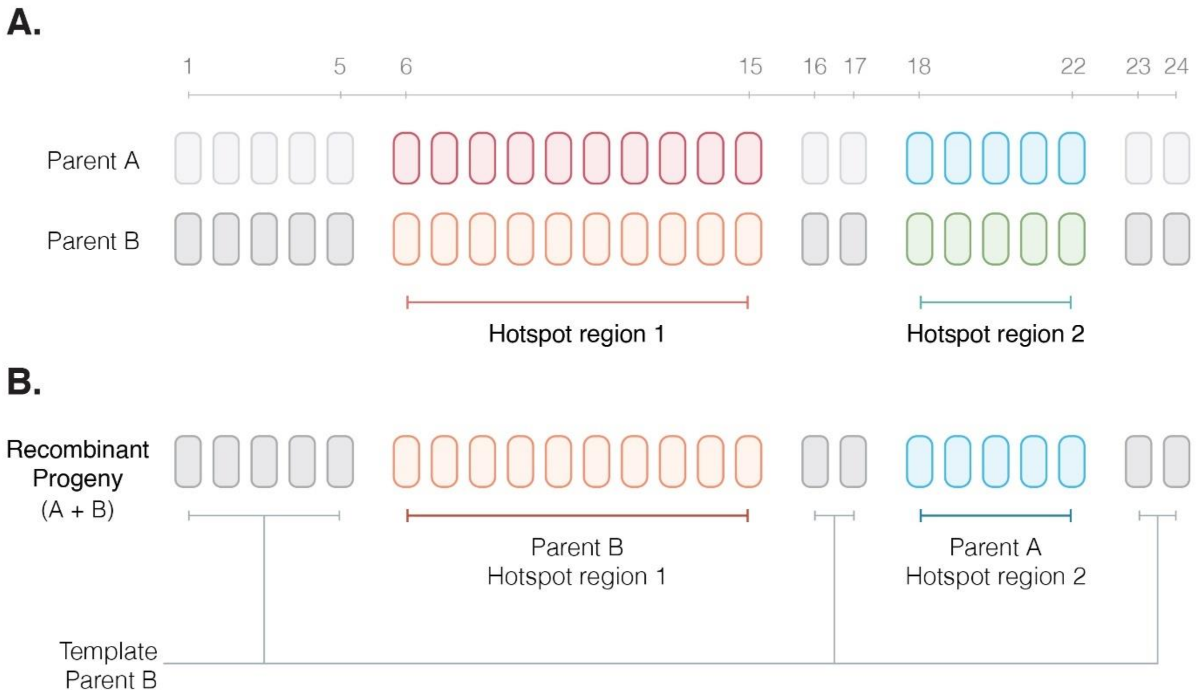

**Figure S - 11.** The formation of recombinant progeny genomes. (A) Two parents, A and B that have coinfected a cell. Their genomes comprise of two recombination hotspots, each with their own nucleotide configurations. The resultant progeny (B) formed will be a chimera of the two parents A and B. Here the parental template is from parent B, who also contributes its genome to hotspot one, while hotspot two's region is configured using parent A's genome.

The recombination mechanism is triggered during the process of progeny assembly. By this time the parent genome of the progeny that will act as its template and each recombinant hotspot's respective doner parents are selected. Here the template genome is the main parental genome and the recombinant hotspot's donors are those regions' parents, together they form the new progeny's pedigree. Essentially the new progeny's parents are all viral particles that produced its genome. It can be understood that even though viruses do not take part in sexual reproduction, the force of recombination allows the exchange of genetic material in a similar manner.

Apollo allows an unlimited number of recombinant hotspots ($R \in \{1,2,\dots,\infty\}$) and each hotspot ($r \in \{1,2,\dots,R\}$) has two parameters that can be subjected to phenotypic change based on the genome. These parameters are namely the probability of recombination ($\varphi \in [-\infty,+\infty]$) and selectivity ($\omega \in [0,\infty]$).

The probability of recombination is essentially the probability that the hotspot region of a particular viral particle will undergo recombination. We account for this factor due to studies revealing the rise of recombination events in a region due to the occurrence of mutations in another. For instance in the SARS-CoV-2 genome's 427N and 436Y variant point mutations that gave rise to the occurrence of recombination events in the spike protein[64]. Selectivity is the likelihood that the current viral particle's hotspot genome will be selected during a recombination event during progeny formation. This is to account for the fact that genetically similar or closely related gene regions have a higher likelihood of undergoing recombination[65].

Both parameters similar to the fitness, survivability, and proof-reading parameters have reference parameters; $E$ ($E \in [0,1]$) for the probability of recombination and it is an additive parameter and $L$ ($L \in [0,\infty]$) for selectivity is multiplicative. Similar to fitness and survivability they contribute to a recombinant hotspot's selection forces as follows:

$$\omega = \begin{cases} \omega = 1, & mutation = \text{"Neutral"} \\ \omega > 1, & mutation = \text{"Positive"} \\ \omega < 1, & mutation = \text{"Negative"} \end{cases} \tag{2.60}$$

$$\varphi = \begin{cases} \varphi = 0, & mutation = \text{"Neutral"} \\ \varphi > 0, & mutation = \text{"Positive"} \\ \varphi < 0, & mutation = \text{"Negative"} \end{cases} \tag{2.61}$$

We can calculate a particular particle's ($i$) hotspot's ($r$) overall probability of recombination ($O$) and selectivity ($T$) in relation to their reference parameters as follows:

$$O_{i\_r} = \begin{cases} 0, & E + \varphi_{i\_r} < 0 \\ 1, & E + \varphi_{i\_r} > 1 \\ E + \varphi_{i\_r}, & if\ 0 \leq E + \varphi_{i\_r} \leq 1 \end{cases} \tag{2.62}$$

$$T_{i\_r} = L \times \omega_{\mathrm{i\_r}} \tag{2.63}$$

A region undergoing recombination ($k_{i\_r}$) during the replication process is determined using a Bernoulli distribution.

$$k_{i\_r} = Bernoulli(O_{i\_r}) \tag{2.64}$$

The probability of a particular particle's ($i$) hotspot ($r$) being selected ($U_{i\_r}$) as the template, given a total of $j$ parental genomes co infecting the cell is:

$$U_{i\_r} = \frac{T_{i\_r}}{\sum_{x=1}^{j} T_{x\_r}} \tag{2.65}$$

Then using a cumulative probability technique computationally, we then select the particle whose genome will contribute to the hotspot given $k_{i\_r} = 1$.

It should be noted that the process of recombination occurs before mutation. The recombination step occurs during progeny formation and once the progeny sequence has been assembled the mutations are induced.

### SECTION 3. SOFTWARE ARCHITECTURE

Apollo is founded on CATE's (CUDA Accelerated Testing of Evolution) architecture. CATE is our first large scale parallel processing solution. It utilizes all state-of-the-art parallel processing hardware technologies of the GPU, CPU, and SSD. CATE's architecture is adaptable to the available hardware resources allowing for an adaptable software solution[1,15]. Motivated by this success we decided to expand on these innovations and use our proven architecture to drive Apollo's simulations.

At its core Apollo utilises a large-scale parallel processing architecture. Powered by an algorithm that synergistically navigates its processes across the GPU, CPU, and even SSDs. Apollo similar to CATE is capable of adapting to the presence of the SSD and in its absence is capable of working on traditional HDDs (Hard Disk Drives). Similar to CATE, Apollo also uses a fragmented data storage structure. This fragmented structure allows the implementation of additional parallel processing strategies that further accelerate Apollo's computational speeds and memory efficiency.

Apollo's architecture which we will discuss in detail in this section is designed to allow the aforementioned three modules of Network, Host, and Genome to work synergistically with the parallel processing hardware. The architecture orchestrates the simulations to be conducted at accelerated speeds and resource efficiency.

The main architecture is comprised of three main steps. The first being parametrization, where the user manually configures the availability of the hardware resources, as well as the parameters associated with the simulation. This is followed by the second step of initialization. Here the parameters are validated, the required data are generated, and the prerequisites required for the simulation are generated. Third and finally the simulation engine where the epidemic is simulated complete with host-to-host infection, within host viral infection across tissues and viral mutation and phenotypic expression.

In this section, we will look into this three-step architecture in detail. However, we will begin with Apollo's use of parallel processing technologies first, followed by its file architecture and data structures. These implementations are what enable accelerated large-scale simulations with a high efficiency in hardware resource usage.

#### 3.1. Central Processing Unit (CPU) vs Graphical Processing Unit (GPU)

The technology of parallel processing involves the action of running multiple instructions at the same time. In modern computing two hardware technologies are available with the capability of parallel processing. These are the CPU and GPU. The CPU and GPU are designed for the execution of instructions or processes. However, the GPU in contrast to the CPU is specially designed for this function of parallel processing. The primary difference between the GPU and CPU is in their number of cores (**Figure S - 12**). Theoretically, the more cores a system has the more tasks it can perform in parallel[1,66–69].

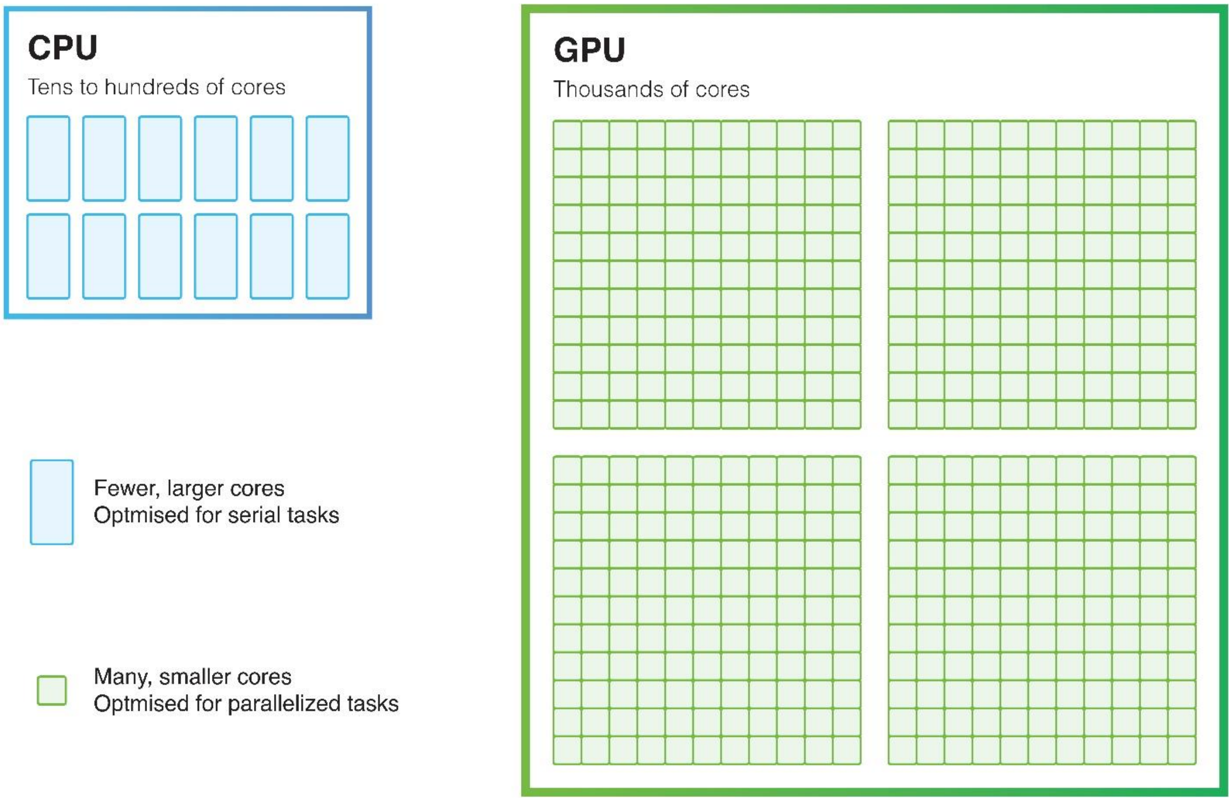

**Figure S - 12:** Architecture comparison of the CPU and GPU. The CPU comprises of several cores. The CPU as a whole is optimized for parallel processing tasks with low latency. In contrast, the GPU has smaller cores but the system as a whole is optimized with a denser core structure for parallel processing. Therefore, GPUs with their extremely large core count is able to outperform CPUs in tasks that require large scale parallel processing.

The GPU has a large number of smaller less powerful cores. These cores with the GPU’s architecture are specifically designed and optimized for parallel processing. The CPU, on the other hand, has fewer cores. However, these cores on their own are more powerful in terms of compute speed. However due to the sheer volume of cores that occupy the GPU, it becomes magnitudes faster than a CPU[70].

Apollo’s algorithm is designed to strike a balance between the CPU and GPU in the allocation of processes between the two technologies. Tasks that will usually occupy a smaller number of parallel processes, for instance, the configuration of each host in the network are assigned to CPU based parallel processing. Whereas tasks that require a large number of parallel processing tasks, for instance simulating the mutations of each individual viral particle will be conducted via the GPU.

In CATE, we utilized a single GPU architecture. This meant that CATE was able to assign the parallel processing tasks only to a single GPU. In Apollo, we have expanded on this implementation to accommodate multiple GPUs. Apollo is able to delegate its processes among the GPUs provided for its use, allowing for a greater parallel processing architecture.

#### 3.2. Segmented file structure

One of the key innovations of CATE that contributed to its unprecedented processing speeds was its segmented file structure. Apollo too enjoys the benefits of this divide and conquer data storage strategy. During the simulation of viral particles, the individual viral sequences must be stored. Therefore, we use a segmented storage structure to store the files to be readily accessed during the simulations.

The sequences will be stored in batches of a user determined size. For instance, if the user has set the "Intermediate Sequences per file" parameter to 10,000, the maximum number of sequences that will be stored in a sequence file will be 10,000 sequences and in the event of more sequences, they will be stored in a new sequence file. Similar to CATE the file names are used to index each segment recording the indexes of the first and last sequences stored in the file. This in turn allows the generation of a sorted list for the file structure that can then be traversed using quick search algorithms, including our own Compound Interpolated Search (CIS) algorithm[1]. This solution to the data retrieval problem not only increases the speed of the computations but also provides the additional benefit of allowing the software to be memory efficient. As we can precisely control the data being read into the memory.

Furthermore, Apollo allows a series of folder and file management controls. Due to the large volume of sequences that can be simulated using Apollo, the segmented file structure can cause the generation of a large number of files, that will in turn require a larger footprint on the user's on-device storage. To accommodate for this Apollo provides the user with the option to compress and pack completed generational data.

The segmented file structure also enables the full use of SSD storage. Allowing multiple files to be read and processed in parallel, instead of sequentially processing one file at a time. Apollo's algorithm is adaptable to both SSDs and HDDs. In the absence of SSD storage, it will revert to a single file read, but the algorithm is designed to remove any redundancy during data reads and have as minimal difference as possible in comparison to SSD file access. Such strategies are possible due to the segmented file structure being sorted.

#### 3.3. Overview of Apollo's architecture

As mentioned before Apollo's architecture can be summarised into three main steps namely, parameterization, initialization, and finally simulation engine. The layout of these steps in the context of its overall architecture is shown in main text **Figure 1**. In this sub section, we will look at each of these steps in detail and how they work synergistically with Apollo's modules (SECTION 2).

##### 3.3.1. Parametrization

The parametrization step allows the user to configure the simulation across all three hierarchical levels of the population network, within host dynamics and tissue structures and viral genomics including mutation and recombination factors. The parametrization step is also where the user will configure the hardware resources available. This includes the number of CPU cores, the number of GPUs available for computational data processing as well as the availability of SSDs. These user specified parameters will then govern the simulation including its speed and efficiency dependent on the resources available. The parameterization is done using JSON scripting.

##### 3.3.2. Initialization

The initialization step begins with the validation of the user specified parameters. Apollo checks for any discrepancies in relation to the options selected and the variables that have been declared. Following the validation of the parameters, the relevant data structures are created, and prerequisites are configured.

Then Apollo begins with the first step of the simulation. This is the generation of the contact network. The contact network governs the spread of the infection across the population. Once the contact network has been established Apollo will distribute the host profiles and assign the different nodes with their profile. Based on the assigned host profiles their tissue structures and other unique factors will be allocated. Finally, at random, a host will be selected to be infected with the reference sequences.

##### 3.3.3. Simulation engine

The simulation engine is responsible for the seamless integration of Apollo's modules in tandem with the user defined parameters to orchestrate the intended epidemiological simulation. The sophisticated algorithm is comprised of multiple components each optimized to execute its intended tasks in accelerated manner while being resource efficient. As shown in the main text **Figure 1** the simulation engine comprises of 12 sub tasks. We will understand the architecture of the simulation engine by discussing each task independently.

###### 3.3.3.1. Compartment classification

Apollo has three main epidemiological compartments. The first being the Susceptible (S) compartment comprises of individuals who are not infected with the viral population but are able to be infected. Secondly, Infected (I) are individuals who are actively harboring the viral population, and finally third we have the recovered population. The Recovered (R) population may comprise of individuals who have ceased to exist due to mortality from the infection or those who have been infected and now have recovered and are not susceptible any longer. The factors of being infectious and mortality are determined using the viral load present within the specific tissues of the host.

However, even though Apollo has the three main compartments of SIR you may note that as discussed previously (2.2.5.6 and **Figure S - 7**) we can go beyond the simple SIR model to SIRS, SEIR, and SEIRS. Apollo is able to enjoy these alternatives due to the role of tissues (2.2.3.1). For instance, once a susceptible host is infected with the virus, we can prevent them from immediately infecting other susceptible individuals by setting a required viral load that has to be reached before the host becomes infectious. This will then allow us to incorporate an Exposed (E) period. To allow recovered individuals to become infectious Apollo comes equipped with a parameter that enables reinfection of recovered hosts. These hosts will move from the recovered population back into the susceptible population.

##### 3.3.3.2. Infecting hosts

Once Apollo has sorted the individuals in the new generation to their relevant compartments it will then begin by infecting new individuals from the infectious population. Apollo also has the option to reinfect already infected hosts.

##### 3.3.3.3. Infected population

These newly infected individuals will then be added to the infected population. It is these hosts that will then be simulated in the simulation engine for the generation being processed.

##### 3.3.3.4. Simulating infected population

The infected population being infected can be of types. Individuals newly infected or those that are continuing their infection from the previous generation. Apollo is configured to treat both scenarios seamlessly. Each infected host is simulated one at a time. This strategy was selected both to ease computational complexity and help with resource requirements.

##### 3.3.3.5. Per tissue simulation

Within the host, each tissue structure is handled individually at a time. First, it is verified if the tissue being processed has a viral population within it. If it does its population will undergo the replication cycle.

##### 3.3.3.6. Determination of generational phase

Each tissue can have its own independent phase cycle. There can be multiple phases experienced by a single tissue during the course of infection. As mentioned before the generational phase types are neutral, stationary, and depreciation. The user can configure any combination of phases using these three types.

In this step, Apollo determines the generational phase the tissue is currently in given the host's generation in its infection course. Dependent on the phase the parent population will then be adjusted.

##### 3.3.3.7. Infecting cells

The cells are infected based on their affinity to the viral particle and if configured the number of cells available for infection. It is assumed that for each generation a continuous number of cells are available. In the event, that a cell limit is not assigned there is an unlimited number of cells available to satisfy the parent viral population.

##### 3.3.3.8. Determination of parent's progeny

This step is highly parallelized and is conducted on the GPU. Apollo is designed to be adaptable to a multi-GPU architecture. Based on the genome of the parents their fitness is determined and the number of progenies that will be produced by each parent is determined. Then in the presence of recombination, each progeny's recombinant regions will be determined, and the recombinant parents will too be assigned. For this mechanism, the parent viral particles occupying the cell are determined as well as the genomic makeup that caters to the recombination hotspots. Once this is done the total number of progenies that are being generated will be tallied.

##### 3.3.3.9. Generation and configuration of progeny

This too is highly parallelized and GPU dependent. Each progeny's genome is assembled based on their parent's and recombinant parents' genomes. Following this, their mutations are induced while taking into account the proofreading mechanism. The accuracy of the mechanism is influenced by the viral genome. The survivability of each progeny is also determined to estimate those that survive to become parents in the next generation.

##### 3.3.3.10. Cross tissue viral transmission

In the presence of viral transmission between tissues, it will occur after all the viral particles in all tissues have completed their replication cycles. The new progeny will then migrate from one tissue to another based on the user-specified parameters. Each individual transmission event can be configured separately by the user.

##### 3.3.3.11. Sampling of infected hosts

Once all the simulations for all the infected hosts in the given generation have been simulated, we can then sample them given the activation of the sampling mechanism. The sampling is conducted at random of the infected population. Sequences will then be obtained from the sampling tissues and if there are any sampling effects that affect the infectivity of the sampled hosts they will be applied.

##### 3.3.3.12. Configuring the next generation

Once the current generation's simulations are completed. The summary of the simulation will be written, and the simulation will move forward to the next generation. All relevant data structures and prerequisites will be created, and the simulation engine will execute the next generation in the overall simulation. This process will be carried on till a termination event is reached such as the limit of the total number of generations we need the simulation to be carried out for.

### SECTION 4. TESTING AND VALIDATION

To validate Apollo's claims of robustness in its ability to simulate a broad range of epidemic models as well as its efficiency in resource usage we tested it against a series of experimental designs.

Apollo was benchmarked using Compute Canada's (CC) Narval High Performance Computing (HPC) cluster. The configurations consisted of AMD Milan, Intel Gold Skylake, Intel Broadwell or Intel Cascade Lake Central Processing Units (CPU) cores, Nvidia A100SXM4 or NVIDIA v100SXM2 Graphical Processing Units (GPU) with Solid State Device (SSD) including NVMe SSD storage. These benchmark configurations are subject to change and in these instances, they will be specifically stated.

We begin with a series of WF simulations such as the theory of fixation[49]. This enabled us to test the implementation of Apollo's foundational theories upon which it was built. Apollo by design in its default state follows the standard WF model complete with its assumptions except for Apollo being a haploid model.

Next, we introduced neutral mutations into our WF model simulations. This enabled the testing and validation of Apollo's mutation mechanic. We will then be able to validate protocols involved with the mutation mechanism such as the clock and site models and the configuration of mutational hotspots.

#### 4.1. Benchmarking: Software processing speeds and efficiency

In order to evaluate Apollo's processing capabilities, it was tested against a series of varying scenarios across a broad range of systems.

For the baseline, we selected the configurations of Compute Canada's Beluga cluster. Beluga comprises of a NVMe SSD storage system with NVIDIA V100SXM2 GPUs and Intel Gold 6148 Skylake CPUs. We used 20 CPU cores, followed by 50GB of RAM. The number of genomic data loaded to the GPU at a time was set to 100000, and the number of cells being processed at a time was set to 10000. The file management system of Apollo created intermediary files was set to bin size of 10000 entries at maximum.

For our benchmarks, we estimated the per generation processing time of Apollo under varying population sizes. The selected sequences spanned 701 bases and simulations were tested against stationary population sizes. The selected ten population sizes were 10000, 20000, 30000, 40000, 50000, 60000, 70000, 80000, 90000 and 100000. We measured the time taken by Apollo to process a generation under each population under various test conditions. Measurements were taken from 100 generations for each test condition to evaluate the average processing times.

Our initial testing was for Apollo to process the different population sizes under control scenarios where no mutation, recombination, or selection mechanisms were in effect. Then we tested Apollo's processing capabilities with the inclusion of these mechanics. Finally, we tested how Apollo's processing times vary with different hardware.

#### 4.1.1. Estimating the baseline processing times and its rate of change with varying population sizes

To get a baseline control assessment of Apollo's speed and efficiency it was tested on Beluga from Compute Canda. Under these hardware conditions, we obtained the average time taken to process different population sizes. The results from the tests are depicted in **Figure S - 13**.

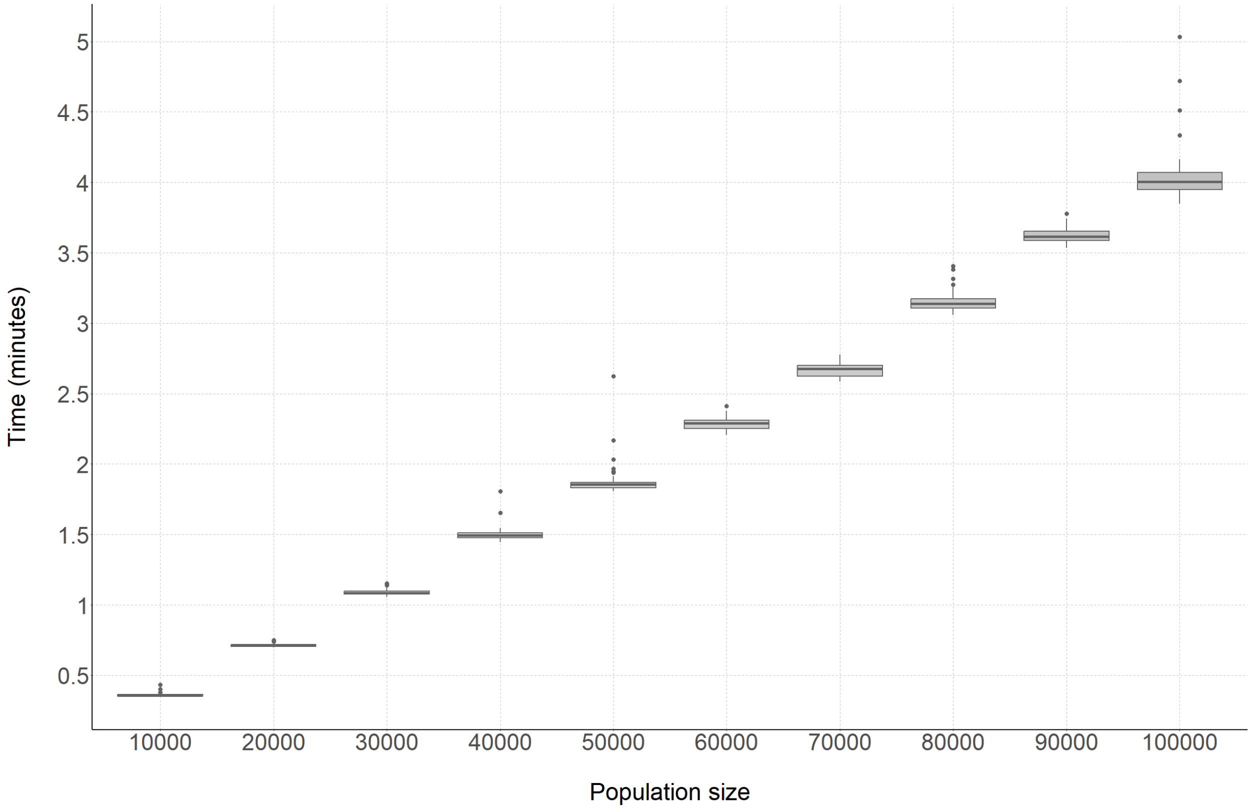

**Figure S - 13.** Boxplot of the runtimes for each population size for Apollo conducted using the Compute Canada, Beluga cluster. We can see a linear increase in the average per generation processing time with the increasing bin size.

Apollo shows a linear increase in the processing times proportional to the increase in population size. We observed on average a 1.3297 minute increase in per generation processing time for every 10000 step increase in population size.

This result will act as our baseline to compare the rest of our analyses on Apollo's processing times.

#### 4.1.2. Benchmarking processing times for different mechanisms in Apollo that govern variation

Here we evaluated the change in processing times due to the mechanisms of mutation and recombination. Apollo was tested against the presence of either mutation or recombination followed by the activation of both mechanisms. We selected 192 mutational hotspots and 14 recombination hotspots. The results are depicted in **Figure S - 14**.

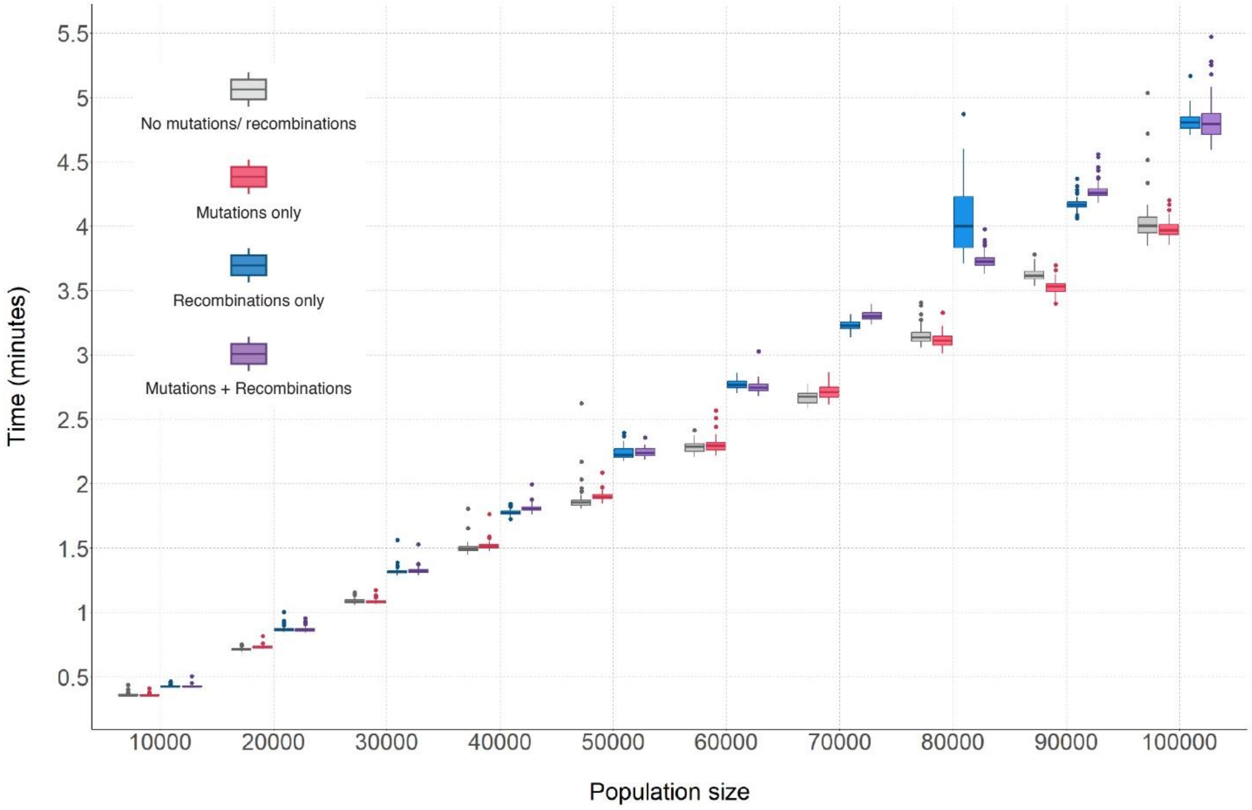

**Figure S - 14.** Boxplot graph comparing the effects of mutation and recombination mechanisms on runtime. As observed the mutation and recombination mechanisms cause an increase in the per generation processing time. The mutation mechanism (red) appears to have a lesser impact on the baseline processing time (grey) compared to the recombination mechanism (blue). When both mechanisms are activated (purple) as expected there is an increase in the overall processing time, but only by a few seconds.

As observed in **Figure S - 14** we can see that the variation mechanisms cause an increase in the overall per generation processing times. Mutations cause an average increase of about 0.997 times, recombinations cause a 1.205 increase in processing time, and mutations and recombinations together cause an average overall increase of 1.201.

#### 4.1.3. Benchmarking processing times under different hardware

To evaluate the capability of Apollo to adapt to the resources available in different hardware we tested it against the A100 GPU (**Figure S - 15**). The A100 is known to have a markable improvement over the V100 in terms of performance with a larger number of CUDA cores and the ability to process at 19.5 teraflops while the V100 stands at 14 teraflops.

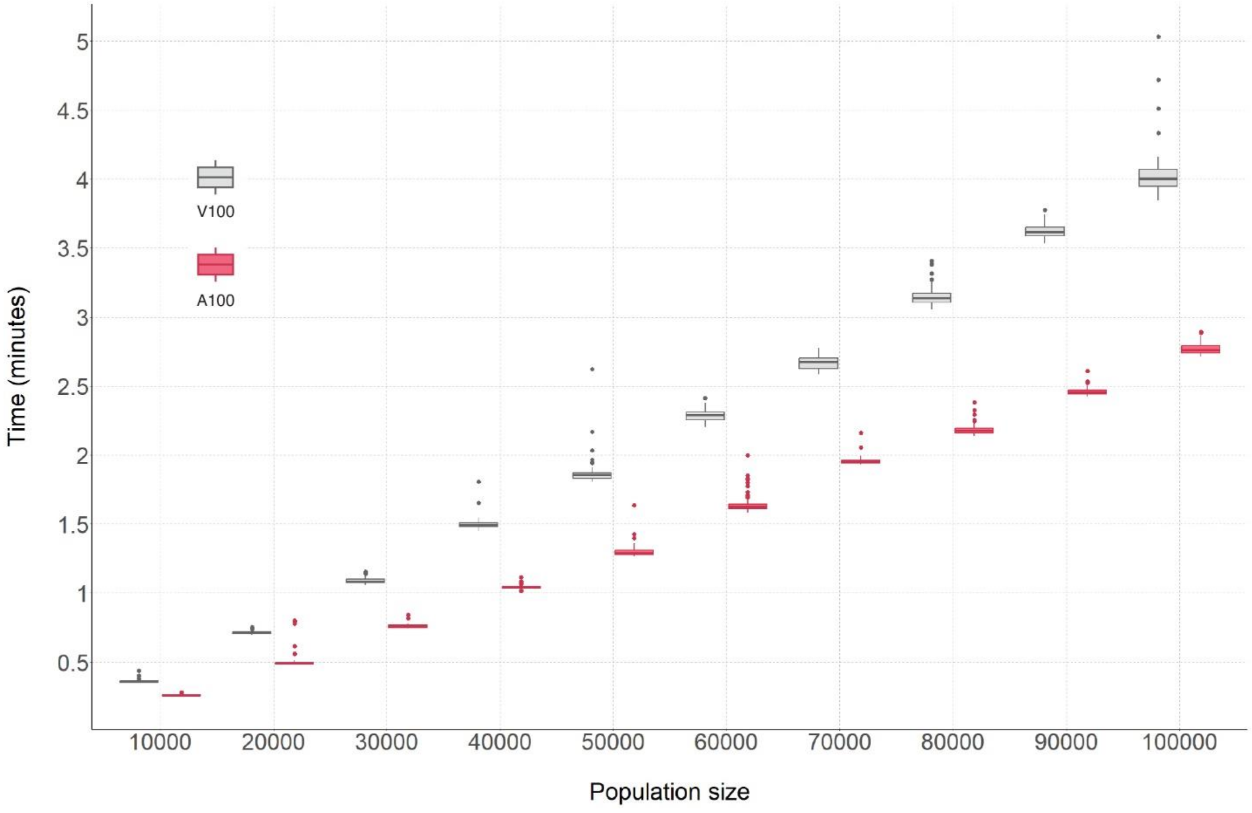

**Figure S - 15.** Comparative analysis of the performance change of Apollo with V100 (grey) vs A100 (red) GPU architectures. We can observe a marked increase in Apollo's performance under the A100 GPU.

As expected, there was a marked increase in Apollo's performance when using the more powerful A100 architecture. Apollo had on average 1.4 times faster per generation processing time compared to the V100. This clearly shows Apollo's adaptability to different hardware platforms and also strengthens its claim to be scalable and adaptable to future parallel processing architecture. We can expect faster processing time from Apollos under more robust and powerful computer hardware.

#### 4.2. Standard Wright Fisher simulations

For the fundamental WF simulations conducted to test Apollo's foundational framework, two tests were carried out. In both instances five of the six WF assumptions were maintained, the exception being the haploid genomes. Therefore, these tests allow us to understand the change in allele frequency based on the effects of genetic drift alone. Under the principles of genetic drift, the current population is sampled at random to determine the next generation. At the genomic level, this act of random sampling causes deviations in the genetic composition of each successive generation. The cumulative effect of these deviations results in loss of genetic variation over time followed by the fixation of alleles. Once fixation occurs all individuals in the entire population contain the same genetic makeup.

##### 4.2.1. Experimental design and results

The simulations were designed with a single node who had only a single tissue compartment. The genomes were devoid of mutational or recombination hotspots. All allele positions regardless of their occupying base contributed equally towards fitness, survivability, and proofreading (which was deactivated since there were no mutations). All genomes would have the same reference values regardless of their base composition. The population size within the node was set to be constant. These are the default parameters of Apollo hence it should follow the WF model. Therefore, we would expect to observe fixation of a particular genome (or haplotype) in the within host population as the simulation progresses through time.

In the first experiment, only two different genomes were present. The genomes were of size one base in length and were labeled Allele A and Allele B. Each genome had a population of 100 viral particles at the start; therefore, the preliminary frequency was 0.5 for each. The rate of progeny generation was based on a negative binomial distribution of $n = 10$ and $p = 0.55$. The simulation was run for 500 generations.

The subsequent experiment expands on the scope of the previous experiment. Instead of two distinct genomes, 100 unique haplotypes were created for a population size of 1000 where each haplotype had 10 copies each for an individual haplotype frequency of 0.01 at the start. The simulation was run for 2818 generations. The results of both tests are depicted in **Figure S - 16**.

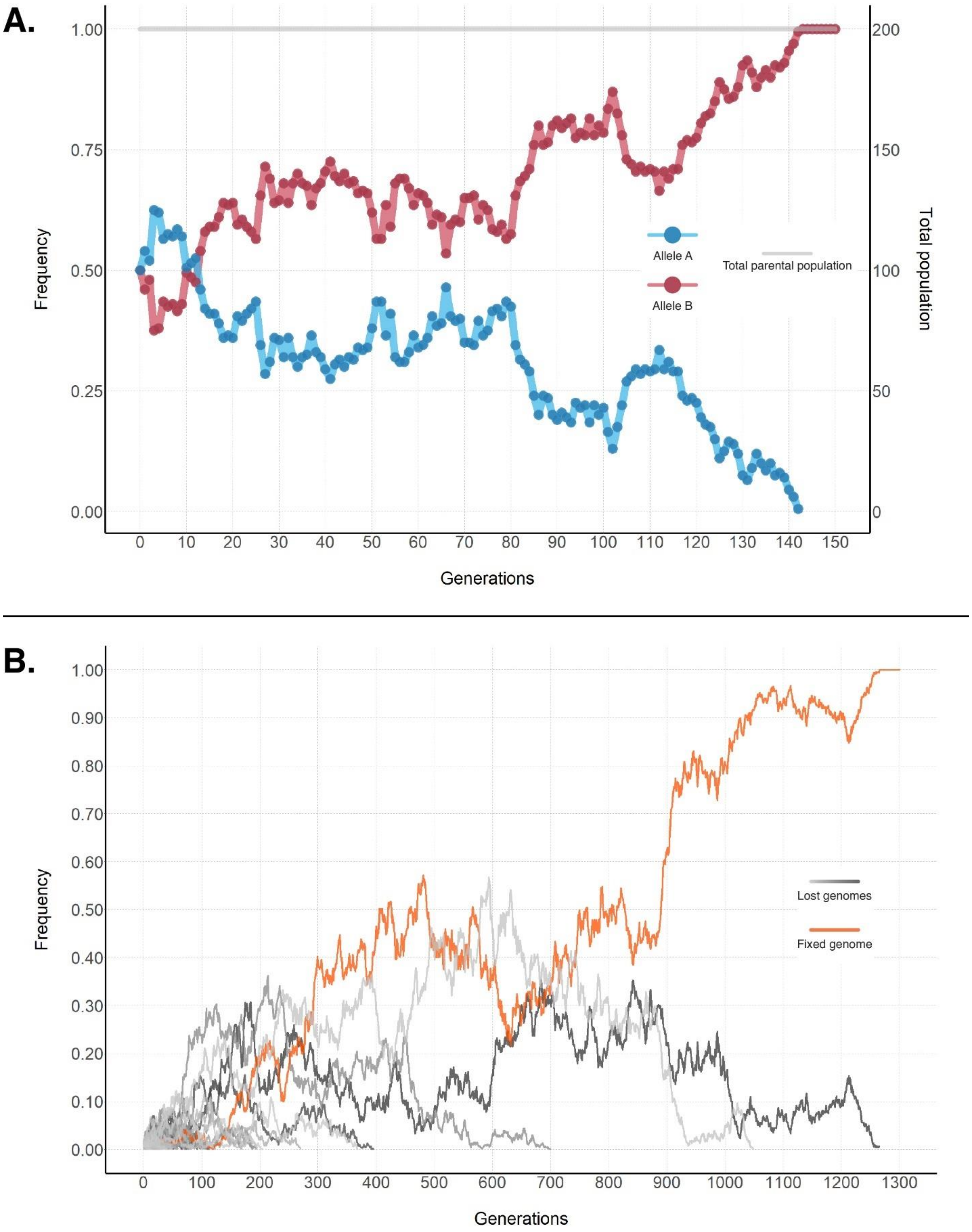

**Figure S - 16.** Simulations under the standard Wright Fisher model for haploid viral genomes. The generations are abbreviated to show up to fixation (A) The changes in the two alleles' frequencies in the population across successive generations are shown. Allele A (red) reaches fixation while allele B (blue) is lost. The total parental population (grey) is constant across generations. (B) The changes in the frequency of 100 haplotypes. One genome reaches fixation (orange) while the others are lost at varying time points.

Data analysis was conducted with the aid of Apollo's utility tools such as "Haplotype Counter". The Haplotype Retriever was able to detect the unique haplotypes in each successive generation as well as their frequencies and counts.

Under Apollo's default parameters, we can see that the Wright-Fisher model is being adhered to (**Figure S - 16**). In all instances regardless of population size or the number of unique haplotypes under the forces of genetic drift alone fixation did occur. Apollo maintained a constant population size across the successive generations (**Figure S - 16A**) and the population was sampled at random for the next generation.

#### 4.3. Wright Fisher with neutral mutations

Under the WF model mutations can be introduced while maintaining the prospect of neutrality. Under this neutral model, these mutations provide no benefit or disadvantage relative to the remaining population. Additionally, if we remove the possibility of back mutations, we can intuitively predict that these new mutations should reach fixation.

##### 4.3.1. Experimental design and results

To validate Apollo based on the above stated hypothesis we ran simulations similar to that in Section 4.2.1's first experimental setup but now with the introduction of mutations. To this end, a mutational hotspot was added for the single locus with a mutation rate that follows a Poisson distribution with $mean\ (\mu) = 0.01$ and the starting population was increased to 1000 viral particles. Two experimental designs were set up under the aforementioned conditions, and both were run for 5000 generations.

In the first instance, we started with an equal number of viral particles for two haplotypes (**Figure S - 17**). The haplotypes were of base A and base T. They would each have the ability of mutating to base C with a probability of 0.5 to produce a new haplotype, or not change their current base with a probability of 0.5 (**Figure S - 17 A and B**). The mutated haplotype of base C cannot undergo mutation to change its base. All haplotypes are neutral to each other and have no selective advantage. Therefore, with time eventually, haplotype C should be fixed in the population.

In our second instance, we expanded on our previous experiment. The haplotype A was configured to produce only mutated haplotypes of base G in addition to itself with base change probabilities of 0.5 for both. Similarly, haplotype T would only produce mutated haplotypes of base C. Both G and C base haplotypes would not mutate (**Figure S - 18 A and B**). Therefore, under the effects of genetic drift one of the mutated haplotypes should reach fixation and their frequency in the population should be affected by their original haplotype.

In our first experiment as expected, there was a dramatic increase in the mutated haplotype (Allele C) as both the original haplotypes (Allele A and Allele T) would produce it with a 50 percent probability during a mutation event (**Figure S - 17A and C**) while the mutated form produced a

100 percent of its own genome. It was observed that the haplotype with allele T would first become extinct followed by the haplotype with allele A. In this manner, both original haplotypes disappear from the population leaving only the mutated haplotype even though its mutation is neutral (**Figure S - 17C**).

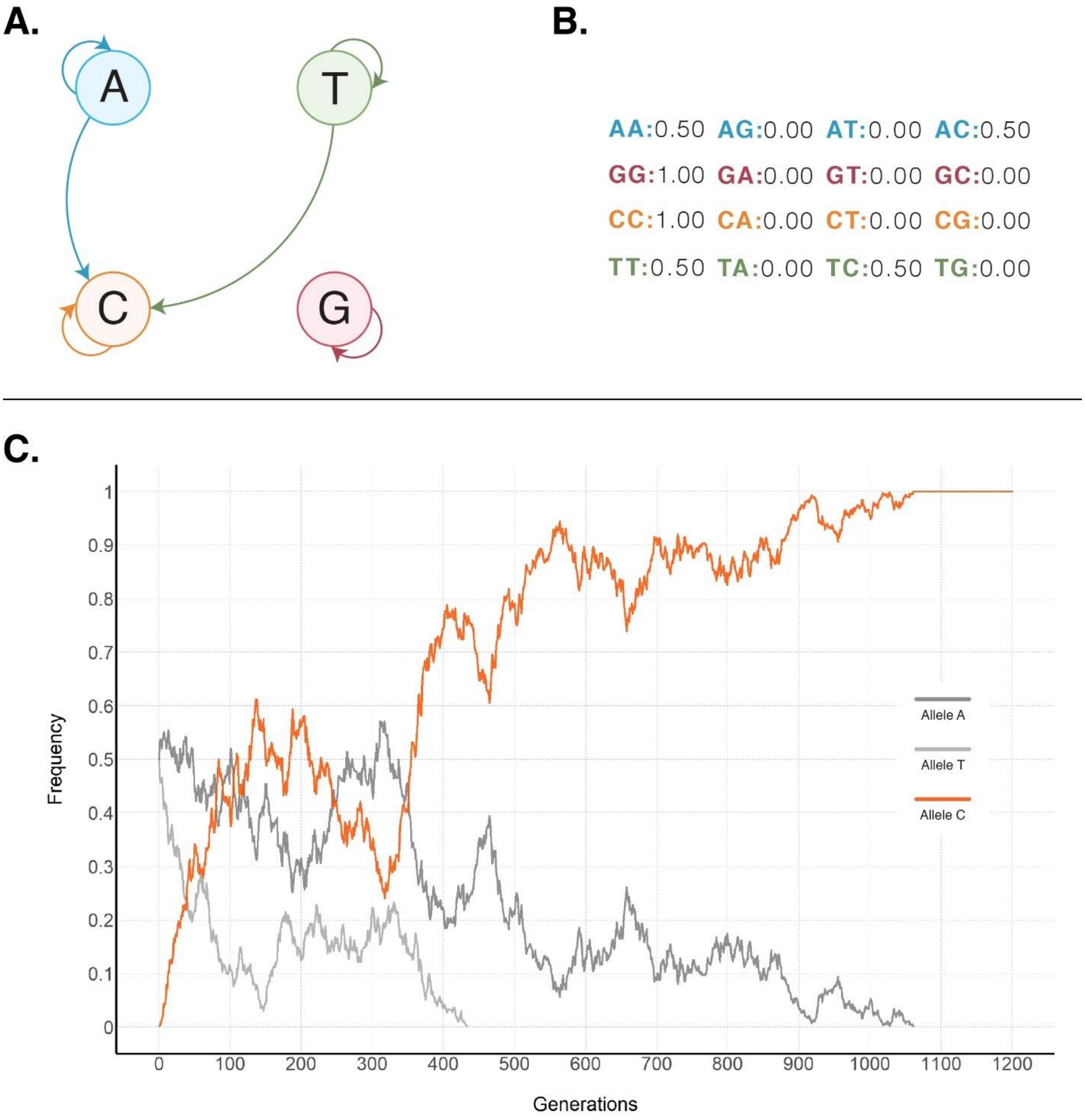

**Figure S - 17.** Experimental setup and results of the first simulation for Wright Fisher model with neutral mutations. (A) Shows the Markov chain for the site model and (B) shows how the site model is configured within Apollo. As shown even though base G (Guanine) is not involved in the simulation it still has to be configured. (C) The results of the simulation show the variation in the frequencies of the different haplotypes. As observed the mutated haplotype containing Allele C (orange) reaches fixation eventually, while the original haplotypes (Allele A in dark grey and Allele T in light grey) cease to exist in the population.

The above test ran for a total of 5000 generations in 1.99 hours. A total of 40,907,149 viral particles were generated which means on average 5714 viral particles were generated per second.

To further validate our claims, we performed an additional test (**Figure S - 18**). Here the two original haplotypes produced two separate mutated haplotypes. As expected initially the mutated haplotypes saw an exponential increase in the population due to the original haplotypes producing them with a 50 percent probability during a mutation event (**Figure S - 18B and C**). Akin to the previous experiment the mutated forms did not undergo back mutation. We can see that eventually, the mutated haplotype containing Allele G reached fixation in the population (**Figure S - 18C**). However, a series of interesting trends can also be observed before the fixation event. From the original haplotypes, the haplotype with Allele A is persistent long after the Allele T haplotype. This allows it to boost the frequency of Allele A haplotype. The population of the mutated haplotype with allele C drops in comparison to that of the mutated form with allele G since haplotype T has disappeared. Nonetheless, the mutated haplotype with allele C persists longer than the haplotype with allele A since the original haplotype loses progeny that mutate to form allele G haplotypes (**Figure S - 18C**).

The above test ran for a total of 5000 generations in 1.98 hours. A total of 40,915,622 viral particles were generated which means on average 5734 viral particles were generated per second.

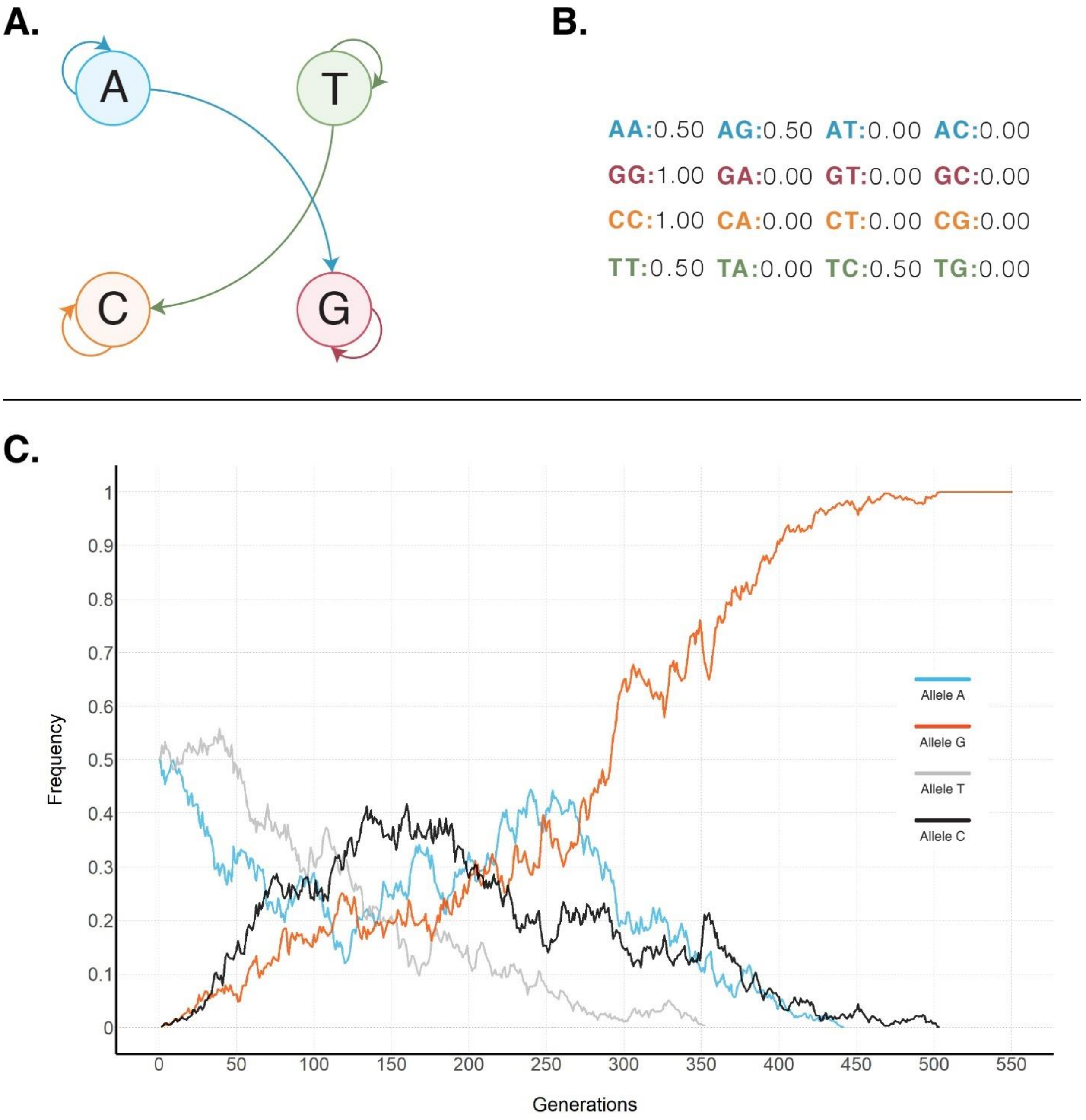

**Figure S - 18.** Experimental setup and results of the first simulation for Wright Fisher model with neutral mutations resulting in two mutated haplotypes. (A) Shows the Markov chain for the site model and (B) shows how the site model is configured within Apollo. (C) The results of the simulation show the variation in the frequencies of the different haplotypes. As observed the mutated haplotype containing Allele G eventually reaches fixation, while the original haplotypes A and T cease to exist in the population followed by the other mutated haplotype C.

This loss in haplotypes and the resultant fixation occurs due to the lack of back mutations from the mutated form. This proves the validity of Apollo's mutation mechanism in its ability to simulate neutral mutations and the changes in a population's diversity under the effects of genetic drift alone.

#### 4.4. Wright Fisher model with selection

Apollo can accommodate selection forces to govern the evolutionary path of a virus in an epidemic. To show how the forces of selection can affect the virus in an epidemic we designed the following experiment. We started with a single loci genome with base A. It would mutate to form either itself or a mutated form with base T with a probability of 0.5 each (**Figure S - 19**). The mutational hotspot was added for the single locus with a mutation rate that follows a Poisson distribution with $mean\ (\mu) = 0.01$ and the starting population was increased to 1000 viral genomes of the original base A haplotype. And the simulation was conducted for 5000 generations.

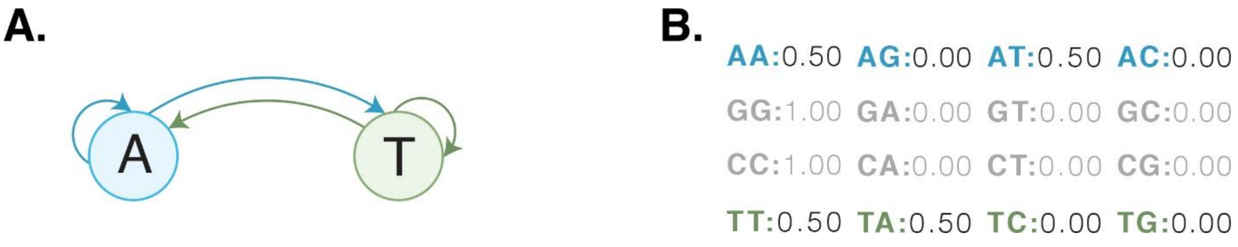

**Figure S - 19:** (A) Shows the Markov chain for the site model and (B) shows how the site model is configured within Apollo. As shown even though base G (Guanine) and C (Cytosine) are not involved in the simulation they still have to be configured.

Under the neutral theory and the site model, the mutated allele T in the population should rise eventually to a frequency of about 0.5 and fluctuate within its range in the absence of any form of selection.

To show the effects of selection, we can repeat the previous experiment but introduce a selection force. This was done in terms of survivability. The probability of a progeny surviving to the next generation for the mutated haplotype (Allele T) was reduced from 100 percent to 85 percent. The original haplotype's (Allele A) probability of survival remained at 100 percent. In this instance, we should observe a fluctuation in the distribution of two haplotypes across generations when compared to the first experiment. Intuitively we can understand that the mutated haplotype frequency should consistently be below that of the original haplotype. After the experiments, we observe our expected patterns for the frequencies of the haplotypes (**Figure S - 20**).

A.

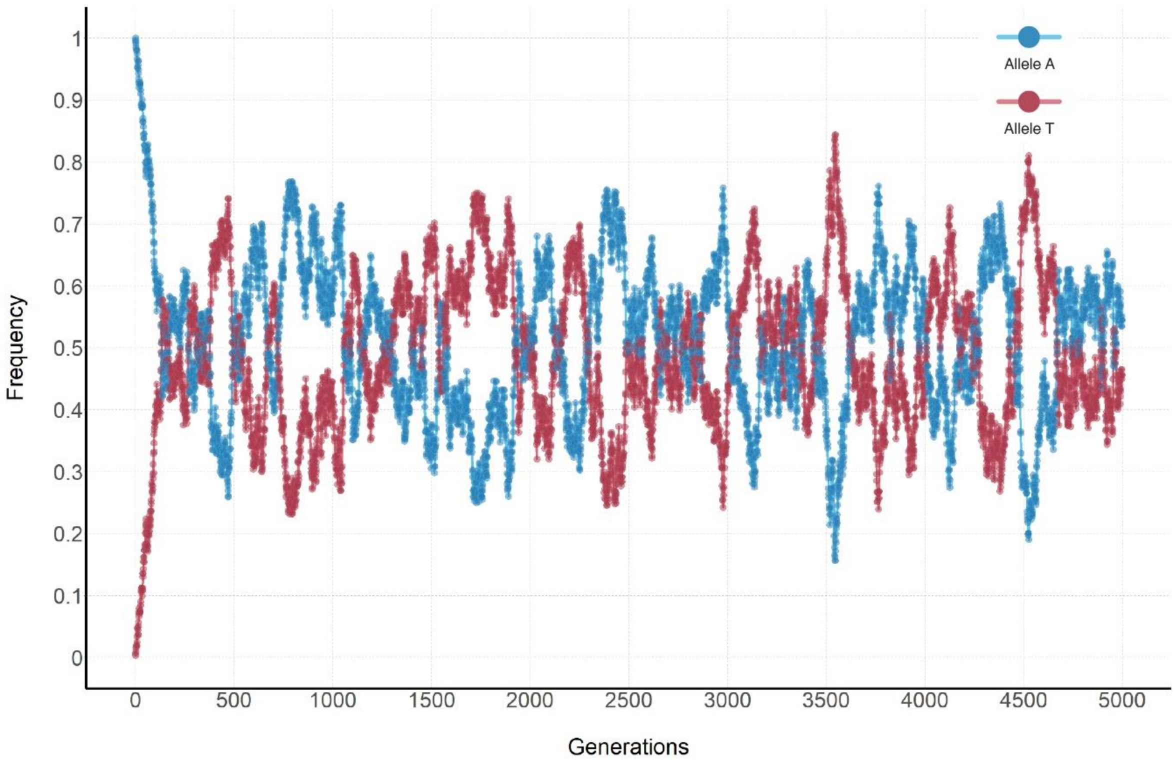

B.

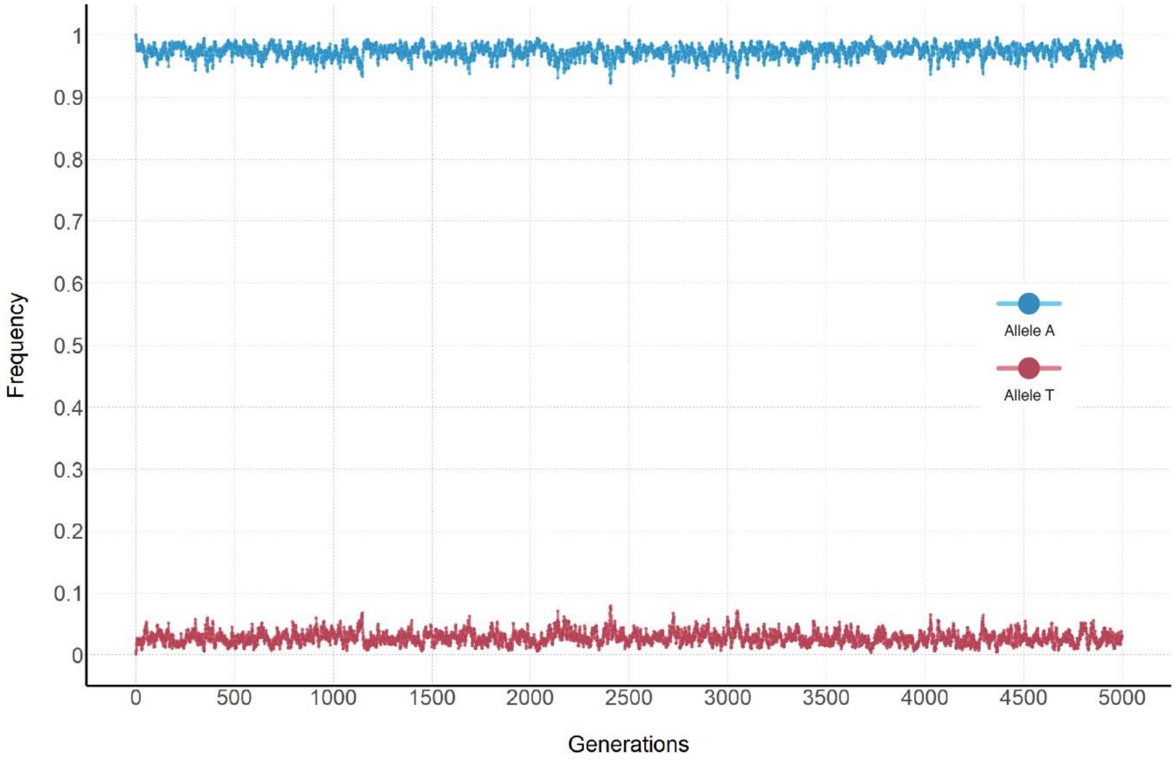

**Figure S - 20:** The change in haplotype frequencies in the presence and absence of selection forces. (A) The change in haplotype frequencies in the absence of selection forces. As observed the mutated haplotype's frequency quickly rises to meet that of the original haplotype. They fluctuate around the 0.5 frequency value which matches the site model governing the simulation. (B) Shows the change in haplotype frequencies in the presence of selection. As expected even though the mutated haplotype does appear in the population it exists at a much lower frequency due to the negative selection pressure.

Even though the survivability changes by only 15% there is a significant drop in the allele frequency of the mutated haplotype in comparison to when it evolves under neutrality. This shows that Apollo's selection forces under genomic variation work as intended including its base substitution site models.

##### 4.5. Simulating quasispecies

The theory of the Quasispecies principle is used to explain the diverse viral populations that have been found to exist within a host once the mutation-selection balance has been achieved. To achieve quasispecies a genome must experience a high rate of mutation usually caused by replication errors. This elevated genomic diversity is one of the primary candidates that contribute towards the evolution of viruses to overcome drug therapeutics and the host immune response.

To showcase Apollo's ability to simulate quasispecies we have designed a simple experiment. The viral population's genome is composed of a single base, and it can take the form of any one of the bases A, T, G, and C. In our example, base A does not undergo mutation, while T, G, and C do undergo mutation. They will produce progeny of themselves with a probability of being unchanged during a mutation event with a probability of 0.5 and will change the base with a probability of 0.25 each for either one of the other two bases (**Figure S - 21A** and **B**).

Taking the probability of mutation as our fitness landscape we can generate the matrix $W$ as follows:

$$\mathrm{W} = \begin{bmatrix} 1 & 0 & 0 & 0 \\ 0 & 0.5 & 0.25 & 0.25 \\ 0 & 0.25 & 0.5 & 0.25 \\ 0 & 0.25 & 0.25 & 0.5 \end{bmatrix}$$

Solving for the matrix the eigenvalues and their respective eigenvectors are as depicted in online methods **Table 1**. From the results, it can be observed that only two quasispecies states could be viable in the system. This being those with eigen values of 1 and nonnegative values in the eigenvector. It can also be observed that these two combinations represent the scenario where only the viral particles with allele A being fixed in the system while the rest have become extinct and the other being that allele A has become exist and the other sequences will exist in the system together.

Though individually the remaining alleles of T, G, and C are of lesser fitness than A due to the fact that they mutate to produce less of themselves, while A only produces its own progeny, the quasispecies of T, G, and C have an equal fitness to that of A. This is represented by the equivalent eigenvalue for both in online methods **Table 1**.

Therefore, in our simulation, we expect to observe a fixation of either T, G, and C in the population with A becoming extinct or A becoming fixed and the former quasispecies system being extinct. However, due to the ability of T, G, and C to be able to produce progeny of each other and given a mutation rate that is not too low, it will be more likely that A will become extinct in the population. We set up our experiment with the same mutation rate as that of the previous experiments followed by a run of 5000 generations.

This is the phenomenon we observed during our simulation as depicted in **Figure S - 21**. Once the selection mutation balance is achieved the quasispecies of T, G, and C would exist in the population together. We observed that allele A became extinct in the population.

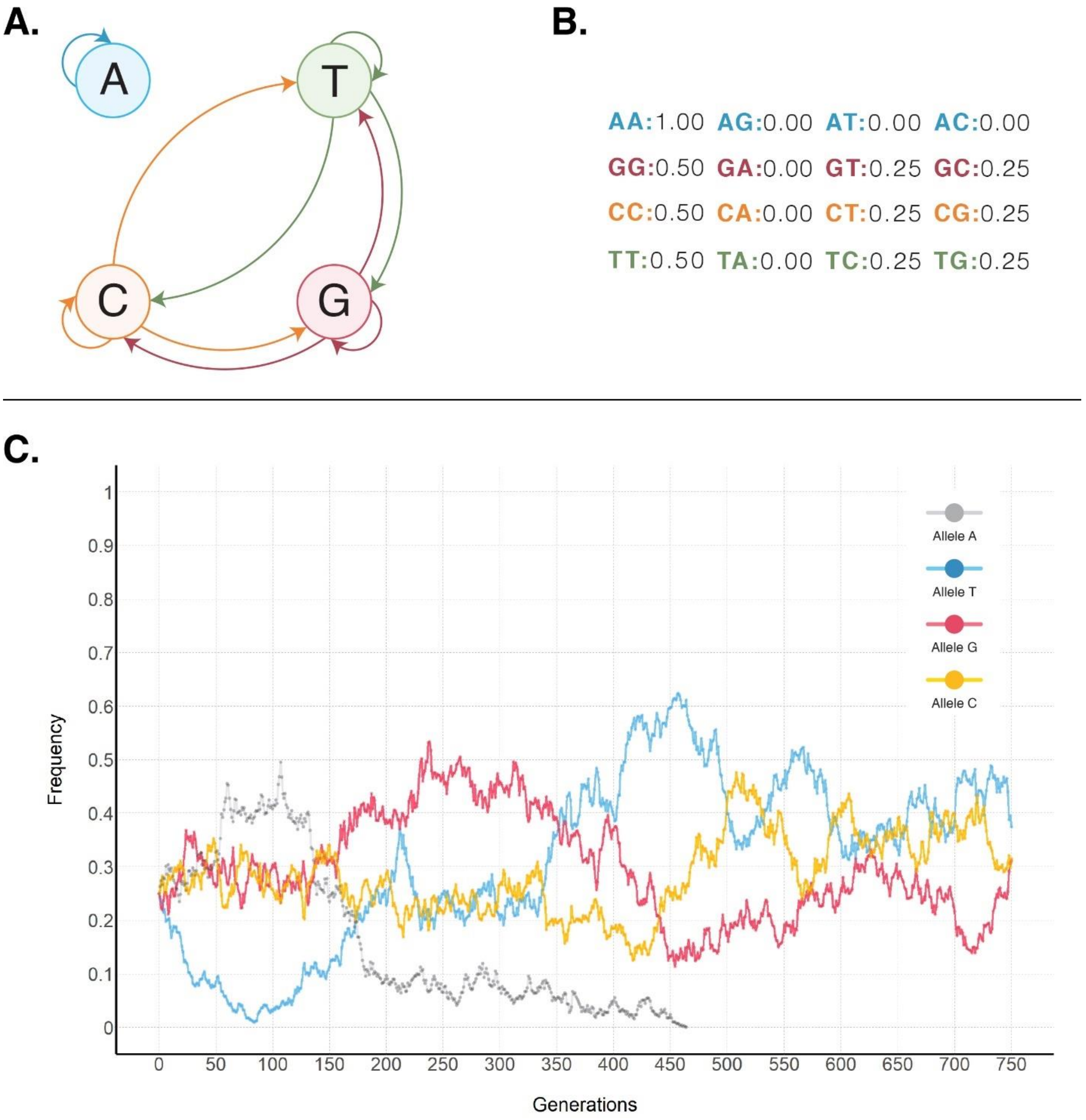

**Figure S - 21.** Depiction of the experimental setup for the site model and results for the quasispecies experiment. (A) The configured site model for base changes during mutation and (B) the Apollo's site model format with the base transition probabilities for the site model. (C) A snippet of the simulation with the results of the change in frequency of each variant. As shown variants with allele A have become extinct with time, while variants with alleles T, G, and C have formed a quasispecies mutation selection balance and continue to persist in the population.

#### 4.6. Simulation of real-world data

To assess Apollo's proficiency in simulating within host dynamics in relation to viral infection we attempted to replicate the infection cycle observed in an individual infected with HIV. Using Apollo, we conducted an analysis with the goal of emulating the progression of HIV and its infection within a host. Subsequently, we validated Apollo's fidelity by juxtaposing its simulated data against the real-world clinical results acquired through the course of the individual's infection.

##### 4.6.1. Experimental goals and setup

The goal of the real-world simulations was to exhibit Apollo's ability to replicate real world clinical data through simulations alone. This endeavor has a number of promising implications in the study of epidemiology.

Successful simulation of viral sequences would allow researchers to validate parameters in relation to viral mutation rates, site models, recombination, and phenotypic information. It would also allow the understanding of the viral cloud that exists within hosts and the effects of reservoirs and on the viral population and disease progression[17,38,71,72]. The test was therefore broken into two phases.

The first phase involved only the PBMC tissue data of the individual between the first two visits. The data points span from the 11$^{th}$ of May 1993 to the 14$^{th}$ of September 1993. This is a time frame of 126 days or four months and four days including the first and last days. We would estimate Apollo's ability to simulate the sequences extracted at the second time point given only the first time point's sequences. The initial focus on the first four months would enable us to simplify the experimental design and identity and rectify any discrepancies before moving onto the more complicated complete simulation spanning the total period of 882 days.

The second phase would comprise the entirety of the collected real world clinical data. We would attempt to use Apollo to simulate the entirety of the individual's infection cycle, across all five tissues ranging from the first visit to their last visit. This is a span of 882 days or two years four months and 30 days. We would then attempt to recover the sequences that were found during each visit from Apollo's simulations.

These tests would allow us to evaluate the information that Apollo produces as well as its fidelity and resource efficiency. We will also be able to estimate if Apollo's simulations can be used to further the insights that were obtained via the clinical analyses.

##### 4.6.2. Details on real world clinical data

Real world clinical data of an HIV infected individual was provided in the form of viral sequences and temporal information on sampling by the Southern Alberta Clinic in Calgary, Alberta, Canada[20]. The individual was infected with HIV and was undergoing Anti-Retroviral Therapy (ART)[73–76]. Their ART comprised of monotherapy using Didanosine (DDI)[77–80]. The region of the HIV genome that was sequenced consisted of an area spanning 701 bases from the Reverse Transcriptase section of polymerase (pol) region[81–83]. The individual was sampled across a period of three years through seven visits (**Online Methods Table 2**). Samples were obtained from five different tissues, namely, Peripheral Blood Mononuclear Cells (PBMC), colon, stomach, duodenum, and esophagus[84–87].

##### 4.6.3. Analysis of the sequenced region

The sequences were obtained using first generation Sanger sequencing. Using NCBI's Basic Local Alignment Search Tool (BLAST) algorithm[88–90] we identified the section of the HIV genome's RT pol region the sequences were obtained from (**Figure S - 22**).

Using the BLAST results (**Figure S - 22**) we were able to confirm the exact region the sequence was obtained from. The BLAST matched our query region with a complete genome of HIV-1 which was a consensus sequence (GenBank: MN919177.1)[91].

On identifying the region of the genome that was being sequenced, we were able to obtain information regarding its evolutionary and mutational dynamics (**Figure S - 23**). As shown the target region to be analysed consisted of 701 base pairs. Typically, in the HIV genome, the p51 Reverse Transcriptase (RT) is found within the pol region. It starts from base position 2550 and extends to base 3870. The sequences obtained from the infected individual are nestled within this region as depicted in **Figure S - 23**.

Synthetic construct strain HIV-1 type 1b consensus, complete sequence

Sequence ID: MN919177.1 Length: 9719 Number of Matches: 1

**Range 1: 2550 to 3250** GenBank Graphics

| Score | Expect | Identities | Gaps | Strand |
|---|---|---|---|---|
| 1201 bits(650) | 0.0 | 684/701(98%) | 0/701(0%) | Plus/Plus |

```
Query  1     CCCATTAGCCCTATTGAGACTGTACCAGTAAAATTAAAGCCAGGAATGGATGGCCCAAAA  60
             ||||||||||||||||||||||||||||||||||||||||||||||||||||||||||||
Sbjct  2550  CCCATTAGCCCTATTGAGACTGTACCAGTAAAATTAAAGCCAGGAATGGATGGCCCAAAA  2609

Query  61    GTTAAACAATGGCCATTGACAGAAgaaaaaataaaagcattagtagaaatttgtacagaa  120
             ||||||||||||||||||||||||||||||||||||||||||||||||||||||||||| 
Sbjct  2610  GTTAAACAATGGCCATTGACAGAAGAAAAAATAAAAGCATTAGTAGAAATTTGTACAGAG  2669

Query  121   atggaaaaggaagggaaaatttcaaaaattGGGCCTGAAAATCCATACAATACTCCAGTA  180
             ||||||||||||||||||||||||||||||||||||||||||||||||||||||||||||
Sbjct  2670  ATGGAAAAGGAAGGGAAAATTTCAAAAATTGGGCCTGAAAATCCATACAATACTCCAGTA  2729

Query  181   TTTGCCATAAAGAAAAAAGACAGCACTAGATGGAGAAAATTGGTAGATTTCAGAGAACTT  240
             ||||||||||||||||||||||| |||| |||||||||||| ||||||||||||||||||
Sbjct  2730  TTTGCCATAAAGAAAAAAGACAGTACTAAATGGAGAAAATTAGTAGATTTCAGAGAACTT  2789

Query  241   AATAAAAGAACTCAAGACTTCTGGGAAGTTCAATTAGGAATACCACATCCCGCAGGGTTa  300
             ||||| ||||||||||||||||||||||||||||||||||||||||||||||||||||||
Sbjct  2790  AATAAGAGAACTCAAGACTTCTGGGAAGTTCAATTAGGAATACCACATCCCGCAGGGTTA  2849

Query  301   aaaaagaaaaaaTCAGTAACAGTACTGGATGTGGGTGATGCATATTTTTCAGTGCCATTA  360
             ||||||||||||||||||||||||||||||||||||||||||||||||||||| || |||
Sbjct  2850  AAAAAGAAAAAATCAGTAACAGTACTGGATGTGGGTGATGCATATTTTTCAGTTCCCTTA  2909

Query  361   GATAAAGAATTCAGGAAGTATACTGCATTTACCATACCTAGTATAAACAATGAGACACCA  420
             |||||||| |||||||||||||||||||||||||||||||||||||||||||||||||||
Sbjct  2910  GATAAAGACTTCAGGAAGTATACTGCATTTACCATACCTAGTATAAACAATGAGACACCA  2969

Query  421   GGGATTAGATATCAGTACAATGTGCTGCCACAGGGATGGAAAGGATCACCAGCAATATTT  480
             |||||||||||||||||||||||||| |||||||||||||||||||||||||||||||| 
Sbjct  2970  GGGATTAGATATCAGTACAATGTGCTTCCACAGGGATGGAAAGGATCACCAGCAATATTC  3029

Query  481   CAGAGTAGCATGACAAGAATCTTAGAGCCTTTTAGAAAACAAAATCCAGAAATAGTCATC  540
             || ||||||||||||| ||||||||||||||||||||||||||||||||| ||||| |||
Sbjct  3030  CAAAGTAGCATGACAAAAATCTTAGAGCCTTTTAGAAAACAAAATCCAGACATAGTTATC  3089

Query  541   TATCAATACATGGATGATTTGTATGTAGGATCTGACTTAGAAATAGGGCAGCATAGAATA  600
             |||||||||||||||||||||||||||||||||||||||||||||||||||||||||| |
Sbjct  3090  TATCAATACATGGATGATTTGTATGTAGGATCTGACTTAGAAATAGGGCAGCATAGAACA  3149

Query  601   AAAATAGAGGAACTGAGACAACATCTGTTGAAGTGGGGATTTACCACACCAGACAAAAAA  660
             ||||||||||| ||||||||||||||||||| ||||||||||||||||||||||||||||
Sbjct  3150  AAAATAGAGGAGCTGAGACAACATCTGTTGAGGTGGGGATTTACCACACCAGACAAAAAA  3209

Query  661   CATCAGAAAGAACCTCCATTCCTTTGGATGGGTTATGAACT  701
             |||||||||||||||||||||||||||||||||||||||||
Sbjct  3210  CATCAGAAAGAACCTCCATTCCTTTGGATGGGTTATGAACT  3250
```

**Figure S - 22.** NCBI BLAST results showing the region of the HIV genome from which the sequences were obtained from.

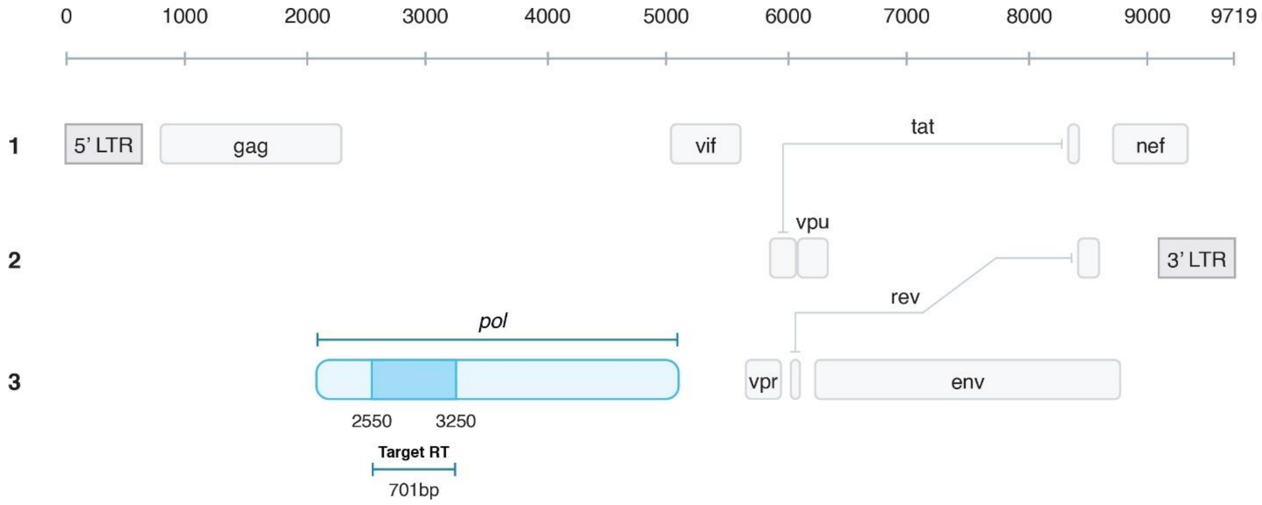

**Figure S - 23.** Detailed structure of the HIV genome. The target region as shown (dark blue) spans a region of 701 bases of the polymerase (pol) region (light blue).

##### 4.6.3.1. Identification of recombination hotspots

Through a study conducted by Smyth *et al* in 2014, we were able to identify 14 recombination hotspots in the target region[92]. The average recombination events per nucleotide per round of infection (REPN) was selected to be $1.8 \times 10^{-3}$. The coordinates and subsequent recombination rates of each hotspot were calculated and configured into Apollo's parameters[92] (**Table S - 1**).

**Table S - 1.** Details of the 14 recombination hotspots. Their start and stop coordinates as positioned based on the complete HIV genome and the calculated rates of recombination.

| Hotspot ID | Start coordinates | Start coordinates | Rate of Recombination |
|---|---|---|---|
| 1 | 2573 | 2615 | 0.07818 |
| 2 | 2615 | 2651 | 0.06660 |
| 3 | 2651 | 2681 | 0.05580 |
| 4 | 2681 | 2726 | 0.08280 |
| 5 | 2726 | 2771 | 0.08280 |
| 6 | 2771 | 2825 | 0.09900 |
| 7 | 2825 | 2870 | 0.08217 |
| 8 | 2870 | 2909 | 0.07200 |
| 9 | 2909 | 2966 | 0.10440 |
| 10 | 2966 | 3011 | 0.08226 |
| 11 | 3011 | 3065 | 0.09900 |
| 12 | 3065 | 3116 | 0.09360 |
| 13 | 3116 | 3167 | 0.09360 |
| 14 | 3167 | 3218 | 0.09417 |

##### 4.6.3.2. Determination of mutational hotspots

To determine the mutational hotspots, we used a Multiple Sequence Alignment (MSA) strategy using MUSCLE (MUltiple Sequence Comparison by Log-Expectation) alignment. The MEGA (Molecular Evolutionary Genetic Analysis) software was used[93,94].

For phase one we collected the sequence data from the two time points from only PBMC data. Then using MSA we identified 30 different segregating sites. These are our mutation hotspot sites. Using literature, we estimated the mutation rate to on average follow a Poisson distribution of mean 0.33333 per replication cycle[95,96]. The base substitution transition matrix for each segregating site was then determined by careful analysis of the MSA data. For phase two we used all the sequence information across all seven time points from all five tissues. In this instance, a total of 192 segregating sites were identified.

#### 4.6.4. Within tissue Phases of infection

The phases of the tissues were configured to reflect the stages observed during the course of infection of an HIV infected individual undergoing ART. These involve the stages of primary infection, where an exponential growth or eclipse phase of the virus is observed (0 to 4 weeks from infection), followed by the stage of acute HIV syndrome or primary infection phase (5 to 9 weeks from infection), then occurs clinical latency or chronic infection (9 weeks to 8 years from infection), followed by opportunistic diseases (9 to 11 years from infection), where a rise in the viral load is again observed and death[97].

These phases were calculated as time frames based on the infected time of the individual. For the initial test where we simulated four months of infection and for the second test two years about four months of infection was simulated. The parameters for the phases of infection for the tissues for each test are described in the main text **Table 3**.

As shown in main text **Table 2** the duodenum tissue lacks any sampled sequences from the initial sampling effect. Therefore, its phasing of generations has been configured to account for this as described in main text **Table 3**. Due to the integration of transmission of viral particles between tissues, we are able to introduce viral particles into the tissues such as the duodenum.

##### 4.6.5. Transmission of viral particles between tissues

For the second test due to the incorporation of multiple tissue structures, we activated Apollo's tissue transmission feature. Using the works of Chaillon *et al.* and Goyal *et al.* the migration patterns among the respective tissues were parametrised[72,98]. To configure the rate of transmission among the different tissues the binomial parameters of $n = 30$ and $p = 0.75$ were used. The start generation from which migration begins was set to 20. This was so that a sufficient viral population build up before migration from one tissue to the next occurs. For the transmission of viral particles between tissues Duodenum to Colon migration was set to start after generation 40. This was because as shown in main text **Table 3** the Duodenum does not have a starting viral population. Therefore, we provide time for its viral load to build up before transmission begins.

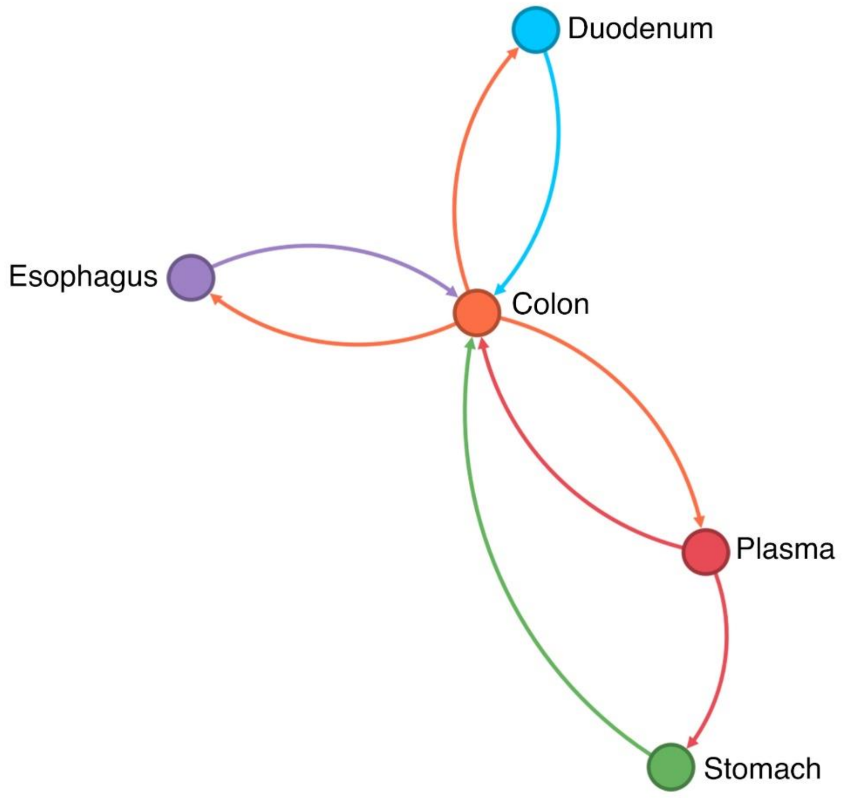

**Figure S - 24.** Within host cross tissue transmission patterns configured for the simulation. The network graph illustrates how the viral particle transmission patterns connect the five tissue structures together.

#### 4.6.6. General simulation parameters

Simulations were configured to run for a given period of time to represent the infection periods being targeted. For the first simulation, it was run from 1993-May-11 to 1993-September-14, which is about 52 generations, and for the second test we ran from 1993-May-11 to 1995-October-10 which translates to about 400 generations.

The replication time in Apollo captures time from viral attachment to the cell to the release of progeny viral particles. The average replication time was set to be 2.2 days and a standard deviation of 0.22 (gamma distribution shape 100 and scale 0.022)[97,99,100]. The progeny generation rate was set using a Binomial distribution with $r = 35$ and $p$=0.80[18,97,101–103]. The reference survival rate for the progeny was set at 0.15. This was subject to change based on the sequence occupied by a viral particle and the configured survival profile. We estimated that most viral particles produced don't survive for long and only a few survive till reproduction. This is due to the challenges faced against the host immune system, the recorded number of progeny produced on average by a viral particle and the amount of virus particles present in a host at any given time[97,104]. The affinity of each tissue's cells to the virus was estimated using a gamma distribution of $\text{shape} = 8$ and $\text{scale} = 6$. We factored that these parameters provided the most accurate HIV growth rate within the body.

#### 4.6.7. Results of the first test simulating four months of infection

The goal of this test was to start with the initial 12 sequences obtained from the PBMC tissue in the first visit and then extract the 11 sequences from the second visit at the end of the simulation. We realized that the 11 sequences from the second visit can be summarised into four unique sequences. We ran three replicate tests. In all instances, we were able to extract all four target sequences, in their entirety.

Analyzing the rate of change of the viral population we saw that Apollo captured the four phases observed during the course of viral infection (**Figure S - 25**).

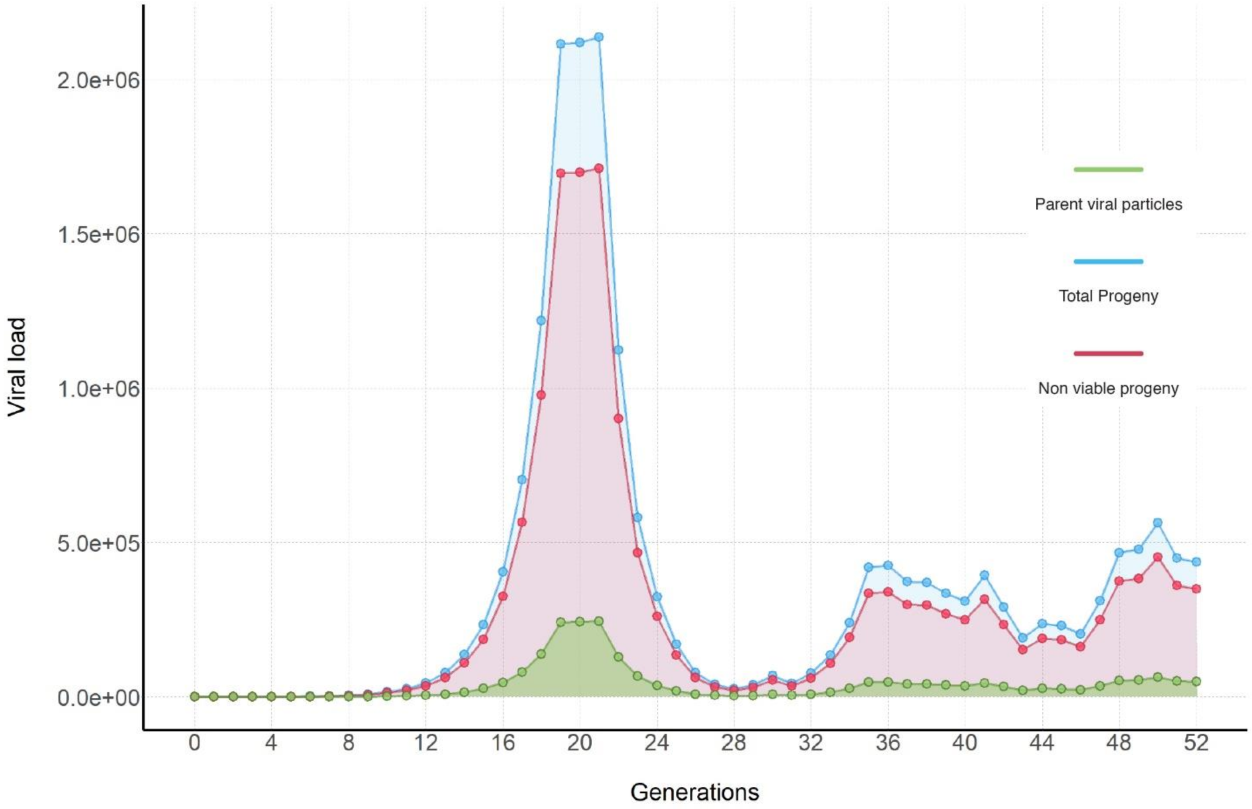

**Figure S - 25.** Change in viral population infecting the host as simulated by Apollo. We can see that the initial eclipse phase, followed by the acute infection phase and subsequently the period of clinical latency, and finally the cause of opportunistic infection leading to an eminent rise in the viral population once more. The viral particles that survived from the progeny to maturity to undergo cell attachment and reproduction are depicted in green. The progeny generated in each generation are shown in blue and out of them those that did not survive till reproduction are depicted in red.

We can even observe the latent rise in the viral population due to the incidence of opportunistic diseases and HIV viral particles that are resistant to therapeutics and host immune responses. This is caused by the fact that we included a fitness landscape model for viral survivability based on variation. This survivability model was designed by analyzing the sequences that were obtained during the second sampling efforts in the real world.

Next using Apollo’s utility tool Haplotype counter, we were able to analyze the different viral strains that were generated during the simulation and see if we had recovered the target four sequences (**Figure S - 26**).

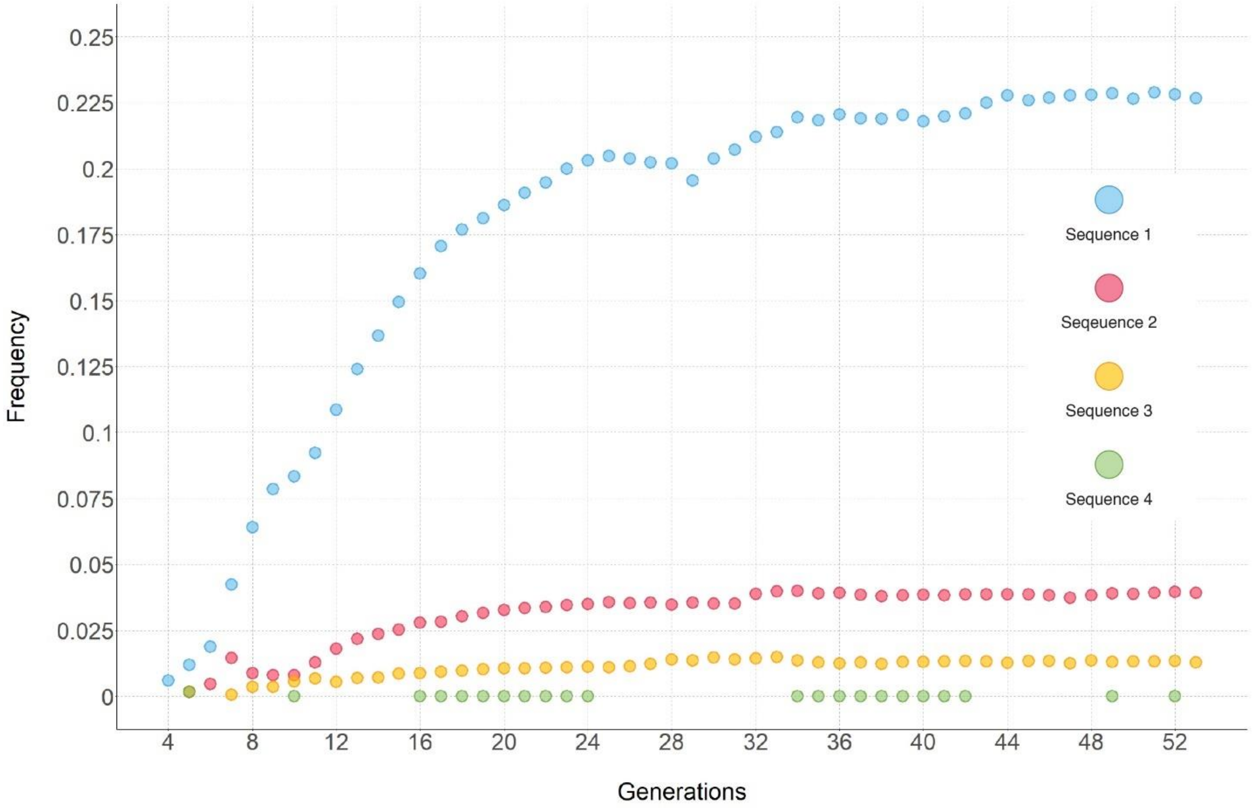

**Figure S - 26.** Details on the recovered four sequences via simulation. The graph depicts the frequency with which each sequence occupied the within host viral population at each generation that they were observed in. The sequence’s frequency of occurrence also reflects the survivability fitness model that was applied to Apollo, which was designed using the real-world data.

All four sequences were obtained in their entirety. We can also analyze the pedigree of these sequences and how they evolved from the first, zero generation sequences using the data generated by Apollo. This provides novel insights and adds new layers of information to the already existing real-world data. It also shows that Apollo is able to replicate real world scenarios accurately provided proper parameterization.

Apollo took less than three days on average to complete the above simulations. Apollo conducts the simulation using less than 20GB of RAM memory and on the GPU, it used less than 6GB of VRAM.

#### 4.6.8. Results of the second test simulating two years and four months

In our second test, we attempted to test all within host mechanisms of Apollo by simulating the infections across all five tissues. We attempted to recover the sequences provided by the real-world clinical tests at the end of the simulation.

We first analyzed the change in viral populations across the different tissues and the results are depicted in **Figure S - 27**.

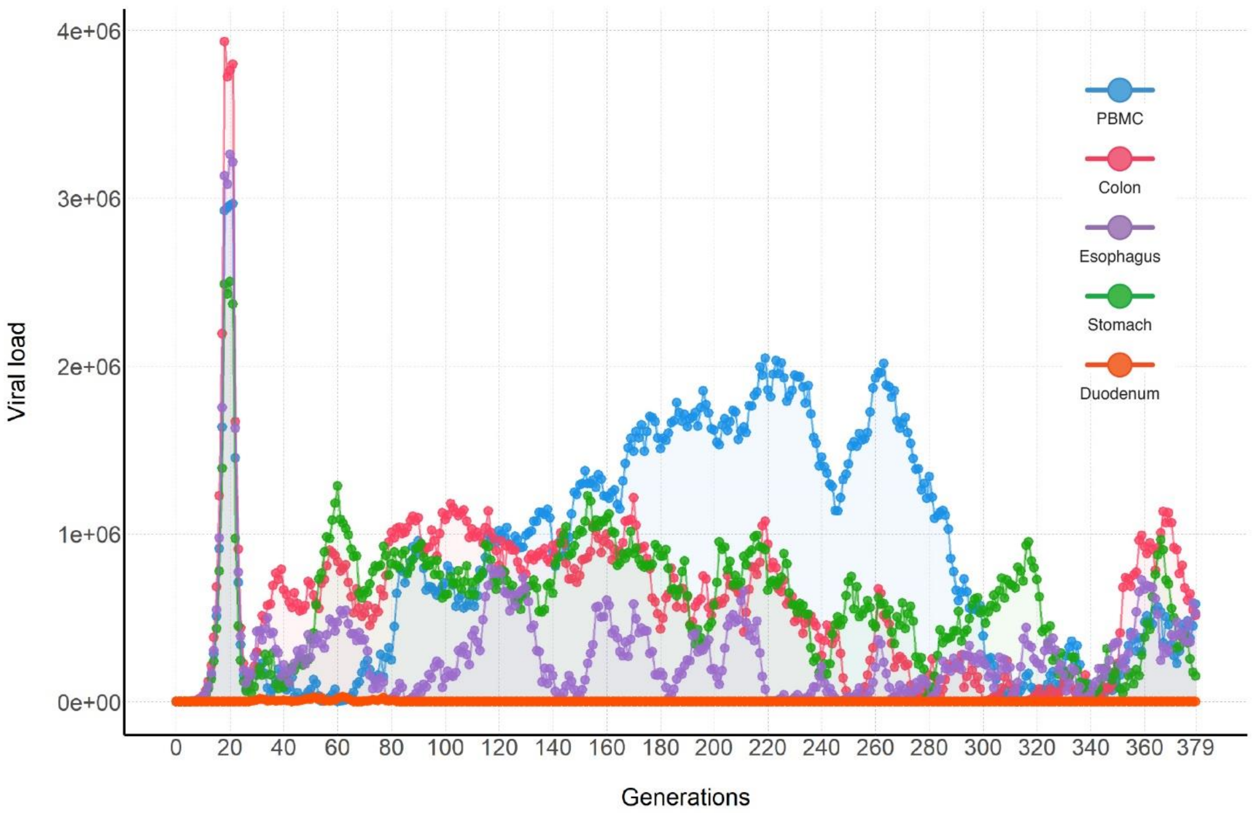

**Figure S - 27.** Change in the viral populations of each tissue with time as simulated by Apollo. The graph depicts the progeny viral population in each generation for each tissue. We can observe that the Colon and PBMC housed a majority of the viral population and the transmission between tissues helped maintain the viral populations of the other tissues as well.

Through the analysis of the change in viral population across tissues with time, we can observe how the transmission of the viral particles across tissues has helped maintain the viral populations. It has also led to more diverse viral populations caused by cross tissue infection.

Next, we analyzed our results on sequence recovery. We observed that we were able to recover most of the target sequences with an accuracy of 98.959% or greater or simply put two base mismatches or less (**Figure S - 28A**). At this accuracy, we were able to retrieve 50 different sequences. At 100% accuracy, Apollo was able to simulate 19 sequences (**Figure S - 28B**). In addition, using our complimentary tools we are able to retrieve the tissues these sequences occur in and the generations they can appear in. Additionally, Apollo enables the user to trace the genealogy of these sequences.

Therefore, through our real-world analysis, we successfully prove Apollo's capability to capture real world dynamics and render complete simulations. Our simulations were able to reconstruct the cross-tissue transmission patterns of the virus that occur within the host organism including the replication cycles, external forces of ART, and host immune responses. Finally, Apollo proved its fidelity via the accurate reconstruction of the clinical sequences complete with supplement information on the sequences' frequency of occurrence and genealogy.

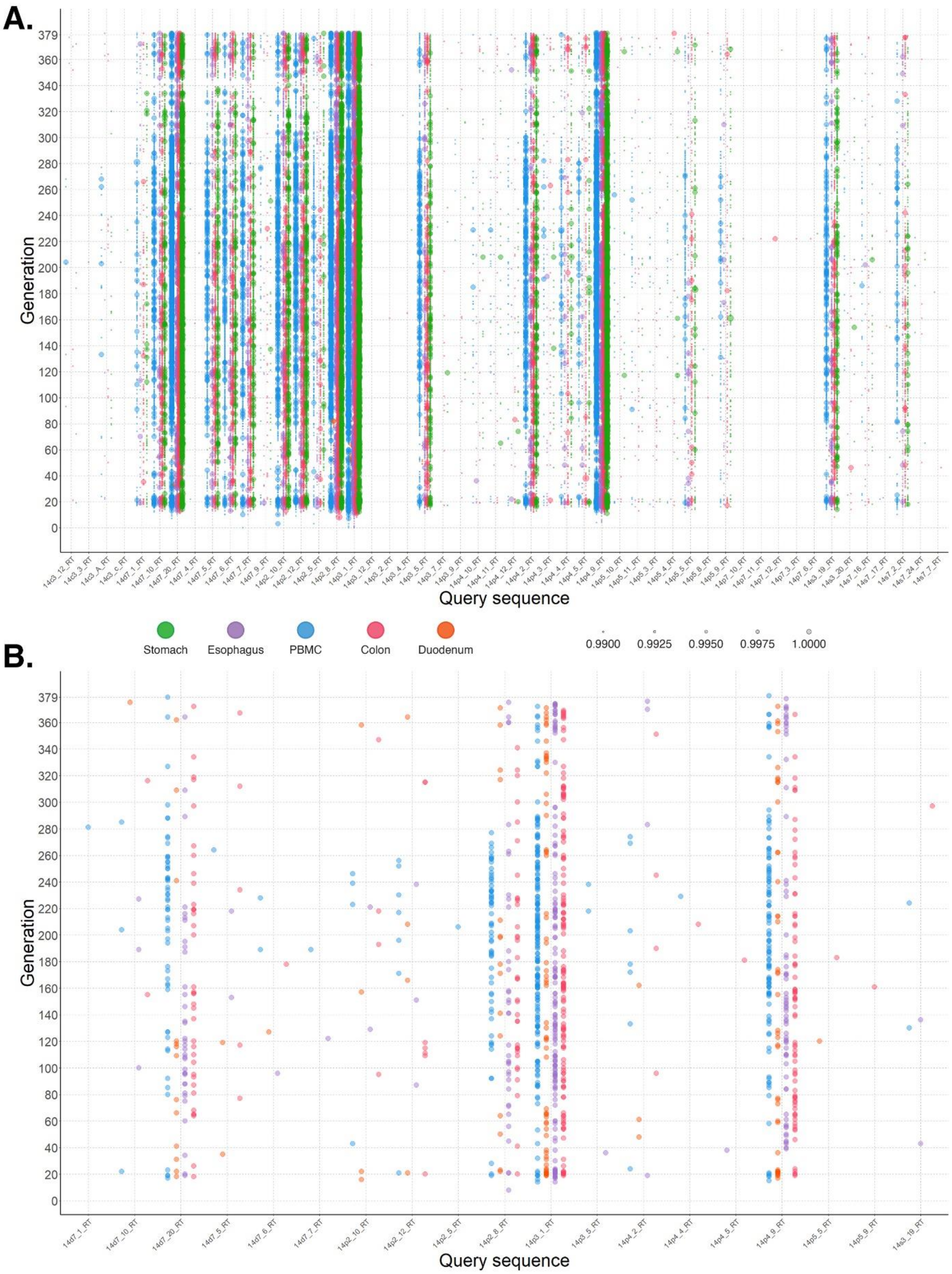

A.
Generation
Query sequence
Stomach
Esophagus
PBMC
Colon
Duodenum
0.9900
0.9925
0.9950
0.9975
1.0000
B.
Generation
Query sequence

**Figure S - 28.** Details on the reconstructed sequences by Apollo that match the sequences retrieved from the HIV infected subject via real world clinical data. The query sequence is shown in the x axis followed by the generation the sequence appeared in the y axis. The dots are coloured by the tissue where the sequence occurred. (A) Shows the sequences that were retrieved with two base mismatches or less in comparison to the target sequences. The size of the dot represents the accuracy of the sequence to the query sequence. A total of 50 sequences were retrieved via simulation. (B) Contains the subset of reconstructed sequences that were perfectly matched the sequences retrieved via real world clinical testing. A total of 19 such sequences were retrieved.

#### 4.7. Evaluation of viral inference pipelines

Through our own experiences developing transmission inference pipelines for epidemics and pandemics such as HIV's AIDS[20] and SARS-CoV-2's COVID19[105] we realized that it was challenging to quantify the accuracy of our predictions without gold standard datasets. The generation of gold standard datasets was difficult due to the lack of simulators capable of accounting for within host evolutionary dynamics with the integration of the hierarchies of the network, host, cellular, and viral genomics.

##### 4.7.1. Experimental goal and setup

We aimed to prove the viability of using Apollo to generate gold standard data. These data sets can then be utilized to assess the accuracy of predictions made by inference tools and pipelines. We analyzed the inferences made using our pipelines that utilized the TransPhylo[18,106] and BEAST2 software.

We began by simulating the infection of an epidemic in a population of 300 individuals connected via an Erdős-Rényi model with a 0.75 probability of linkage (**Figure S - 29**). We had three individual types: non LTFU (0.70), complete LTFU (Lost To Follow Up) (0.15), and partial LTFU (0.15). The varying parameters of the profile types are shown in **ONLINE METHODS Table 4**.

The non LTFU profile type represents individuals who after being sampled will conduct quarantine and treatment activities. Thereby they will be removed from the infectious population. The Lost to Follow Up (LTFU) individuals are those that remain infectious even after sampling. They are found in real world settings such as in HIV where LTFU takes up about 30% of the infected population. We have segregated LTFU individuals into two categories. Those that remain infectious completely unphased and those whose infectivity is reduced. Respectively we have labelled them as Complete LTFU and Partial LTFU. To also factor in the death factor of the population we have reduced the terminal load of the partial LTFU population so that those whose viral load exceeds a particular threshold will cause the host to reach mortality. The simulation was run with a start date of May 11th of 1993.

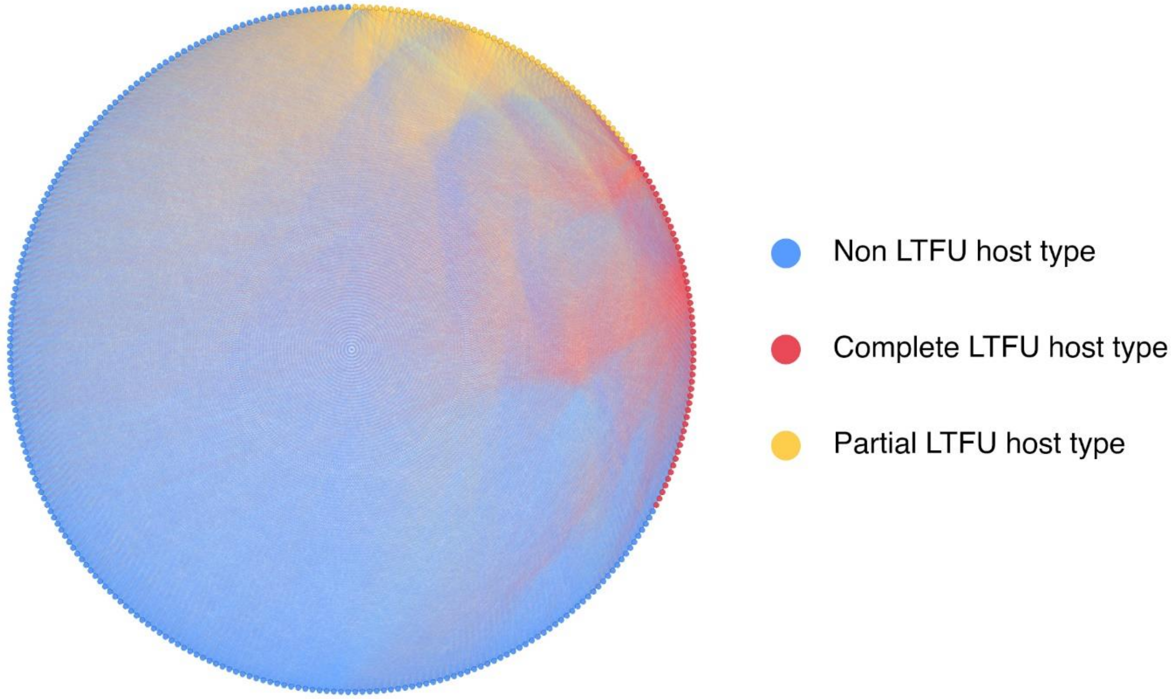

**Figure S - 29.** Contact network used for the simulation of the epidemic. Individuals are connected via an Erdős-Rényi graph model. The host types of non LTFU, complete, and partial LTFU are distributed across the network at percentages of 70%, 15%, and 15%.

To test TransPhylo we need to activate Apollo's sampling mechanic. TransPhylo is dependent on phylogenetic trees with tip dates. The tip dates are the sampling dates. The sampling mechanic was used to conduct 25 sampling events at random and the rate was configured using a Binomial distribution of $n = 10$ and $p = 0.25$.

These sequences were then used to construct a Bayesian phylogenetic tree in BEAST2[107] and the resultant tree was used to infer the transmission network using TransPhylo. We then evaluated the results from TransPhylo with the gold standard data from Apollo. Hence, we are now capable of quantitatively validating the predictions from the transmission inference pipeline.

For the generation of the phylogenetic tree, BEAST2 was used. Tip dates were activated along with a gamma site model and a GTR substitution model. The clock model was an optimized relaxed clock, and the prior was the birth death skyline serial model. The MCMC chain was of length $10^9$. The final tracer diagram was evaluated and ensured that each parameter had an

Estimated Sample Size (ESS) greater than 200 (**Figure S - 30A**). TransPhylo was configured to best represent the simulation using the parameters depicted in main text **Table 5** and the remaining parameters were left at default.

We first evaluated the TransPhylo pipeline by studying its tracer diagrams (**Figure S - 30B**). On ensuring convergence of TransPhylo's algorithm we then extracted the predicted transmission network along with predictions regarding the temporal transmission tree and incidence of disease in the population (**Figure S - 30**). We then compared these predictions to Apollo's gold standard data.

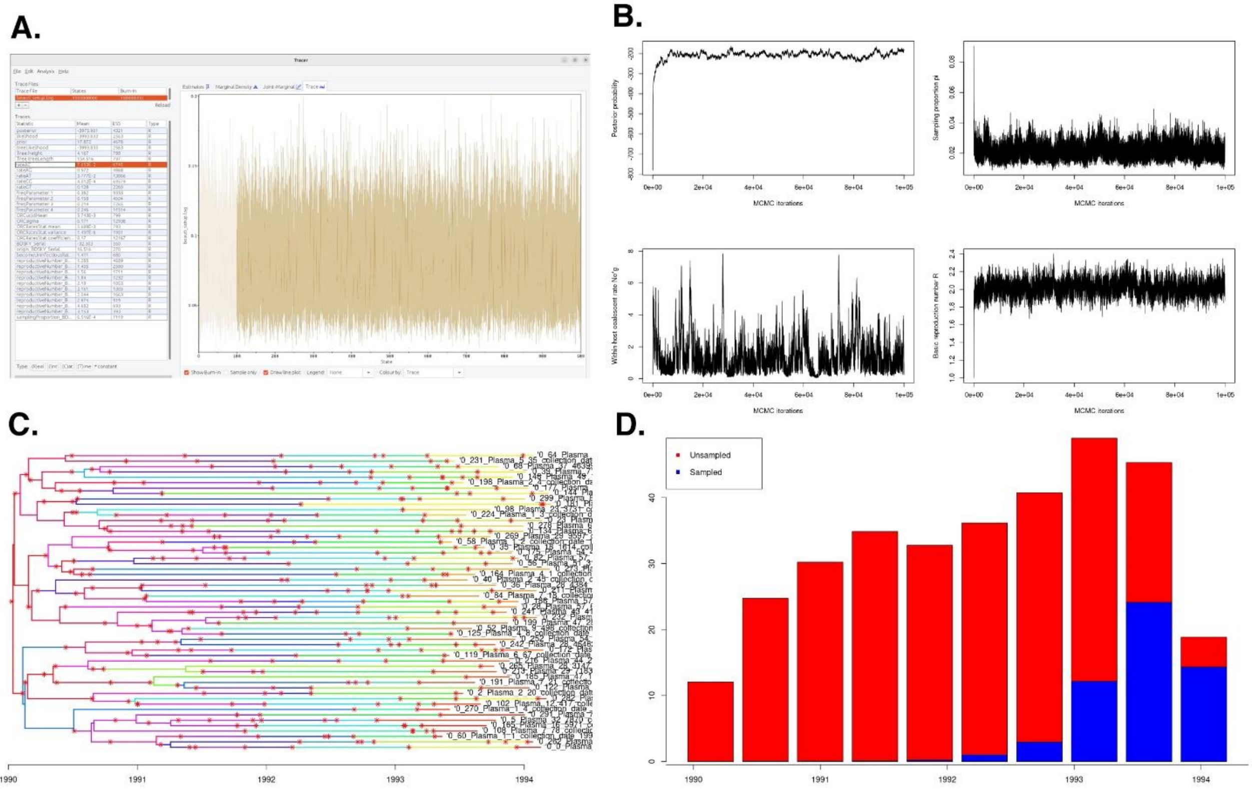

**Figure S - 30.** Analysis of the pipeline's BEAST2 and TransPhylo processes. (A) Depicts the generated tracer diagrams from BEAST showing the convergence of its inferences with ESS greater than 200. (B) TransPhylo generated tracer diagrams show that the algorithm has reached stabile convergence. (C) TransPhylo inferred transmission tree and (C) plot of incidences of sampled to unsampled cases over time. Observing (B) and (C) itself we can see that there is an error in the inferences as the start of infection has been predicted to be 1990 instead of 1993. Even the first occurrence of sampled individuals is placed into 1992.

#### 4.7.2. Evaluation of inference pipeline predictions against the gold standard

Initial observations showed major deviations in the inferred transmission network from that of Apollo's ground truth. Using this information, we attempted to fine tune our pipeline's parameters. However, we saw a tendency by the pipeline to overestimate the TMRCA and miss the disease incidence time over two years. The inference pipeline also showed an overestimation of the infected population and missed direct host to host transmission.
