## Supplementary material for "Apollo: A comprehensive GPU-powered within-host simulator for viral evolution and infection dynamics across population, tissue, and cell": User Manual

### Apollo: A comprehensive GPU powered Within-host Viral simulator complete with tissue and cellular hierarchies for studying viral evolutionary and infection dynamics.

Apollo powered by CATE

Deshan Perera, Alexander Platt and Quan Long

Department of Biochemistry & Molecular Biology, Cumming School of Medicine, University of Calgary, Calgary, AB T2N 4N1, Canada.

3.3.7.16. Infection sequence Fixed (*profile_h file parameter*) 32

3.3.7.17. Sampling effect (*profile_h file parameter*) 32

3.3.7.18. Sampling less infectious effect Alpha (*profile_h file parameter*) 32

3.3.7.19. Sampling less infectious effect Beta (*profile_h file parameter*) 33

3.3.7.20. Tissue profiles (*profile_h file parameter*) 33

3.3.7.21. Tissue ($\boldsymbol{t}$) (*profile_h file parameter*) 33

3.3.7.22. Cell limit (*profile_h file parameter*) 33

3.3.7.23. Cell limit Binomial trials (*profile_h file parameter*) 34

3.3.7.24. Cell limit Binomial probability (*profile_h file parameter*) 34

3.3.7.25. Viral distribution type (*profile_h file parameter*) 34

3.3.7.26. Viral distribution Binomial trials (*profile_h file parameter*) 34

3.3.7.27. Viral distribution Binomial probability (*profile_h file parameter*) 34

3.3.7.28. Viral distribution Gamma shape (*profile_h file parameter*) 35

3.3.7.29. Viral distribution Gamma scale (*profile_h file parameter*) 35

3.3.7.30. Replication phases (*profile_h file parameter*) 35

3.3.7.31. Number of phases (*profile_h file parameter*) 36

3.3.7.32. Phase ($\boldsymbol{x}$) Mode (*profile_h file parameter*) 36

3.3.7.33. Phase ($\boldsymbol{x}$) Time ratio (*profile_h file parameter*) 36

3.3.7.34. Phase ($\boldsymbol{x}$) Variance (*profile_h file parameter*) 37

3.3.7.35. Phase ($\boldsymbol{x}$) Alpha (*profile_h file parameter*) 37

3.3.7.36. Phase ($\boldsymbol{x}$) Beta (*profile_h file parameter*) 37

3.3.7.37. Overview of host profile parameter file 37

3.3.8. Overview of the Node/ Host parameter file 39

3.4. Sequence parameter file 40

#### SECTION 1. SOFTWARE OVERVIEW

Apollo is a forward in time viral simulation software. It is written in C/C++ and CUDA programming languages and designed to run on Linux and Unix kernels. Apollo requires a CUDA enabled NVIDIA GPU.

Apollo is freely available for download and use under the MIT license from the Open Software Initiative. It is provided as part of the CATE software and is available at the following links:

**GitHub:** https://github.com/theLongLab/CATE

**GitHub Wiki:** https://github.com/theLongLab/CATE/wiki/Apollo

**Anaconda:** https://anaconda.org/deshan_CATE/cate

##### 1.1. User interaction

Apollo is a command line driven software. Users interact with the software by passing instructions through the terminal. Typically, the desired function along with our bespoke parameter files are passed into the software. During execution of the software, it provides the user with detailed messages of its progress, in addition to any errors and possible solutions to rectify them.

Apollo comes equipped with its main simulator function as well as five complementary tools.

##### 1.2. Installation

Apollo can be installed via CATE[1,2] to be used either online or on-device.

###### 1.2.1. Online installation

Apollo can be used via Google Colab. The link to the Google Colab notebook can be accessed here:

Open in Colab

###### 1.2.2. On-device installation

For on device installation, it can be installed either via Anaconda or building of the GitHub code.

###### 1.2.2.1. Install via Anaconda

To install CATE using the Anaconda package execute the following command.

`conda install deshan_cate::cate`

###### 1.2.2.2. Build using GitHub

To build the GitHub source code follow the steps below:

1. Download the repository from GitHub:

   `git clone https://github.com/theLongLab/CATE`

2. Navigate into the download folder:

   `cd CATE/`

3. Load the CUDA compiler module if necessary:

   `module load cuda/11.3.0`

4. Compile the code:

   `nvcc -std=c++17 *.cu *.cpp -o "CATE"`

Test the programme installation by running the command: `CATE - h`

#### SECTION 2. AVAILABLE FUNCTIONS

Apollo comes with its main simulator function and five other complementary tools designed to analyse the generated results. Detailed descriptions of all these functions are described in **Table S - 1**.

**Table S - 1.** A brief description of Apollo's functions. It shows the relevant arguments that have to be passed into Apollo via the command line for the specific function's execution.

| Function name | Argument | | Description |
|---|---|---|---|
| Simulator | --simulator | -sim | This is Apollo's main function. Conducts the epidemic simulation based on the user configured parameters. |
| ***Tools*** | | | |
| Haplotype retriever | --hapretrieve | -hr | Retrieves unique sequence configurations complete with the tissue, generation, sequence configuration and frequency. The haplotype retriever also provides information on virions sequences that became parents, progeny that survived till the end of the generation and all sequences. |
| Pedigree retriever | --pedretrieve | -pedr | The pedigree retriever maps the parental relationships of a given sequence within a given host's tissue from a given generation. It provides the pedigree of all sequences found in the given generation of the host's tissues. The pedigree is traced for each sequence till the initial ancestral genomes are identified. |
| Segregating sites matcher | --segmatch | -segm | In an individual, identifies a given set of query sequences by alignment to simulated sequences by the given degree of matching segregating sites. For each matched sequence a summary of matched sequence completed with the percentage matched, mismatched bases if any and their occurrences in generation and tissue are provided. |
| Base substitution model to JSON | --site2json | -s2j | Converts the base substitution csv to Apollo's JSON format. |
| Recombination hotspots to JSON | --recom2json | -r2j | Converts the recombination hotspots csv to Apollo's JSON format. |

#### SECTION 3. SIMULATOR PARAMETERIZATION

Apollo's parametrization is done using four main parametrization files. They are as follows:

A. Master parameter file
B. Network parameter file
C. Node/ Host parameter file
D. Sequence parameter file

These four main parameter files can be used to configure the simulation in its entirety and are used to point to other subordinate parameter files. In this section we will look at each parameter file and its respective parameters. We will then explain each parameter and its use. If a parameter files' configuration requires additional files, we will clarify their structure and implementation as well.

It should be noted that the parameters themselves are case sensitive. Essentially the sentence case of their initialisations as listed in this section should be followed as is. Through each subheading under the parameter file section, we will explain the respective parameter. The heading will specify its sentence case for initialisation.

When assigning values for variables it should be noted that all lists, locations, values with decimal numbers and text should be written enclosed with quotation marks (""). Numerical integer values have to be written without quotation marks.

Commenting on Apollo's JSON script files can be done using a hash (#). Any text preceded by a hash are considered as comments and will be ignored by Apollo.

##### 3.1. Master parameter file

The master parameter file is the simplest of the four main parameter files and it is used to configure the hardware, folder, and overall simulation parameters. It is also the parameter file responsible for pointing Apollo to the other three main parameter files. Now let's look at its individual parameters.

###### 3.1.1. Hardware parameters

The following parameters are used to configure the hardware that is available for Apollo's use.

###### 3.1.1.1. CUDA Device IDs

Used to configure respective CUDA devices to be used by Apollo and the number of CUDA devices. The use will specify the IDs of the CUDA devices to be used separating each device ID by a comma. If only one CUDA device is available, the user can just list the ID of the single CUDA device to be used.

E.g.: *"Cuda device IDs":"0,1,3"*

In the above example Apollo will recognise that it has three separate CUDA devices at its disposal. It will then configure itself to the use the CUDA devices that the respective IDs point to.

###### 3.1.1.2. CPU cores

Used to configure the number of CPU cores available for use.

E.g.: *"CPU cores":5*

###### 3.1.1.3. GPU max units

Specifies the maximum number of parallelisation tasks the GPU is allowed to execute at a time. This parameter is used to prevent GPU overloading. Apollo will conduct automatic scheduling of its parallelisation functions based on this value. Users should take into account the amount of GPU RAM available as well as their GPUs processing capability when configuring this parameter.

E.g.: *"GPU max units":100000*

###### 3.1.1.4. Process cell rate

Specifies the number of cells that will be processed at a time. This parameter is used to prevent overloading of computer RAM. Similarly, with the GPU users should take into consideration their computers availability of RAM memory when configuring this parameter.

E.g.: *"Process cell rate":5*

###### 3.1.1.5. Multi read

Used in the presence of SSD hardware. Unlike Hard Disk Drives (HDDs) SSDs are capable of conducting parallel read and write functions. This parameter takes a Boolean value of either “Yes” or “No”. If set to Yes, Apollo will conduct parallel read and write executions whereas if set to No it will conduct single read and write executions.

E.g.: *"Multi read":"YES"*

##### 3.1.2. Folder configuration and management

The following parameters are used to point Apollo to the location of the output and intermediary folders it needs of its execution as well as configure the number of sequences that will be stored in each batch sequence folder in the intermediary folder. The parameters that are responsible for folder management to control the number of files present at a time will also be explored here. We will explain this folder architecture more in the software architecture section. It does not matter if the folder does not exist in the specified location. In such an event Apollo will create the folder for the user automatically.

###### 3.1.2.1. Output folders

This is the designation where all the results will be written to. Apollo creates two folders within the output folder. They are the network_Data and the node_Data folders. The network_Data folder contains summary information on the simulation that has occurred in relation to the whole population. The node_Data folder contains the information of each individual host that has gotten infected.

E.g.: *"Output folders":"results"*

###### 3.1.2.2. Intermediate folders

The intermediate folder contains information that is needed by Apollo during a simulation run. These are usually the individual viral particle information from each generation.

E.g.: *"Intermediate folders":"intermediates"*

###### 3.1.2.3. Intermediate Sequences per file

This parameter is used to control the maximum number of sequences that can be stored in an intermediate file. This parameter is taken into consideration because most clusters have a file limit per user. If each viral particle is stored as an individual sequence this could cause the user to run out of their disk usage quota. To provide the user with a level of control to prevent this from occurring we have provided this parameter.

E.g.: *"Intermediate Sequences per file":1000*

###### 3.1.2.4. Enable folder management

Folder management builds upon the previous parameter. If this parameter is enabled Apollo will reduce the folders from a completed generation into a tar file. This is a Boolean variable that can take the values of either “YES” or “NO”.

E.g.: *"Enable folder management":"Yes"*

###### 3.1.2.5. Compress folders

If the above folder management parameter (3.1.2.4) is set to enabled (“YES”), then instead of reducing the folder to a tar file they will undergo an additional compression step and be compressed to tar.gz. This is ideal where in situations where the user is limited by their on-disk storage.

E.g.: *"Compress folders":"NO"*

###### 3.1.3. Overall simulation management

These parameters are responsible for setting up the start date and if needed to specify length of time the simulation needs to run for.

###### 3.1.3.1. Start date

Use to specify the start date from which the simulation starts, so that the progression of the simulation can be traced using chronological time as well.

E.g.: *"Start date":"2024-01-10"*

###### 3.1.3.2. First infection

This controls how the first patient zero host is infected. It can be either "RANDOM" or "FIXED". In Random the provided parent sequences are infected at random into a host's entry tissue structures. However, in Fixed, the infection of patient zero's tissue is dependent on the configuration of the tissue sub folders in the parent sequence folder.

E.g.: *"First infection":"Random"*

###### 3.1.3.3. Stop after generations

If the user wants to stop the simulation after a given period of time, they can activate this parameter. It takes a Boolean value of either "YES" or "NO".

E.g.: *"Stop after generations":"Yes"*

###### 3.1.3.4. Mode to stop

This parameter works in conjunction with the above parameter (3.1.3.3). If it is set to "YES" this will specify the mode of stopping. This is a Boolean variable of either "Generations" or "Date". If set to "Generations" it will stop after a fixed number of generations and if set to "Date" it will stop after a specified date.

E.g.: *"Mode to stop":"Generations"*

###### 3.1.3.5. Number of generations

If the above parameter for method of stopping the simulations (3.1.3.4) is set to "Generations" then this parameter will be activated. Apollo will terminate after the user specified number of generations have been successfully run in the overall simulation.

E.g.: *"Number of generations":25*

###### 3.1.3.6. End date

This parameter is activated if the method of stopping the simulations (3.1.3.4) is set to "Date". Apollo will terminate after the user specified date has been reached in the overall simulation.

E.g.: *"End date":"2024-01-12"*

##### 3.1.4. Locations for Network, Nodes master and Sequence master parameter files

These three parameters are used to specify the locations of the other three main parameter files.

###### 3.1.4.1. Network profile

Specifies the location of the network parameter file used to design the contact network.

E.g.: *"Network profile":"/network_Profiles/network_Master.json"*

###### 3.1.4.2. Nodes master profile

Specifies the location of the overall node profile file. This parameter file will configure the overall parameters common to each host.

E.g.: *"Nodes master profile":"/individual_Profiles/node_Master.json"*

###### 3.1.4.3. Sequence master profile

Specifies the location of the sequence configuration parameter file.

E.g.: *"Sequence master profile":"/sequence_Profiles/sequence_Master.json"*

##### 3.1.5. Overview of Master parameter file

These are all the parameters involved in the Master parameter file. We have provided an example of a fully configured Masters parameter file in Figure S - 1.

Master_parameter_file.json

```
{
  # Configuration of the Cuda device IDs accessible to Apollo
  "CUDA Device IDs":"0",

  # Hardware configuration parameters
  "CPU cores":40,
  "GPU max units":100000,
  "Process cell rate":25,
  "Multi read":"YES",

  # Output and Intermediate folder locations.
  "Output folders":"results",
  "Intermediate folders":"intermediates",

  # The maximum number of sequences that can be stored in an intermediate file
  "Intermediate Sequences per file":10000,

  # Indicates the start date for the simulation
  "Start date":"2024-01-10",

  # Indicate whether the user wants to run the simulation for a specified period of time
  "Stop after generations":"Yes",
  # Generations or Date can be used to stop the simulation as well
  "Mode to stop":"Generations",
  "Number of generations":50,
  "End date":"2024-01-12",

  # Network profile parameter file location
  "Network profile":"/network_Profiles/network_test.json",

  # Node profile parameter file location
  "Nodes master profile":"/individual_Profiles/node_Master.json",

  # Sequence profiles parameter file location
  "Sequence master profile":"/sequence_Profiles/sequence_Master.json",

  # Folder management controls
  # Compresses folders to a TAR.GZ or reduces them to a single TAR file
  "Enable folder management":"Yes",
  "Compress folders":"No"
}
```

**Figure S - 1:** Fully configured parameter file for a simulation run. Parameters do not need to be listed in any particular order, but they are case sensitive.

##### 3.2. Network parameter file

Th network parameter file is used to select the graph model for the contact network. The user has five network graph models to select from. They are random model, Erdős-Rényi model, Barabási Albert model, standard caveman model and dynamic caveman model. We will look at each individual model's parameter.

###### 3.2.1. Overall network model parameters

Parameters that affect the generation of the contact network.

###### 3.2.1.1. Network type

It can take either one of the five values. Each represents a type of graph model.

**A. Random model:** Generates a random graph where there is a fixed number of nodes, and each node has the same probability of being assigned to each other. It is a modification on the Erdős-Rényi model and is the simplest form of graph provided by Apollo.
**B. ER model:** This the standard implementation of the Erdős-Rényi model of $\boldsymbol{G}(\boldsymbol{N}, \boldsymbol{p})$. Where $\boldsymbol{N}$ nodes form a network graph model with a probability of $\boldsymbol{p}$ for two nodes being linked.
**C. BA model:** Generates a contact network based with a fixed number of nodes based on the Barabási-Alabert (BA) model. Apollo implements a modification to the standard model proposed by Albert Barabási and Réka Albert by allowing the user to control the number of nodes an incoming node will connect to.
**D. SC model:** Generates a contact network based on the Caveman model. The user can specify the number of cohorts/ caves ($\boldsymbol{C}$) to be present and the size of each cave ($\boldsymbol{s}$) (number of nodes per cave). The total number of nodes in the network will be the product of the two values ($\boldsymbol{Cs}$).
**E. DC model:** This a modification on the above caveman model. It allows the user to generate a network with caves of variable sizes. Additionally, more than one node can form contacts with its neighbours as well as nodes can form connections beyond just their neighbouring caves. The total node number present in the network can change based on the user's specifications.

E.g.: *"Network type":"BA model"*

###### 3.2.2. Random model

Parameterisation of the random model.

###### 3.2.2.1. Random model number of nodes

Specify the number of nodes that will be present in the contact network for the random graph model.

E.g.: *"Random model number of nodes":100*

###### 3.2.3. Erdős-Rényi model

Parameterisation of the Erdős-Rényi model.

###### 3.2.3.1. ER Random model total number of nodes

Specifies the total number of nodes that will be present in the ER model network.

E.g.: *"ER Random model total number of nodes”:100*

###### 3.2.3.2. ER Random model probability of linkage

Probability that two nodes in the generated contact network will be linked.

E.g.: *"ER Random model probability of linkage”:“0.75”*

###### 3.2.4. Barabási Albert (BA) model

Parametrisation of the BA model.

###### 3.2.4.1. BA model number of nodes

Specify the number of nodes that will be present in the contact network for the BA model.

E.g.: *"BA model number of nodes":100*

###### 3.2.4.2. BA model standard new connections

This is a modification for the standard BA model. Typically, each incoming node will form a connection only with a single existing node weighted by the power law. Apollo enables the user to customise the BA model where the incoming node can attach to multiple existing nodes (user specified number).

This parameter can take either one of three values: “Fixed”, “Negative binomial” and “Poisson”. The number of connections the incoming node will attach to is based on these distributions.

If “Fixed” is selected the number of new connections will always be a fixed number, if “Negative binomial” is selected the number of connections will follow a negative binomial distribution and finally “Poisson” will ensure the number of new connections formed by an incoming node will follow a Poisson distribution.

E.g.: *"BA model standard new connections":"FIXED"*

###### 3.2.4.3. BA model fixed new connections

This parameter is activated if the above parameter (3.2.4.2) is set to “FIXED”. This ensures that the number of new connections formed by an incoming node is always the same. If this is set to 1 then the generated contact network is the standard BA model as defined by Barabási and Albert.

E.g.: *"BA model fixed new connections":1*

###### 3.2.4.4. BA model Negative binomial successes

Parameter used to set the number of successes ($r$) in the binomial distribution if 3.2.4.2 is set to “Negative binomial”.

E.g.: *"BA model Negative binomial sucesses":1*

###### 3.2.4.5. BA model Negative binomial probability

Parameter used to set the probability of success ($p$) in the binomial distribution if 3.2.4.2 is set to “Negative binomial”.

E.g.: *"BA model Negative binomial probability":"0.75"*

###### 3.2.4.6. BA model Poisson mean

Parameter used to set the mean ($\lambda$) of the Poisson distribution if 3.2.4.2 is set to “Poisson”.

E.g.: *"BA model Poisson mean":"1"*

**3.2.5. Standard Caveman model**

We refer to this option as the standard caveman model since Apollo provides a modified version of the caveman model which we refer to as the dynamic caveman model.

**3.2.5.1. SC_model number of caves**

Specifies the number of caves that will be present in the contact network.

E.g.: *"SC_model number of caves":10*

**3.2.5.2. SC_model number of nodes per caves**

Specifies the number of nodes per individual cave in the contact network.

E.g.: *"SC_model number of nodes per caves":10*

**3.2.6. Dynamic caveman model**

This is Apollo's modified version of the standard caveman model. Apollo essentially provides the user with control of the number of individuals that can be present in a cave. The number of individuals in each cave does not have to be fixed. Additionally, the user can control the number of nodes that form connections with the neighbouring caves as well as those that will form connections with caves outside their immediate vicinity.

It should also be noted that unlike in the standard caveman model no rerouting of intra cohort nodes will occur when connections with other caves occur.

**3.2.6.1. DC_model number of caves**

Specifies the number of caves that will be present in the contact network.

E.g.: *"DC_model number of caves":25*

###### 3.2.6.2. DC_model node distribution

The parameter is used to determine the distribution that will be used to determine the number of nodes that will be present in the caves in the dynamic caveman model. It can take either one of the two values “Negative binomial” or “Poisson”.

E.g.: *"DC_model node distribution":"Negative binomial"*

###### 3.2.6.3. DC_model node Negative binomial successes

Parameter used to set the number of successes ($r$) in the binomial distribution if 3.2.6.2 is set to “Negative binomial”.

E.g.: *" DC_model node Negative binomial sucesses":1*

###### 3.2.6.4. DC_model node Negative binomial probability

Parameter used to set the probability of success ($p$) in the binomial distribution if 3.2.6.2 is set to “Negative binomial”.

E.g.: *"DC_model node Negative binomial probability":"0.75"*

###### 3.2.6.5. DC_model node Poisson mean

Parameter used to set the mean ($\lambda$) of the Poisson distribution if 3.2.6.2 is set to “Poisson”.

E.g.: *"DC_model node Poisson mean":"1"*

###### 3.2.6.6. DC_model neighbouring nodes percentage

Defines the probability of the nodes in a cave being neighbor joining nodes.

E.g.: *"DC_model neighbouring nodes percentage":"0.25"*

###### 3.2.6.7. DC_model global nodes percentage

This parameter is used to determine the probability of the neighbour joining nodes becoming global nodes that will form relationships with distant caves.

E.g.: *"DC_model global nodes percentage":"0.55"*

##### 3.2.7. Overview of Network parameter file

These are all the parameters involved in the Master parameter file. We have provided an example of a fully configured Masters parameter file in **Figure S - 2**. When configuring the file, the user only needs to declare the parameters that are associated with their selected network graph model.

Network_parameter_file.json

```
{
  # Select network type

  # Random Model is selected by typing Random model
  # Barabasi Albert Model is selected by typing BA model
  # Standard Caveman Model is selected by typing SC model
  # Dynamic Caveman Model is selected by typing DC model

  "Network type":"DC model",

  # Total number of caves that will be present in the DC network model
  "DC_model number of caves":25,

  # Determine the number of nodes that will be present in each cave in the network
  # Negative binomial, every caves number of nodes will determined by a NEGATIVE BINOMIAL
distribution.
  # Poisson, every caves number of nodes will determined by a POISSON distribution.
  "DC_model node distribution":"Negative binomial",

  # Assignment of parameter values; sucesses and probability for DC model with NEGATIVE BINOMIAL
for per cave node distributions.
  "DC_model node Negative binomial sucesses":10,
  "DC_model node Negative binomial probability":"0.55",

  # Determine the percentage of nodes that will take part in forming neighbouring node
attachments and from them how many will form global attachments.
  "DC_model neighbouring nodes percentage":"0.25",
  "DC_model global nodes percentage":"0.55"
}
```

**Figure S - 2:** Fully configured network parameter file for a simulation. The user has selected a dynamic caveman model with a negative binomial distribution for the generation of the number of nodes per cave.

##### 3.3. Node/ Host parameter file

The node profile parameter file is a bit more complicated than the previous two parameter files. This is mainly due to the fact that in its configuration we have to configure separate parameter files for the individual node profile types it uses that may be present in a population (**Error! Reference source not found.A**, the different coloured nodes, each node color represents a different host profile). We will also introduce the use of nesting in the declaration of parameters.

###### 3.3.1. Viral replication time

The replication time of a virus is defined as the time taken from viral attachment to a host cell to release of progeny. This is also known as the generation time. In Apollo it is measured in days and is defined in the form of a gamma distribution.

###### 3.3.1.1. Shape replication time

Defines the shape of the gamma distribution for replication time.

E.g.: *"Shape replication time":"20.0"*

###### 3.3.1.2. Scale replication time

Defines the scale of the gamma distribution for replication time.

E.g.: *"Scale replication time":"0.020"*

###### 3.3.2. Duration of infection

This is the typical time the virus spends within the host from initial entry of the pathogen into the host to the end of the infection. This can be dependent on a number of factors such as immune response and drug therapy, which can be included into the infection cycle (we will look at this in the later parameters), but here we define the average time the infection lasts in an average individual. The duration of infection is measured in days and is represented using a gamma distribution.

###### 3.3.2.1. Shape days in host

Defines the shape of the gamma distribution for the duration of infection.

E.g.: *"Shape days in host":"20.0"*

###### 3.3.2.2. Scale days in host

Defines the scale of the gamma distribution for the duration of infection.

E.g.: *"Scale days in host":"0.27"*

##### 3.3.3. Progeny generation

This parameter is used to capture the rate of progeny generation, or the number of progenies produced by a single viral particle during replication. Apollo provides the user with three separate distributions to chose from to define the rate of progeny production. The options are "Negative binomial distribution", "Gamma distribution" and "Poisson distribution".

###### 3.3.3.1. Progeny distribution type

Used to select either one of the three distributions to define progeny generation. The options are "Negative binomial distribution", "Gamma distribution" and "Poisson distribution."

E.g.: *"Progeny distribution type":"Negative Binomial"*

###### 3.3.3.2. Progeny Negative Binomial successes

Parameter used to set the number of successes ($r$) in the binomial distribution if 3.3.3.1 is set to "Negative binomial".

E.g.: *"Progeny Negative Binomial sucesses":10*

###### 3.3.3.3. Progeny Negative Binomial probability

Parameter used to set the probability of success ($p$) in the binomial distribution if 3.3.3.1 is set to "Negative binomial".

E.g.: *"Progeny Negative Binomial probability":"0.55"*

###### 3.3.3.4. Progeny Poisson mean

Parameter used to set the mean (λ) of the Poisson distribution if 3.3.3.1 is set to “Poisson”.

E.g.: *"Progeny Poisson mean":"10"*

###### 3.3.3.5. Progeny Gamma shape

Parameter used to set the shape ($k$) of the Gamma distribution if 3.3.3.1 is set to “Gamma”.

E.g.: *"Progeny Gamma shape":"10"*

###### 3.3.3.6. Progeny Gamma scale

Parameter used to set the scale (θ) of the Gamma distribution if 3.3.3.1 is set to “Gamma”.

E.g.: *"Progeny Gamma scale":"0.5"*

##### 3.3.4. Sampling of hosts

Sampling of infected hosts is an important aspect of an epidemiology study. In certain epidemiological model sampling can affect the disease model. For instance sampled individuals can be defined as identified sources of infection and be removed from the infectious population or become less infectious[3,4]. Apollo contains parameters to capture such dynamics as well.

###### 3.3.4.1. Sampling present

Boolean variable used to activate the sampling mechanic. It can contain the values of either “YES” which will enable sampling in the simulation or “NO” which will deactivate the sampling mechanism for the simulation.

E.g.: *"Sampling present":"Yes"*

###### 3.3.4.2. Resampling of nodes

If a sampled host can be sampled again, ensures that all infected hosts are sampled once if set to “No”. It is a Boolean variable of “Yes” or No”.

E.g.: *"Resampling of nodes":"Yes"*

###### 3.3.4.3. Limit samples obtained

Used to limit the number of times hosts are sampled. It is a Boolean variable, can take the values of "YES" or "NO". If activated by setting to "YES" the programme will complete its simulation after hosts have been sampled a fixed number of times.

E.g.: *"Limit samples obtained":"Yes"*

###### 3.3.4.4. Max samples to obtain

If the above parameter 3.3.4.3 to limit the number of times hosts are sampled is activated this parameter is used define the limit of host sampling.

E.g.: *"Max samples to obtain":20*

###### 3.3.4.5. Sampling rate Binomial trials

Rate of sampling hosts from the infected population is defined via a binomial distribution. This parameter defines the number of trials ($n$).

E.g.: *"Sampling rate Binomial trials":2*

###### 3.3.4.6. Sampling rate Binomial probability

Rate of sampling hosts from the infected population is defined via a binomial distribution. This parameter defines the probability of success ($p$).

E.g.: *"Sampling rate Binomial probability":"0.75"*

###### 3.3.4.7. Distribution per node sequences sampled

Apollo can be configured to obtain multiple sequences from each host during a sampling event. If the host being sampled using this parameter, we can control the number of times sequence samples are obtained during the sampling process.

It can be configured to take a fixed number of samples per sampling event by setting the parameter to "Fixed" or follow a binomial distribution by setting the parameter to "Binomial".

E.g.: *"Distribution per node sequences sampled":"FIXED"*

###### 3.3.4.8. Per node sampling rate Binomial trials

If the above parameter 3.3.4.7 is set to “Binomial” then this parameter is used to define the number of trials in the binomial distribution ($n$).

E.g.: *"Per node sampling rate Binomial trials":2*

###### 3.3.4.9. Per node sampling rate Binomial probability

If the above parameter 3.3.4.7 is set to “Binomial” then this parameter is used to define the probability of success in the binomial distribution ($p$).

E.g.: *"Per node sampling rate Binomial probability":"0.5"*

###### 3.3.4.10. Per node sampling rate Fixed

If the above parameter 3.3.4.7 is set to “Fixed” then this parameter is used to define the fixed number of sampling events per generation.

E.g.: *" 2.3.4.10. Per node sampling rate Fixed ":1*

##### 3.3.5. Susceptible Infected Recovered Susceptible SIRS or SEIRS models

In certain epidemiological models an individual infected once recovered will maintain lifetime immunity for the disease, hence they will be removed from the susceptible population (SIR or SEIR models). But in certain instances, the recovered individual can become susceptible to the disease again. Apollo provides the parametrisation of these models via the activation of the parameter below.

###### 3.3.5.1. Infected to Recovered

This a Boolean variable of either “YES” or “NO”. If set to “YES” an infected host once recovered will become susceptible again. If set to “NO” the host will be removed from the population once the infection has completed.

E.g.: *"Infected to Recovered":"No"*

##### 3.3.6. Tissues

Apollo provides the user with the ability to define the within host environment. The first in this endevour is to define the host tissue structures (**Error! Reference source not found.C**). The user n eeds only to take into consideration the tissues that play a role in the viral disease infection. These tissues can then be defined, and their subsequent roles can be allocated. Apollo also allows the user to configure the migration of viral particles between tissues.

Here we will look at the different parameters that have to be configured when declaring and defining tissues within the hosts. The basic tissue types are common to all hosts regardless of their host profiles. Users are provided with parameters to configure the number of tissues that can be present in a host, the tissue names, the roles played by each tissue and the migration of viral particles from one tissue to the next.

It should be noted that Apollo requires at least one tissue type to be declared at minimum. There is no limitation to the number of tissues the user can configure in their simulation.

Apollo also recognises five roles played by the tissues. They are as follows:

A. Viral entry tissues
B. Infectious load tissues
C. Terminal load tissues
D. Viral exit tissues
E. Sampling tissues (Optional, dependent on sampling being activated)

We will look at each of these roles in detail under their respective parameter declaration. Each role has a minimum of one tissue and a can have a maximum of a combination of tissues.

#### A. Nesting parameters

Now it is a good time to also introduce Apollo's nested parameter structure. For certain parameters such as tissue configuration, the main parameter can spawn subordinate parameters. For instance, each tissue will have their own set of properties, therefore, to contain each tissues parameters we can use nesting in the JSON script itself. We will explain this further using the provided example in **Figure S - 3**.

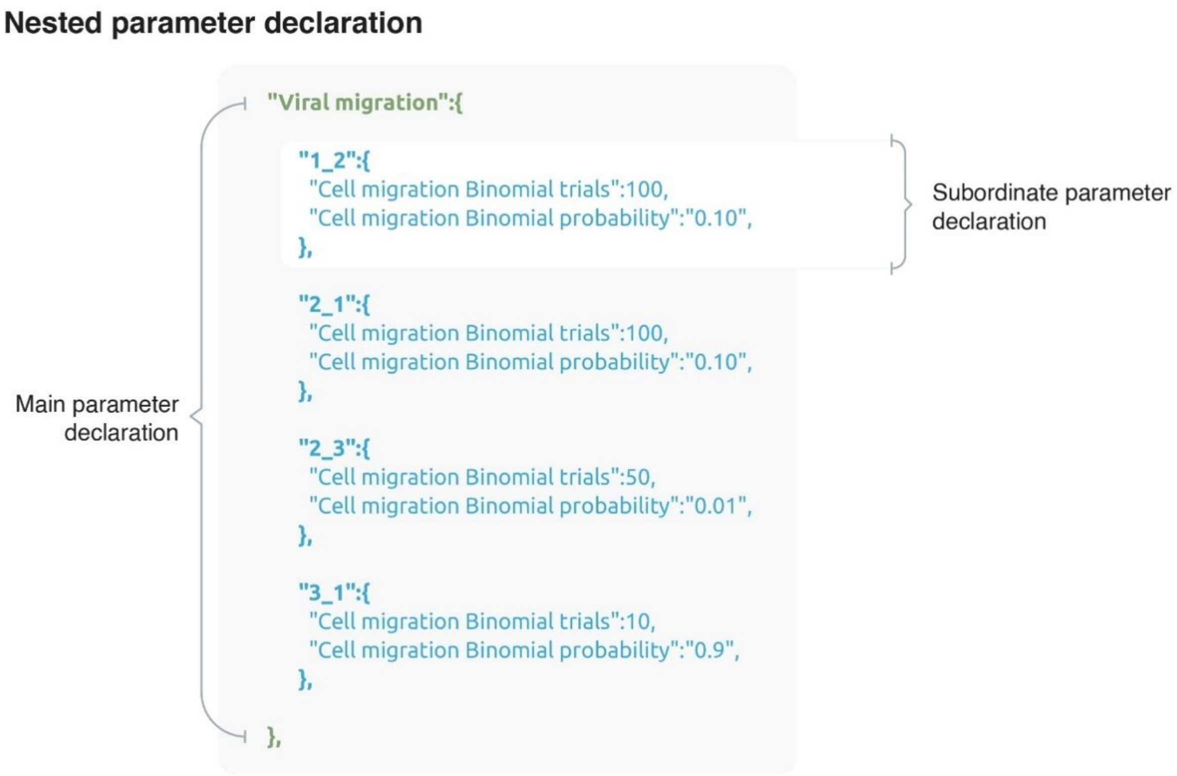

**Figure S - 3:** Example of a nested parameter declaration. As shown "Viral migration" is the main parameter declaration. The user has then within its scope declared the different tissue to tissue migrations that occur. For instance, in the first subordinate parameter declaration viral particles migrate from Tissue 1 to Tissue 2 (indicated by 1_2) under a binomial distribution ($\mathbf{100}, \mathbf{0}.\mathbf{10}$).

As shown in the **Figure S - 3** the main parameter ("Viral migration") is first declared and following the colon (:) we open curly braces ({) to define the scope of the parameter. Any parameter that falls between the curly braces ({}) of the main parameter are subordinate parameters of that parameter. By the end of this 3.3.6 sub section we will learn how to declare such parameters.

**3.3.6.1. Tissue profiles**

We start the declaration of tissues using this parameter. It is a main parameter, therefore once declared it should be followed by the opening of curly braces. All properties that belong to the tissues must be nested under the parameter Tissue profiles.

E.g.: *"Tissue profiles":{…………}*

**3.3.6.2. Number of tissues**

Here the user declares the number of tissues ($T$) that they wish to define.

E.g.: *"Number of tissues":3*

**3.3.6.3. Tissue (*t*) Name**

This parameter is used to declare the names of each tissue. We replace ($t$) by the tissue number.

E.g.: For instance, if we have set the above parameter (3.3.6.2) for the number of tissues as 3, then we define the names of the 3 tissues as follows:

*"Tissue 1 Name":"Blood",*

*"Tissue 2 Name":"Lung",*

*"Tissue 3 Name":"Gut"*

**3.3.6.4. Viral entry tissues**

These are the tissues that are available for viral entry. During the incidence of viral infection, these tissues will be the ones to receive the incoming viral particles. The tissues are listed using their numerical reference and the tissues are separated by a comma (,).

E.g.: In this example two tissues are available as entry tissues. So, when the viral particles infect the susceptible host, it will enter these tissues.

*"Viral entry tissues":"1,2"*

###### 3.3.6.5. Infectious load tissues

The summation of the viral particles in these tissues will contribute to a host becoming infectious.

E.g.: The summation of the viral loads in tissues 1 and 3 will contribute to a host becoming infectious.

*"Infectious load tissues":"1,3"*

###### 3.3.6.6. Terminal load tissues

The summation of the viral particles in this tissue will contribute to a host reaching mortality (dead).

E.g.: *"Terminal load tissues":"1,2,3"*

###### 3.3.6.7. Viral exit tissues

These are the tissues that are available for viral exit. During the incidence of viral infection, these tissues from which the viral particles will exit to infect the new host.

E.g.: *"Viral exit tissues":"1"*

###### 3.3.6.8. Sampling tissues

These are the tissues that are available for sampling. Viral particles from these tissues will be sampled. This variable is dependent on sampling being activated (3.3.4.1).

E.g.: *"Sampling tissues":"1,2,3"*

###### 3.3.6.9. Viral tissue migration

This parameter activates the migration of viral particles. It is a Boolean variable of either “YES” or “NO”.

E.g.: *"Viral tissue migration":"Yes"*

###### 3.3.6.10. Viral migration

This is the declaration of the viral migration parameter. This is a main parameter that holds subordinate parameters.

E.g.: *"Viral migration":{…………}*

#### 3.3.6.11. $N_1$_$N_2$

This is a subordinate parameter of Viral migration (3.3.6.10) that is used to define the rate of viral particle migration from one tissue to the next. The configuration is done via a binomial distribution.

E.g.: To configure the transmission of viral particles from tissue 1 to tissue 2, we declare:

*"1_2":{*

*"Cell migration Binomial trials":100,*

*"Cell migration Binomial probability":"0.10",*

*}*

###### 3.3.6.12. Cell migration Binomial trials

This is a subordinate parameter of the above parameter 3.3.6.11. This parameter defines the number of trials ($n$) in the binomial distribution for cell migration.

E.g.: *"Cell migration Binomial trials":100*

###### 3.3.6.13. Cell migration Binomial probability

This is a subordinate parameter of the above parameter 3.3.6.11. This parameter defines the probability of success ($p$) in the binomial distribution for cell migration.

E.g.: *"Cell migration Binomial probability":"0.10"*

##### 3.3.6.14. Overview of tissue declaration

We will take a quick look at an example for the declaration of tissues with **Figure S - 4** below.

tissue declaration and configuration

```
"Tissue profiles":{

  "Number of tissues":3,

  "Tissue 1 Name":"Blood",
  "Tissue 2 Name":"Lung",
  "Tissue 3 Name":"Gut",

  "Viral entry tissues":"1,2",
  "Infectious load tissues":"1",
  "Terminal load tissues":"1,2,3",
  "Viral exit tissues":"1",
  "Sampling tissues":"1,2,3",

  "Viral tissue migration":"Yes",

  "Viral migration":{

    "1_2":{
      "Cell migration Binomial trials":100,
      "Cell migration Binomial probability":"0.10",
    },

    "2_1":{
      "Cell migration Binomial trials":100,
      "Cell migration Binomial probability":"0.10",
    },

    "2_3":{
      "Cell migration Binomial trials":50,
      "Cell migration Binomial probability":"0.01",
    },

    "3_1":{
      "Cell migration Binomial trials":10,
      "Cell migration Binomial probability":"0.9",
    },
  },
}
```

Number of tissues

Tissue names

Functional roles

Status of cell migration

Inter tissue viral migration

**Figure S - 4:** Code snippet for declaration of three different tissue types, complete with their names, functional roles, and rates of viral migration between the tissues.

In the example **Figure S - 4** the user has configured three different types of tissues named Blood, Lung, and Gut. They have assigned the roles of the tissues, with certain roles such as Viral entry having multiple tissues. Viral migration between tissues have also been activated and the rates of the migrations have been configured via the binomial distributions.

##### 3.3.7. Node/Host Profiles

Apollo supports heterogeneous nodes in its networks. This means that no two nodes in the population needs to be the same and have differentiating attributes between them. To state a few per host configurations that can be performed users can define the respective tissue's affinity to viral particles, the number of cells available for infection per tissue and limits of viral loads in terms of infectivity, and mortality.

Now we will look at the different parameters that are associated with the configuration of a node profile and how to create multiple profiles within the same simulation to be distributed across the contact network.

###### 3.3.7.1. Number of node profiles

First and foremost, lets define the number of node profiles that will be present through this parameter.

E.g.: *"Number of node profiles":3*

###### 3.3.7.2. Location of node profiles

The parameter files per each node profile is sperate from the main Node/ Host parameter file. They can be defined and then stored in a central folder whose location is given in the main Node/ Host parameter file. The next set of parameters will involve the configuration of these host profile files.

It should be noted that profile file should be named as $\text{profile\_}h$, $h$ being the profile number. So, if you have $H$ total profiles as defined by the above parameter 3.3.7.1, then $\text{profile\_}h$ will be: $\text{profile\_}h \in \{\text{profile\_1}, \text{profile\_2}, \text{profile\_3}, \ldots, \text{profile\_}H\}$.

E.g.: *"Location of node profiles":"/individual_Profiles/all_Profiles"*

###### 3.3.7.3. Profile name (*profile_h file parameter*)

States the host's group name for the profile.

E.g.: *"Profile name":"Adults Medicated"*

###### 3.3.7.4. Probability of occurrence (*profile_h file parameter*)

The probability ($h$) of this host profile being present in the contact network ($h \in [0,1]$). The combined probabilities of the different profiles ($p_h$) should add up to 1.

$$p_h \in [0,1] \text{ where } h = \{1,2,3,\ldots,H\}$$
$$\sum_{h=1}^{H} p_h = 1$$

E.g.: *"Probability of occurence":"0.75"*

###### 3.3.7.5. Infectious load distribution type (*profile_h file parameter*)

This parameter works in unison with the aforementioned parameter 3.3.6.5. It will define the distribution used to parametrize the minimum viral particle count that will make the infected host infectious and be viable to transmit the disease from one host to the next. This parameter can take either one of two values, "Binomial" for a binomial distribution to determine the number of viral particles required to become infectious or "Fixed" where it is always a fixed number of viral particles for that host type. It can be understood by using the binomial parametrisation it can be ensured that even though two hosts belong to the same profile in terms of their infectivity they can be different.

It should also be noted that manipulation of this parameter can enable the user to change the compartment model of the network from a SIR (Susceptible Infectious Recovered/ Removed) model to an SEIR (Susceptible Exposed Infectious Recovered/ Removed) model. Intuitively we can see that if the infectious load is set to one then it will be a SIR model where an infected person will immediately become infectious. Whereas if we set it to a value greater than one the viral load will have to build up inside the host (Exposed phase) before becoming infectious.

E.g.: *"Infectious load distribution type":"Fixed"*

###### 3.3.7.6. Infectious load binomial trials (*profile_h file parameter*)

The number of trials (n) in the binomial distribution for determining the infectious load when parameter 3.3.7.5 is set to “Binomial”.

E.g.: *"Infectious load Binomial trials":1000*

###### 3.3.7.7. Infectious load Binomial probability (*profile_h file parameter*)

The probability of success (p) in the binomial distribution for determining the infectious load when parameter 3.3.7.5 is set to “Binomial”.

E.g.: *"Infectious load Binomial probability":"0.75"*

###### 3.3.7.8. Infectious load fixed (*profile_h file parameter*)

The exact fixed viral load for the infectious load when parameter 3.3.7.5 is set to “Fixed”. If this value is set to one, the model will behave as an SIR compartment model for that host type. If you set it across all host profiles the entire simulation will behave as a SIR compartment model else it will follow a SEIR model.

E.g.*: "Infectious load Fixed":1*

###### 3.3.7.9. Terminal load distribution type (*profile_h file parameter*)

This parameter works in unison with the aforementioned parameter 3.3.6.6 to determine the minimum number of viral particles required for an infected host to reach mortality (termination of infected host). Similar to the infectious load this parameter can take either one of two values, “Binomial” for a binomial distribution to determine the number of viral particles required to reach mortality or “Fixed” where it is always a fixed number of viral particles for that host type.

E.g.: *"Terminal load distribution type":"Binomial"*

###### 3.3.7.10. Terminal load Binomial trials (*profile_h file parameter*)

The number of trials ($n$) in the binomial distribution for determining the terminal load when parameter 3.3.7.9 is set to “Binomial”.

E.g.: *"Terminal load Binomial trials":100000*

###### 3.3.7.11. Terminal load Binomial probability (*profile_h file parameter*)

The probability of success ($p$) in the binomial distribution for determining the terminal load when parameter 3.3.7.9 is set to “Binomial”.

E.g.: *"Terminal load Binomial probability":"0.75"*

###### 3.3.7.12. Terminal load Fixed (*profile_h file parameter*)

The exact fixed viral load for the terminal load when parameter 3.3.7.9 is set to “Fixed”.

E.g.: *"Terminal load Fixed":1000*

###### 3.3.7.13. Infection sequence distribution (*profile_h file parameter*)

Used to determine the number of individuals that will be potentially infected at once. The parameter helps configure the infectivity of individuals and can take either one of two values, “Binomial” for a binomial distribution to determine the number of hosts being infected or “Fixed” where it is always a fixed number of hosts.

E.g.: *"Infection sequence distribution":"Fixed"*

###### 3.3.7.14. Infection sequence Binomial trials (*profile_h file parameter*)

The number of trials ($n$) in the binomial distribution for determining the number of hosts being infected when parameter 3.3.7.13 is set to “Binomial”.

E.g.: *"Infection sequence Binomial trials":2*

###### 3.3.7.15. Infection sequence Binomial probability (*profile_h file parameter*)

The probability of success (p) for the binomial distribution for determining the number of hosts being infected when parameter 3.3.7.13 is set to "Binomial".

E.g.: *"Infection sequence Binomial probability":"0.5"*

###### 3.3.7.16. Infection sequence Fixed (*profile_h file parameter*)

The exact number of hosts being infected when parameter 3.3.7.13 is set to "Fixed".

E.g.: *"Infection sequence Fixed":2*

###### 3.3.7.17. Sampling effect (*profile_h file parameter*)

In certain epidemiological models sampling can effect the infectivity of a host. For some sampling is the equivalent of an individual getting removed from the infectious population, in some sampling can reduce the infectivity of an individual and in others sampling can have no effect all together.

To capture these dynamics, we use the sampling effect parameter. It can take either one of three values. "No change" meaning sampling will have no effect on the infectivity of an individual, "Less infectious" meaning the infectivity of the individual will drop based on a beta distribution and finally "Removed" where sampled individuals will cease to be infectious. Apollo enables the user to control it at the host profile level.

E.g.: *"Sampling effect":"No change"*

###### 3.3.7.18. Sampling less infectious effect Alpha (*profile_h file parameter*)

This parameter defined the alpha ($\alpha$) of the beta distribution for when the sampling effect parameter 3.3.7.18 is set to "Less infectious".

E.g.: *"Sampling less infectious effect Alpha":"5"*

###### 3.3.7.19. Sampling less infectious effect Beta (*profile_h file parameter*)

This parameter defined the beta (β) of the beta distribution for when the sampling effect parameter 3.3.7.18 is set to "Less infectious".

E.g.: *"Sampling less infectious effect Beta":"10"*

###### 3.3.7.20. Tissue profiles (*profile_h file parameter*)

This is a call to the same parameter from 3.3.6.1. However, inside the profile parameter files we use a separated set of subordinate parameters. In the profile parameter file, we will configure the tissue environment per host profile. This will include the possible cell concentrations that may be present per tissue structure, the phases of infection that the tissues may experience to the affinity of different tissue cell types for viral attachment. Here we move from the tissue level into the cellular level.

###### 3.3.7.21. Tissue ($t$) (*profile_h file parameter*)

This is the subordinate function we use to parameterise each individual tissue type per host profile. We replace the ($t$) with the tissue number that we wish to configure.

E.g.: *"Tissue 1":{ …………}*

###### 3.3.7.22. Cell limit (*profile_h file parameter*)

For some tissues types the user may want to configure a limit for the number of cells that are available for viral infection. In such instances this parameter can be activated. It is Boolean variable of either "Yes" to activate the cell limit or "No" to enable an unlimited number of cells available for infection.

E.g.: "*Cell limit":"YES"*

###### 3.3.7.23. Cell limit Binomial trials (*profile_h file parameter*)

The number of cells available for infection is parametrised via a binomial distribution. When parameter 3.3.7.22 is set to "YES" this parameter can be used to define the number of trials ($\mathrm{n}$) if the binomial distribution.

E.g.: *"Cell limit Binomial trials":100000*

###### 3.3.7.24. Cell limit Binomial probability (*profile_h file parameter*)

Is used to parametrise the probability of success ($\mathrm{p}$) for the binomial distribution when parameter 3.3.7.22 is set to "YES".

E.g.: *"Cell limit Binomial probability":"0.50"*

###### 3.3.7.25. Viral distribution type (*profile_h file parameter*)

This parameter set is used to control the affinity of a viral particle to a cell. This can vary based on the tissue type. It helps determine how many viral particles will attach to each cell during the viruses replication cycle. It can take the form of either a binomial distribution activated by setting this parameter to "Binomial" or as a gamma distribution activated by setting this parameter to "Gamma".

E.g.: *"Viral distribution type":"Binomial"*

###### 3.3.7.26. Viral distribution Binomial trials (*profile_h file parameter*)

This parameter defines the number of trials ($\mathrm{n}$) of the binomial distribution for viral affinity during cell attachment when the above parameter 3.3.7.25 is set to "Binomial".

E.g.: *"Viral distribution Binomial trials":10*

###### 3.3.7.27. Viral distribution Binomial probability (*profile_h file parameter*)

This parameter defines the probability ($\mathrm{p}$) of the binomial distribution for viral affinity during cell attachment when the above parameter 3.3.7.25 is set to "Binomial".

E.g.: *"Viral distribution Binomial probability":"0.5"*

###### 3.3.7.28. Viral distribution Gamma shape (*profile_h file parameter*)

This parameter defines the shape ($k$) of the gamma distribution for viral affinity during cell attachment when the above parameter 3.3.7.25 is set to "Gamma".

E.g.: *"Viral distribution Gamma shape":"3"*

###### 3.3.7.29. Viral distribution Gamma scale (*profile_h file parameter*)

This parameter defines the scale (θ) of the gamma distribution for viral affinity during cell attachment when the above parameter 3.3.7.25 is set to "Gamma".

E.g.: *"Viral distribution Gamma scale":"3.33"*

###### 3.3.7.30. Replication phases (*profile_h file parameter*)

During the course of infection, the host environment can be subjected to various pressures. The replication phases parameter helps to capture these dynamics to a certain degree.

For instance, during the initial incidence of viral infection there is usually little to no interference from the host environment to the viral life cycle, we call this a "Neutral" phase, where the viral particles are free to replicate at will.

Another instance experienced by viruses are during the later stages of infection when the host immune system attempts to counteract the invading organism. At a certain point a balance between the invading virus and the host immune system is reached. In such instances there is observed a stationary viral load in the organism, we call this the "Stationary" phase. This phase can also be used to simulate models of constant population size such as the Wrigh Fisher model, where a constant population size is maintained from one generation to the next.

Finally, during a successful immune response or drug therapy the viral load can be diminished from the host. We have dubbed this as the "Depreciation" phase where the viral load is gradually be reduced from one generation to the next along a beta distribution.

These are the phases that can be captured by the Replication phases parameter. It too is a subordinate parameter and falls under the Tissue ($t$) 3.3.7.21 parameter. To provide a further layer of granularity to the user this is a tissue specific mechanic, meaning that individual tissues can separate phases of infection. This is quite ideal for configuring tissue dynamics such as reservoir tissues that usually contain viral loads that are might be unaffected by drug therapy, or to simulate a situation where a cured infected individual becomes infectious again.

E.g.: *"Replication phases":{…………}*

**3.3.7.31. Number of phases (*profile_h file parameter*)**

This parameter is used to define the number of phases ($X$) that the tissue can have. The user can define any combination of the above mentioned three phase types and have a tissue undergo multiple counts of “Neutral”, “Stationary” and “Depreciation” phases.

E.g.: *"Number of phases":4*

**3.3.7.32. Phase ($x$) Mode (*profile_h file parameter*)**

This is used to indicate the mode of a particular phase. The ($x$) is replaced by the phase being configured ($x \in \{1,2,3 \ldots, X\}$).

E.g.: *"Phase 1 Mode":"Neutral"*

**3.3.7.33. Phase ($x$) Time ratio (*profile_h file parameter*)**

This is used to indicate the ratio of time ($t_x$) the tissue spends in this phase from the total coarse of infection for that particular host. The ($x$) is replaced by the phase being configured. The total time ($T$) should add up to one for that particular tissue.

$$t_x \in [0,1] \text{ and } x \in \{1,2,3 \ldots, X\}$$

$$\sum_{x=1}^{X} t_x = 1$$

###### 3.3.7.34. Phase ($x$) Variance (*profile_h file parameter*)

This parameter is activated when Phase ($x$) Mode (3.3.7.32) is set to "Stationary". It allows a variance to exist around the stationary count number of the viral load based on the time the phase goes into effect. If a variance is not needed it can be simply set to zero (0).

E.g.: *Phase 1 Variance":"100"*

###### 3.3.7.35. Phase ($x$) Alpha (*profile_h file parameter*)

This parameter is activated when Phase ($x$) Mode (3.3.7.32) is set to "Depreciation". It defines the alpha ($\alpha$) of the beta distribution which controls the depreciation ratio of the viral load from one generation to the next in the tissue.

E.g.: *"Phase 3 Alpha":"100"*

###### 3.3.7.36. Phase ($x$) Beta (*profile_h file parameter*)

This parameter is activated when Phase ($x$) Mode (3.3.7.32) is set to "Depreciation". It defines the beta (β) of the beta distribution which controls the depreciation ratio of the viral load from one generation to the next in the tissue.

E.g.: *"Phase 3 Beta":"100"*

###### 3.3.7.37. Overview of host profile parameter file

With that we complete the parameters (3.3.7.3 to 3.3.7.36) for the profile parameter files. The provided example in **Figure S - 5** shows a completed profile parameter file. The file shows complete parameterisation of the host profile labelled "Adults" who has a 75% probability of populating the contact network. It set up as an SIR model with infectious load being fixed. The transmission bottleneck has been confined to two viral particles per transmission event and one tissue structure. The tissue experiences 3 separate phase events beginning with a neutral phase, followed by stationary and depreciation phases. There is no cell limit in the tissue, so the viral particles can attach to an unlimited number of cells. It should also be noted that the profile is affected by sampling. If the host is sampled its probability of infecting another host drops.

profile_1.json

```
{
  "Profile name":"Aults",

  "Probability of occurence":"0.75",

  # Sampled individuals can be REMOVED from the infectious population or become LESS INFECTIOUS
    or NO CHANGE to infectivity
  "Sampling effect":"No change",
  "Sampling less infectious effect Alpha":"5",
  "Sampling less infectious effect Beta":"10",

  # INFECTIOUS load
  "Infectious load distribution type":"Fixed",
  "Infectious load Binomial trials":100,
  "Infectious load Binomial probability":"0.75",
  "Infectious load Fixed":1,

  # TERMINAL load
  "Terminal load distribution type":"Fixed",
  "Terminal load Binomial trials":100000,
  "Terminal load Binomial probability":"0.75",
  "Terminal load Fixed":5,

  # Control the number of hosts that will be infected
  "Infection sequence Fixed":2,

  "Tissue profiles":{

    "Tissue 1":{
      "Cell limit":"NO",

      # Number of viral particles that will attach to a cell
      "Viral distribution type":"Binomial",
      "Viral distribution Binomial trials":10,
      "Viral distribution Binomial probability":"0.5"

      "Replication phases":{

        "Number of phases":3,

        "Phase 1 Mode":"Neutral",
        "Phase 1 Time ratio":"0.2",

        "Phase 2 Mode":"Stationary",
        "Phase 2 Time ratio":"0.5",
        "Phase 2 Variance":"100",

        "Phase 3 Mode":"Depriciation",
        "Phase 3 Time ratio":"0.3",
        "Phase 3 Alpha":"100",
        "Phase 3 Beta":"100",
      },
    },
  },
}
```

**Figure S - 5:** Sample host profile parameter file. The file is for the configuration of the first host profile (profile_1.json) labelled as "Adults". The file configures the probability of the profile type occurring in the contact network and completes the tissue and cellular structures of the host. It also shows the configuration of host traits such as effect of sampling phases the tissue undergoes.

##### 3.3.8. Overview of the Node/ Host parameter file

With this we complete the parameterisation of the Node/ Host parameter file. The provided example **Figure S - 6** shows the parametrisation of a host parameter population where each host has three tissues associated with the viral life cycle.

Node_parameter_file.json

```
{
  # In days the time taken for viral replication, from cell entry to progeny generation
  "Shape replication time":"20.0",
  "Scale replication time":"0.020",

  # In days the time the virus will spend in the host, if unaffected
  "Shape days in host":"20.0",
  "Scale days in host":"0.27",

  "Sampling present":"Yes",

  # If Resampling is YES, then the same node can be sampled multiple times as long as it is not removed.
  "Resampling of nodes":"Yes",

  "Sampling rate Binomial trials":2,
  "Sampling rate Binomial probability":"0.75",

  # The number of sequences sampled from each sampling tissue
  "Distribution per node sequences sampled":"FIXED",
  "Per node sampling rate Binomial trials":2,
  "Per node sampling rate Binomial probability":"0.5",
  "Per node sampling rate Fixed":1,

  # Control if the simulation ends via the max sampled sequences or not
  "Limit samples obtained":"Yes",
  "Max samples to obtain":20,

  # Number of progeny that will be generated by a reference virulant particle
  # Negative Binomial, Gamma, Poisson
  "Progeny distribution type":"Negative Binomial",

  "Progeny Negative Binomial sucesses":10,
  "Progeny Negative Binomial probability":"0.55",

  "Progeny Poisson mean":"10",

  "Progeny Gamma shape":"10",
  "Progeny Gamma scale":"0.5",

  # Infected individuals can remain suceptible to infection if set to YES, else they will be removed if their viral load goes to zero
  "Infected to Recovered":"No",

  "Number of node profiles":3,
  "Location of node profiles":"/individual_Profiles/all_Profiles",

  "Tissue profiles":{

    "Number of tissues":3,

    "Tissue 1 Name":"Blood",
    "Tissue 2 Name":"Lung",
    "Tissue 3 Name":"Gut",

    "Viral entry tissues":"1,2",
    "Infectious load tissues":"1",
    "Terminal load tissues":"1,2,3",
    "Viral exit tissues":"1",
    "Sampling tissues":"1,2,3",

    "Viral tissue migration":"Yes",

    "Viral migration":{

      "1_2":{
        "Cell migration Binomial trials":100,
        "Cell migration Binomial probability":"0.10",
      },

      "2_1":{
        "Cell migration Binomial trials":100,
        "Cell migration Binomial probability":"0.10",
      },

      "2_3":{
        "Cell migration Binomial trials":50,
        "Cell migration Binomial probability":"0.01",
      },

      "3_1":{
        "Cell migration Binomial trials":10,
        "Cell migration Binomial probability":"0.9",
      },

    },
}
```

**Figure S - 6:** Complete Node/ Host parameter file with tissue structure declaration. The file shows the allocation of the generation time of the virus within the host as well as the average time spent within the host by the viral load. The sampling of infected individuals has also been configured complete with the parameters for the roles of the tissues.

##### 3.4. Sequence parameter file

The sequence parameter file is used to configure the dynamics of the viral sequence. Apollo is equipped to capture a series of mechanisms such as mutation, proof reading, recombination, fitness, survivability and even how these factors will change with evolving viral strains. In this section through the configuration of the sequence parameter file we will see how each of there dynamics can be captured.

###### 3.4.1. Parent sequences

The parent sequences are the set of the sequences the initial host (patient zero) of the simulation gets infected with. This set of parent sequences could be a diverse set of sequence or even multiple copies of the same sequence, it can also be just one sequence. This will be the first invading viral population.

It should be noted that the parent sequences should all be of the same length, and this will be the genome length of the virus. The genome length parameter is automatically configured by Apollo using the parent sequences.

###### 3.4.1.1. Parent sequences folder

Specifies the location of the folder that contains the parent sequences. The sequences can be present in one single FASTA file or be in multiple individual FASTA files or a combination of the two.

E.g.: *"Parent sequences folder":"/reference_Genomes"*

###### 3.4.2. Reference parameters

These are the parameters that the reference genome will contain. These parameters can be subject to change based on the mutation of the viral genome and the magnitude of these changes can be configured via different profiles. The main reference parameters that apply to the whole genome are Fitness, Survivability, and where available Proof Reading. They are parametrised using three parameters.

###### 3.4.2.1. Reference Fitness

Configures the reference sequences fitness ($f$). In Apollo fitness is related to the viral progeny generation. If a strain has a fitness of one, then it will produce progeny at the rate defined by the parameters from 3.3.3.1 to 3.3.3.6. If lets say a strain has an overall fitness of two it will produce at twice the rate and if it has an overall fitness of 0.5 it will produce progeny at half the rate and so on. This can take a value from 0 to infinity ($f \in [0, \infty]$).

E.g.: *"Reference Fitness":"1"*

###### 3.4.2.2. Reference Survivability

Configures the reference sequences survivability ($s$). This is the probability of a viral particle surviving till progeny generation. It can take a value between 0 and 1 ($s \in [0,1]$).

E.g.: *"Reference Survivability":"0.5"*

###### 3.4.2.3. Proof reading availability

Specifies if the viral genome has a proof-reading mechanism, where the mismatch errors during replication are mitigated. It is a Boolean variable with values being "YES" if proof reading is available an "NO" to disable the proof-reading mechanism.

E.g.: *"Proof reading availability":"Yes"*

###### 3.4.2.4. Reference Proof Reading

Configures the reference sequences proof reading mechanisms accuracy ($r$) when proof reading is made available by setting parameter 3.4.2.3 to "YES". It is a probability value specifying the probability of the mechanism catching a mis match during replication. It can take a value between 0 and 1 ($r \in [0,1]$).

E.g.: *"Reference Proof Reading":"0.75"*

##### 3.4.3. Sequence profiles

For the standard reference parameters (3.4.2) there are fitness profiles, these profiles state the changes that the parameter will undergo based on the base occupied by various sites in the viral genome of the viral particle. These sites can be intuitively understood to be the segregating sites of the viral genome.

The sequence profiles are separate files that take the form of CSV (Comma separated values) files. For Apollo these files can be tab or comma delimited. Apollo will automatically detect the character used for delimitation. The file takes a form of a matrix ($P$). The rows ($R$) are fixed at four for each base (A, T, G and C) and the columns ($C$) refer to the position of each setting site in the genome. Apollo uses a nonzero start index meaning the first base position in the genome is one. It should be noted that these matrixes do not need to be configured in order. As shown in **Table S - 2** the columns are not ordered. Similarly, the rows do not need to follow the A, T, G, C order.

If the sequence profile file is not configured for a parameter this would be interpreted as that particular reference parameter does not undergo changes based on the diversity of the viral genome. In such an instance the relevant parameter below from 3.4.3.1 to 3.4.3.3 should be set to "NA".

###### 3.4.3.1. Fitness profile file

We use this parameter to specify the location of the fitness profile parameterisation file. The values of the matrix ($F$) of the fitness profile file can take any value between zero and infinity ($f_{ij} \in [0, \infty]$). The overall fitness of the strain will be calculated as the sum product of the values. We will look at an example of how to parameterize the file via **Table S - 2**.

E.g.: *"Fitness profile file":"/sequence_Profiles/profiles/sequence_Fitness_Profile.csv"*

**Table S - 2:** Table depicting the fitness profile values that will be undertaken based on the base being occupied at different segregating sites. The rows refer to the corresponding base and the columns refer to the genome position of the site.

| | **150** | **300** | **450** | **320** |
|---|---|---|---|---|
| **A** | 1 | 1 | 1 | 1 |
| **T** | 2 | 1 | 1 | 1.5 |
| **G** | 1 | 1 | 0.25 | 1 |
| **C** | 1 | 0.5 | 1 | 1 |

###### 3.4.3.2. Survivability profile file

We use this parameter to specify the location of the survivability profile parameterisation file. The values of the matrix ($S$) of the survivability profile file can take any value between negative one and positive one ($s_{ij} \in [-1,1]$). Apollo has the positive to negative range because the overall survivability of the strain will be calculated as the summation of the values. The negative values help reduce the survivability probability and the positive values to increase the survivability. We will look at an example of how to parameterize the file via **Table S - 3**.

E.g.: *"Survivability profile file":"/sequence_Profiles/profiles/sequence_Survivability_Profile.csv"*

**Table S - 3:** Table depicting the fitness profile values that will be undertaken based on the base being occupied at different segregating sites. The rows refer to the corresponding base and the columns refer to the genome position of the site.

| | **150** | **450** | **500** | **1200** | **1800** |
|---|---|---|---|---|---|
| **A** | 0 | 0.25 | 0 | 0 | -0.25 |
| **T** | -0.05 | 0 | 0 | 0.002 | 0 |
| **G** | 0 | 0 | 0.05 | 0 | 1 |
| **C** | 0 | 0 | -0.03 | 0 | -0.005 |

###### 3.4.3.3. Proof reading profile file

We use this parameter to specify the location of the proof-reading profile parameterisation file. Similar to the survivability matrix values of the proof-reading matrix ($R$) can take any value between negative one and positive one ($r_{ij} \in [-1,1]$). We will look at an example of how to parameterize the file via **Table S - 4**.

E.g.: *" Proof reading profile file":"/profiles/sequence_Proof_Reading_Profile.csv"*

**Table S - 4:** Table depicting the proof-reading profile values that will be undertaken based on the base being occupied at different segregating sites. The rows refer to the corresponding base and the columns refer to the genome position of the site.

| | **150** | **300** | **450** | **1200** |
|---|---|---|---|---|
| **A** | -0.25 | 0 | 0 | 0.05 |
| **T** | 0 | 0 | 0 | -0.25 |
| **G** | 0 | -0.5 | 0.002 | -0.75 |
| **C** | 0.25 | 0.5 | 0 | 0.25 |

###### 3.4.4. Transmission of viral particles

These parameters control the transmission of viral particles from one host to the next. An important component of host-to-host viral transmission is the transmission bottleneck. This is the factor that accounts for the viral load that gets transmitted between hosts. Apollo enables the user to control the number of viral particles being transmitted during the incidence of disease infection from an infectious host.

###### 3.4.4.1. Reinfection availability

This parameter enables the control of already infected hosts being reinfected by other infectious individuals in their vicinity of the contact network. This is a Boolean variable that can take the values of either "YES" to enable reinfection of already infected hosts or "NO" to disable the mechanic and allow infection of only susceptible individuals.

E.g.: *"Reinfection availability":"No"*

###### 3.4.4.2. Transmitted sequence distribution

This parameter controls the distribution that will govern the viral transmission bottleneck. Apollo allows the distribution parameter to be either "Binomial" for a binomial distribution or "Fixed" for only a fixed number of viral particles to be transmitted at any given time.

E.g.: *"Transmitted sequence distribution":"Fixed"*

###### 3.4.4.3. Transmitted sequence Binomial trials

This parameter defines the number of trials ($n$) in the binomial distribution for the transmission bottleneck when the above parameter 3.4.4.2 is set to "Binomial."

E.g.: *"Transmitted sequence Binomial trials":2*

###### 3.4.4.4. Transmitted sequence Binomial probability

This parameter defines the probability ($p$) in the binomial distribution for the transmission bottleneck when the above parameter 3.4.4.2 is set to "Binomial".

E.g.: *" "Transmitted sequence Binomial probability":"0.5"*

###### 3.4.4.5. Transmitted sequence Fixed

This parameter set the number of viral particles transmitted between hosts at a fixed value when the above parameter 3.4.4.2 is set to “Fixed”.

E.g.: *"Transmitted sequence Fixed":2*

###### 3.4.5. Mutation parameterization

Mutations are an important aspect that governs the trajectory of a virus through an epidemic. An important aspect of capturing mutations dynamics is their effects. Which we have done via our sequence profiles in the previous section 3.4.3. Now we will look at how to parameterize the very act of mutation. This includes the identification of regions of the genomes that undergo mutations, which in Apollo are called “mutation hotspots”. These hotspots behave independently to each other and can be as simple as a single base position or span multiple bases with each hotspot having varying rates of mutation.

With that let’s look at the configuration of the mutation parameters.

###### 3.4.5.1. Mutation availability

This parameter activates the mutation mechanic in Apollo. It is a Boolean variable with values being either “YES” to enable mutation of the genome or “NO” to deactivate the mutation mechanic.

E.g.: *"Mutation availability":"Yes"*

###### 3.4.5.2. Mutations

This is a main parameter declaration. We have to open the “Mutations” parameter and within its scope declare the subordinate parameters that govern the mutation of the genome along the course of viral infection in the epidemic. All subsequent parameters fall inside the scope of the “Mutations” parameter.

E.g.: *"Mutations":{………}*

###### 3.4.5.3. Number of hotspots

Defines the number of mutation hotspots ($M$) present in the genome.

E.g.: *"Number of hotspots":2*

###### *3.4.5.4.* Hotspot ($m$)

This is a subordinate parameter of 3.4.5.2. It is used to define the mutational parameters of the hotpot such as region, the hotspot spans, mutation rate and probabilities of base change. The ($m$) is replaced with the mutation hotspot being configured ($m \in \{1,2,3 \dots, M\}$). All subsequent parameters from 3.4.5.5 to 3.4.5.11 fall under its scope.

E.g.: *"Hotspot 1":{………}*

###### 3.4.5.5. Region

Defines the start and stop position of the hotspot under configuration. Takes the format of: "start_stop" where start is the starting base position of the mutation hotspot and stop is the last base position of the mutational hotspot. Hotspot regions can overlap over each other. If the user requires to configure a singular base position, they can simply give the same value for the start and the stop positions.

E.g.: *"Region":"100_2000"* or for a singular site *"Region":"100_100"*

###### 3.4.5.6. Clock model

This parameter controls the mutation rate of the hotspot. It is used to specify the number of mutational events that will occur in the region during a generation. The mutation rate of the clock model can be defined using either one of three methods. Setting the parameter as "Poisson" to allow the rate to follow a Poisson distribution, "Negative Binomial" to follow a negative binomial distribution or "Fixed" to have a fixed probability of mutation for each base in the hotspot at every generation.

E.g.: *"Clock model":"Poisson"*

###### 3.4.5.7. Clock model Poisson mean

Used to define the mean (λ) of the Poisson distribution when the above parameter 3.4.5.6 is set to "Poisson".

E.g.: *"Clock model Poisson mean":"2"*

###### 3.4.5.8. Clock model Negative Binomial successes

Specify the number of successes ($r$) in the negative binomial distribution when the above parameter 3.4.5.6 is set to “Negative Binomial”.

E.g.: *"Clock model Negative Binomial sucesses":"10"*

###### 3.4.5.9. Clock model Negative Binomial probability

Specify the probability of successes ($p$) in the negative binomial distribution when the above parameter 3.4.5.6 is set to “Negative Binomial”.

E.g.: *"Clock model Negative Binomial probability":"0.5"*

###### 3.4.5.10. Clock model Fixed probability

Specifies the probability of each base in the mutational hotspot undergoing a mutational event. Activated when the above parameter 3.4.5.6 is set to “Fixed”.

E.g.: *"Clock model Fixed probability":"0.5"*

###### 3.4.5.11. OM (Original base to Mutated base)

Here we configure the base mutation probability. The probability that a base will change to another. There must always be 16 OM parameters to cover all possible base mutations combinations (**Figure S - 7C**). The “O” represents the original state of the base and the “M” the mutated base state ($O, M \in \{A, T, G, C\}$). The parameter will quantify the probability of the original base mutating to a particular base.

E.g.:

| | | | |
|---|---|---|---|
| *"AA":"0.25",* | *"GA":"0.25",* | *"TT":"0.25",* | *"CT":"0.25",* |
| *"AG":"0.25",* | *"GG":"0.25",* | *"TC":"0.25",* | *"CC":"0.25",* |
| *"AT":"0.25",* | *"GT":"0.25",* | *"TA":"0.25",* | *“CA":"0.25",* |
| *“AC":"0.25",* | *"GC":"0.25",* | *"TG":"0.25",* | *"CG":"0.25"* |

It should be noted that each base’s mutation probabilities should sum up to 1.

This is because Apollo treats base mutations as continuous time, finite state Markov Chains[5] (**Figure S - 7A**). Therefore, it takes the form of a $4 \times 4$ transition matrix $X$ (**Figure S - 7B**), where $x_{ij}$ is a real number value ($0 \geq x_{ij} \leq 1$) in the matrix where the row $i$ represents the original state and column $j$ represents the mutated state ($for\ i, j\ = A, T, G, C$).

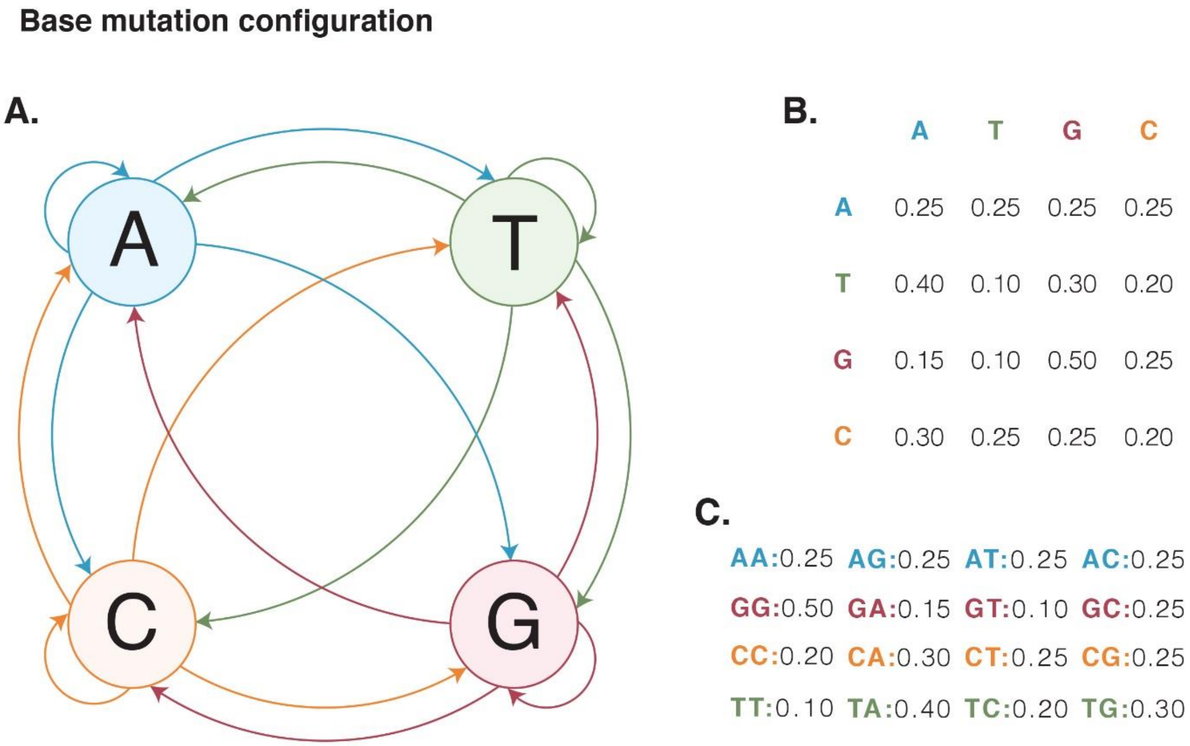

**Figure S - 7:** The DNA base substitution model takes the form of a continuous time finite state Markov chain. (A) Shows the four base states in the Markov chain and their transitions. (B) The matrix representation of the Markov chain, where rows represent the original base state, and the columns represent the mutated state. Each value in the matrix represents the probability of changing from one state to the next. (C) Shows the conversion of the transition matrix to Apollo's OM (Original base to Mutated base) parameter format.

##### 3.4.5.12. Overview of mutations

You may have noticed by now that Apollo's mechanism for mutations, allow epistasis to occur. Where a series of mutations in different regions can have a collective effect on the genome of the viral particle. This is a powerful mechanic that drives evolution, adaptation, and diversity.

Configuring mutation hotspots

```
"Mutations":{

  "Number of hotspots":1,

  "Hotspot 1":{

    "Region":"100_2000",

    # Clock model, can follow a Poisson or Negative Binomial distribution or be Fixed.
    "Clock model":"Poisson",

    "Clock model Poisson mean":"2",

    "AA":"0.25",
    "AG":"0.25",
    "AT":"0.25",
    "AC":"0.25",

    "GA":"0.25",
    "GG":"0.25",
    "GT":"0.25",
    "GC":"0.25",

    "TT":"0.25",
    "TC":"0.25",
    "TA":"0.25",
    "TG":"0.25",

    "CT":"0.25",
    "CC":"0.25",
    "CA":"0.25",
    "CG":"0.25"
  },
}
```

**Figure S - 8:** Sample configuration of a mutation hotspot in the Sequence parameter file. The mutational hotspot region has been defined, including the mutational rate using a Poisson clock model and the probabilities of base transition from one base to the next given the original base state at a position.

###### 3.4.6. Recombination parameterization

Recombination is the process by which genetic information is exchanges between organisms. Essentially regions one organisms genetic code can be exchanged with another through recombination. Recombination has been identified as a driving force of evolution in viruses. Especially that of RNA viruses such as HIV and SARS-CoV-2. It is another evolutionary force that leads to increased genetic diversity.

Apollo comes equipped to capture recombination. It relies on the user configuring regions of the genome that can undergo recombination and then parametrizing these regions based on their likelihood of undergoing recombination as well as their proclivity to be selected over another competitive recombinant gene section. Apollo goes a step further to account for the variations (as a result of mutations or even recombination itself) in the genome to have an affect on these recombination hotspots. This in an extension of Apollo’s ability to account for epistasis (3.4.5.12).

###### 3.4.6.1. Recombination availability

This parameter activates the mutation recombination in Apollo. It is a Boolean variable with values being either “YES” to enable recombination of the genome or “NO” to deactivate the recombination mechanic.

E.g.: *"Recombination availability":"Yes"*

###### 3.4.6.2. Recombination

This is a main parameter declaration for recombination. We have to open the “Recombination” parameter and within its scope declare the subordinate parameters that govern the recombination of the genome along the course of viral infection in the epidemic. All following parameters fall inside the scope of the “Recombination” parameter.

E.g.: *"Recombination":{………}*

###### 3.4.6.3. Number of hotspots

The number of recombination hotspots available ($H$). These are the regions of the genome that will undergo recombination.

E.g.: "*Number of hotspots":3*

###### 3.4.6.4. Hotspot ($\boldsymbol{h}$)

This is a subordinate parameter of 3.4.6.3. It is used to define the parameters of the hotpot such as region the hotspot spans, probability of the hotspot taking art in a recombination event and its affinity to be selected over competing hotspots from other viral particles. The ($h$) is replaced with the recombination hotspot being configured ($h \in \{1,2,3 \dots, H\}$). The following parameters from 3.4.6.5 to 3.4.6.7 fall under this subordinate parameter.

E.g.: *"Hotspot 1":{……}*

###### 3.4.6.5. Region

Similar to the mutational region parameter (3.4.5.5) it defines the start and stop position of the hotspot under configuration.

E.g.: *"Region":"100_200"*

###### 3.4.6.6. Reference probability of recombination

This is the reference genome's probability for recombination for the particular hotspot.

E.g.: *"Reference probability of recombination":"0.5"*

###### 3.4.6.7. Reference selectivity

This is the proclivity of the hotspot with the reference genome's configuration for being selected to form the progeny genome against other viral genomes occupying the same cell.

E.g.: *"Reference selectivity":"1"*

###### 3.4.6.8. Recombination profiles folder

This parameter is outside the scope of 3.4.6.4 but within 3.4.6.2. Each hotspots reference parameters (3.4.6.6 and 3.4.6.7) can be altered via mutations. Therefore, to configure these changes the user can create sequence profile CSV files (**Table S - 5**) for each hotspot similar to the mutation profile files (3.4.3). Each file should be named after the hotspot being configured (E.g.: hotspot_$h.csv$). The file will contain two sections for each parameter, probability of recombination (3.4.6.6 whose overall value will be an additive value similar to 3.4.3.2) and selectivity (3.4.6.7 whose overall value will be calculated as a sum product similar to 3.4.3.1 ).

E.g.:

**Table S - 5:** Sample layout for a recombination profile file for hotspot 1.

| **#hotspot:1** | | | | |
|---|---|---|---|---|
| **Probability** | **150** | **300** | **450** | **1200** |
| **A** | 0 | 0 | 1 | 0 |
| **T** | 0 | 0 | 0 | 0 |
| **G** | -0.25 | 0 | 0.25 | 0 |
| **C** | 0 | 0.5 | 0 | -1 |
| **Selectivity** | **150** | **300** | **450** | **1200** |
| **A** | 1 | 1 | 1 | 1 |
| **T** | 2 | 1 | 1 | 1.5 |
| **G** | 1 | 1 | 0.25 | 1 |
| **C** | 1 | 0.5 | 1 | 1 |

The files should be stored in a singular location and the location can be parameterized using this parameter.

E.g.: *"Recombination profiles folder":"/sequence_Profiles/recombination_profiles"*

##### 3.4.6.9. Overview of recombination

With that we complete the configuration of Apollo's recombination mechanic. Apollo provides a complete capture of recombination in evolution, complete with mechanics to capture the effects of mutation on recombination factors. Through **Figure S - 9** we have an example of how a recombination event can be parameterized in Apollo.

Configuring mutation hotspots

```
"Recombination":{

    "Number of hotspots":3,
    "Recombination profiles folder":"sequence_Profiles/recombination_profiles",

    "Hotspot 1":{
      "Region":"100_200"
      "Reference probability of recombination":"0.5",
      "Reference selectivity":"1",
    },

    "Hotspot 2":{
      "Region":"10_50"
      "Reference probability of recombination":"0.75",
      "Reference selectivity":"1",
    },

    "Hotspot 3":{
      "Region":"10000_20000"
      "Reference probability of recombination":"0.05",
      "Reference selectivity":"1",
    },
  }
```

**Figure S - 9.** Sample configuration of three recombination hotspots complete with their regions, reference probabilities and selectivity for recombination.

##### 3.4.7. Overview of sequence parameter file

With that we complete the parameterisation of the Sequence parameter file. It is evident that Apollo aims to have a complete capture of the evolutionary forces that act on the viral genome during the course of an epidemic.

Sequence_parameter_file.json

```
{
  "Parent sequences folder":"/Simulator_Linux/reference_Genomes",

  "Mutation availability":"Yes",

  "Transmitted sequence distribution":"Fixed",
  "Transmitted sequence Binomial trials":2,
  "Transmitted sequence Binomial probability":"0.5",
  "Transmitted sequence Fixed":2,

  "Reinfection availability":"No",

  "Fitness profile file":"/sequence_Fitness_Profile.csv",
  "Survivability profile file":"/sequence_Survivability_Profile.csv",

  "Proof reading availability":"Yes",
  "Proof reading profile file":"/sequence_Proof_Reading_Profile.csv",

  "Recombination availability":"Yes",

  "Reference Fitness":"1",
  "Reference Survivability":"0.5",
  "Reference Proof Reading":"0.75",

  "Mutations":{

    "Number of hotspots":2,

    "Hotspot 1":{

      "Region":"100_2000",
      "Clock model":"Poisson",

      "Clock model Poisson mean":"2",
      "Clock model Negative Binomial sucesses":"10",
      "Clock model Negative Binomial probability":"0.5",
      "Clock model Fixed probability":"0.5",

      "AA":"0.25",
      "AG":"0.25",
      "AT":"0.25",
      "AC":"0.25",

      "GA":"0.25",
      "GG":"0.25",
      "GT":"0.25",
      "GC":"0.25",

      "TT":"0.25",
      "TC":"0.25",
      "TA":"0.25",
      "TG":"0.25",

      "CT":"0.25",
      "CC":"0.25",
      "CA":"0.25",
      "CG":"0.25"
    },

    "Hotspot 2":{

      "Region":"10000_20000",
      "Clock model":"Negative Binomial",

      "Clock model Poisson mean":"2",
      "Clock model Negative Binomial sucesses":"10",
      "Clock model Negative Binomial probability":"0.5",
      "Clock model Fixed probability":"0.5",

      "AA":"0.25",
      "AG":"0.25",
      "AT":"0.25",
      "AC":"0.25",

      "GA":"0.25",
      "GG":"0.25",
      "GT":"0.25",
      "GC":"0.25",

      "TT":"0.25",
      "TC":"0.25",
      "TA":"0.25",
      "TG":"0.25",

      "CT":"0.10",
      "CC":"0.40",
      "CA":"0.20",
      "CG":"0.30"
    },
  },

  "Recombination":{

    "Number of hotspots":3,
    "Recombination profiles folder":"/sequence_Profiles/recombination_profiles",

    "Hotspot 1":{
      "Region":"100_200"
      "Reference probability of recombination":"0.5",
      "Reference selectivity":"1",
    },

    "Hotspot 2":{
      "Region":"10_50"
      "Reference probability of recombination":"0.75",
      "Reference selectivity":"1",
    },

    "Hotspot 3":{
      "Region":"10000_20000"
      "Reference probability of recombination":"0.05",
      "Reference selectivity":"1",
    },
  },

}
```

**Figure S - 10.** Configured sequence parameter file complete with two mutational hotspots and three recombination hotspots as well as configurations for the transmission bottleneck.

##### 3.5. Summary of simulator parametrization

The user is capable of setting up a complex simulation via the parametrization of four main parameter files. The organisation into four separate files is conducted to make it simpler for the user to organise their simulation set up and if needed conduct varying simulations with changes to only the required file.

For the user's convenience parametrisation does not have to be declared in order. As long as the nesting of parameters is done appropriately the parameters can be present anywhere in the file in no particular order. Apollo will seek the relevant parameters required for the simulation and conduct the analysis. However, the declaration of the parameter is case sensitive, but the variable assignment is not, and the user can assign variables at will regardless on the letter casing.

If any parameters are missing during the execution of the simulation, Apollo will notify the user of the exact parameter that is missing complete with the proper declaration method. The user only needs to point the software to the Master parameter file and through it, Apollo will find the rest of the parameter files including additional auxiliary files that will be needed for the simulation.

With that we complete the setup and execution of Apollo for a simulation. The following sections will explore the theoretical implementations implemented in the software including the statistical methods used in relation to population genetics and within host evolution.

#### SECTION 4. PARAMETERIZATION OF COMPLIMENTARY TOOLS

In this section we will look into the parameterization of the four complimentary tools in Apollo. For all tools these tools except for base substitution model to JSON and recombination hotspots to JSON an Apollo simulation has to have been completely run.

##### 4.1. Haplotype retriever

The haplotype retriever is used to retrieve all unique sequences that were generated in the simulation of a given host.

The function only requires one parameter in addition to the hardware parameters. Do not need to specify a GPU.

###### 4.1.1. Haplotype Node IDs

Defines the infected host from whom the unique haplotypes have to be retrieved from.

E.g.: *"Haplotype Node IDs":"8_8"*

The function provides complete information of each unique haplotype such as: tissue of occurrence, generation of occurrence, complete sequence, number of times the haplotype occurred in a given generation in a given tissue, and frequency of occurrence in a given generation in a given tissue.

Within the results directory under the node_Data directory the function creates three-tab delimited csv files within the query host's folder. The three generated files are as follows:

1. all_Haplotype_Frequencies.csv
   Contains information of all generated unique haplotypes within the query host
2. alive_Haplotype_Frequencies.csv
   Contains information of all haplotypes that survived and did not read mortality before the end of the generation.
3. parent_Haplotype_Frequencies.csv
   Contains information of haplotypes that belonged to virions that became parents and produced progeny.

##### 4.2. Pedigree retriever

Provided a target sequence, the host/ node, tissue and generation the pedigree retrieval function is capable of providing the complete list of ancestral virions and their sequences that led to the current sequence. The function is able to find all virions in the given location matching the query sequence and provides the complete pedigree for each of them. In addition to the hardware parameters the function needs five parameters to run. Do not need to specify a GPU.

###### 4.2.1. Pedigree Node ID

Defines the infected host who contains sequences matching the query sequence.

E.g.: *"Pedigree Node ID":"0_0"*

###### 4.2.2. Pedigree Tissue

Defines the tissues within the infected host containing the matching sequences.

E.g.: *"Pedigree Tissue":"Plasma"*

###### 4.2.3. Pedigree Generation

Defines the generational point with sequences matching the query.

E.g.: *"Pedigree Generation":52*

###### 4.2.4. Pedigree Sequence

The target sequence whose pedigree we wish to obtain. We provide it in the form of a FASTA file.

E.g.: *"Pedigree Sequence":"test_Folder/pedigree/one.fasta"*

###### 4.2.5. Pedigree get Sequences

A Boolean function with values of either “YES” or “NO”. If Yes then the pedigree virions sequences are also retrieved else only the sequence information and parent progeny relationships are provided.

E.g.: *"Pedigree get Sequences":"Yes"*

At all times the pedigree retriever provides two tab delimited csv outputs for each sequence that matches the query. The two files are:

1. pedigree_Relationships.csv
   Provides information of the parent progeny relationships and type of parent. Type of parent refers to if the parent is the template parent or a recombinant parent.
2. sequence_Information.csv
   Provides the sequence information of each ancestral virion such as the host it was found in, tissue and generation.

If the get sequence option is enabled, it creates a FASTA file containing the sequences of the ancestral virions.

##### 4.3. Segregating sites matcher

This function is similar to Haplotype retriever. But instead of searching the whole host for all unique sequences the segregating sites matcher attempts to identify sequences that match query sequences based on given segregating sites. In addition, the user can specify the coverage that the sequences should match. It requires four parameters in addition to the standard hardware parameters. Does not require a GPU.

###### 4.3.1. Segregating match sequences

Location of the FASTA file containing the query sequences.

E.g.: *"Segregating match sequences":"merged.fasta",*

###### 4.3.2. Segregating match Node ID

Defines the infected host who contains sequences matching the query sequence.

E.g.: *"Segregating match Node ID":"0_0"*

###### 4.3.3. Segregating match tissue

Defines the tissues within the infected host containing the matching sequences.

E.g.: *"Segregating match tissue":"Plasma"*

###### 4.3.4. Segregating match cutoff

Defines the coverage of the matching sequences to the query sequences. Provided as a ratio between 0 and 1. 0 being non of the sites match and 1 being a complete match.

E.g.: *"Segregating match cutoff":"0.96875"*

The segregating site matcher will create a folder called seg_Match within the nodes results and provide a tab delimited csv with the complete information of the virions with matching sequences. The information collected include the sequence, tissue and generation of occurrence along with information of the matching degree and if any the mismatched bases.

##### 4.4. Base substitution model to JSON

Converts a csv file with the mutational hotspots' base substitution model to Apollo's JSON format. The file can be either comma (,) or tab delimited. Each row in the file should be hotspot. The file should contain a header and the 24 headings should be as shows in **Table S - 6**.

**Table S - 6.** Depiction of the columns that should be present for mutation along with their description.

| Column heading | Description |
|---|---|
| **Hotspot_index** | The mutational hotspot count |
| **Position_start** | The starting position of the hotspot in the genome. |
| **Position_stop** | The ending position of the hotspot in the genome. |
| **Clock_model** | The clock model being used. It can be either Poisson, Negative Binomial or Fixed. |
| **Poisson_mean** | If Poisson is entered for the clock model, then the mean of the distribution should be entered here. Else it should be left as NA. |
| **Negative_Binomial_Sucess** | If Negative Binomial is entered for the clock model, then the number of successes of the distribution should be entered here. Else it should be left as NA. |
| **Negative_Binomial_Probability** | If Negative Binomial is entered for the clock model, then the probability of success of the distribution should be entered here. Else it should be left as NA. |
| **Fixed_probability** | If Fixed is entered for the clock model, then the probability of observing a mutation per site should be entered here. Else it should be left as NA. |
| **AA** | Probability of base A mutating to base A |
| **AT** | Probability of base A mutating to base T |
| **AG** | Probability of base A mutating to base G |
| **AC** | Probability of base A mutating to base C |
| **TA** | Probability of base T mutating to base A |
| **TT** | Probability of base T mutating to base T |
| **TG** | Probability of base T mutating to base G |
| **TC** | Probability of base T mutating to base C |
| **GA** | Probability of base G mutating to base A |
| **GT** | Probability of base G mutating to base T |
| **GG** | Probability of base G mutating to base G |
| **GC** | Probability of base G mutating to base C |
| **CA** | Probability of base C mutating to base A |
| **CT** | Probability of base C mutating to base T |
| **CG** | Probability of base C mutating to base G |
| **CC** | Probability of base C mutating to base C |

Requires one additional parameter apart from the standard hardware parameters. Does not require a GPU.

##### 4.4.1. Convert file location

Provides the location of the csv file that contains the information of the mutational hotspots.

E.g.: *"Convert file location":"Mutation_Site_model.csv"*

##### 4.5. Recombination hotspots to JSON

Converts a csv file with the recombination hotspots' information to Apollo's JSON format. The file can be either comma (,) or tab delimited. Each row in the file should be hotspot. The file should contain five columns as detailed in

**Table S - 7.** Depiction of the columns that should be present for recombination along with their description.

| Column heading | Description |
|---|---|
| **Hotspot_ID** | The recombination hotspot's count. |
| **Start** | The starting position of the hotspot in the genome. |
| **Stop** | The ending position of the hotspot in the genome. |
| **Probability** | The reference probability of the region undergoing recombination. |
| **Selectivity** | The reference likelihood of the region being selected as the recombination template. |

Requires one additional parameter apart from the standard hardware parameters. Does not require a GPU.

###### 4.5.1. Convert file location

Provides the location of the csv file that contains the information of the recombination hotspots.

E.g.: *"Convert file location":"Recombination_Site_model.csv"*

#### SECTION 5. EXAMPLE OF SIMULATION CONFIGURATION

Here we will look at the configuration of a simulation that will encompass all aspects of Apollo's mechanisms. Any data sets used, and sample parameter files will be available via the CATE GitHub repository, where Apollo is housed.

All parameter file used in this example are housed in the folder called parameters_Apollo.

The simulation overview is as follows:

1. Hardware configuration of GPU, CPU, and SSDs will be conducted.
2. Dynamic Caveman model with five caves will be used to generate the contact network.
3. Three different host types will be present.
4. Hosts will have three different tissue types.
5. Cross-tissue migration will be present.
6. The genome will span 100 bases in length.
7. Two mutational hotspots and two recombination hotspots.
8. Fitness, Survivability, and Proof-reading profiles.
9. Sampling mechanism will be present.

Let's now look at how we will configure the above simulation scenario using the four parameter files.

##### 5.1. Configuring the master parameter file

###### 5.1.1. Hardware

We will set Apollo to use two CUDA devices (0 and 1), 10 CPU cores and be able to use the available SSD's multi read and write technology. On the GPU front we will configure the tool to process $10^6$ units at a time. The software will also process a maximum of 1000 cells at a time.

###### 5.1.2. Folder management

The output folder will be named *results*, the intermediate folder will be *intermediates* and the maximum number of virions an intermediate file can hold will be $10^6$. There will be no folder management in terms of folder compression.

###### 5.1.3. Simulation inception

Within the simulation the start date will be January 1$^{st}$, 2024. The simulation will run for a period of four months till April 1$^{st}$, 2024, and the first individual being infected will have a specific within host viral occupation within the tissues.

###### 5.1.4. Sub parameter file locations

The locations of the three remaining parameters files namely the network parameter file, host parameter file and sequence parameter file are named the network_test.json, node_Master.json and sequence_Master.json respectively.

An example of the parameter master file configured to suit the above discussed requirements is shown in **Figure S - 11**.

parameters_MASTER.json

```
{
  # Configuration of the Cuda device IDs accesible to Apollo
  # Here only one CUDA device is assigned for Apollo's use
  "CUDA Device IDs":"0",

  # Hardware configuration parameters
  "CPU cores":10,
  "GPU max units":100000,
  "Process cell rate":1000,
  "Multi read":"YES",

  # Output and Intermediate folder locations.
  "Output folders":"results",
  "Intermediate folders":"intermediates",

  # The maximum number of sequences that can be stored in an intermeiate file
  "Intermediate Sequences per file":1000,

  # Indicates the start date for the simulation
  "Start date":"2024-01-01",
  "First infection":"Random",

  # Indicate whether the user wants to run the simulation for a specified period of time
  "Stop after generations":"Yes",
  # Generations or Date can be used to stop the simulation as well
  "Mode to stop":"Date",
  "Number of generations":3,
  "End date":"2024-04-01",

  # Network profile parameter file location
  "Network profile":"/parameters_Cancer_test/network_Profiles/network_test.json",

  # Node profile parameter file location
  "Nodes master profile":"/parameters_Cancer_test/individual_Profiles/node_Master.json",

  # Sequence profiles parameter file location
  "Sequence master profile":"/parameters_Cancer_test/sequence_Profiles/sequence_Master.json",

  # Folder management controls
  # Compresses folders to a TAR.GZ or reduces them to a single TAR file
  "Enable folder management":"No",
  "Compress folders":"No",
}
```

**Figure S - 11.** Example of the configured master parameter file called “parameters_Master.json”. The file has been configured according to the requirements discussed in the example.

##### 5.2. Configuring the network parameter file

As mentioned in Section 3.2 on the configuration of the parameter file, each network type has its own unique set of parameters. Since our example utilizes a Dynamic Caveman model with five caves in the system we will look into parameterization.

##### 5.3. Characteristics of the network

For this test let us create a network with five cohorts (caves) with the number of nodes in each cave being determined by a negative binomial distribution. Of the total nodes in each cave 25% will be able to form connection with other immediately neighbouring cohorts and 55% of these nodes will be able to form connections with nodes from distant cohorts (**Figure S - 12**).

network_test.json

```
{
  # Select network type

  # Random Model is selected by typing Random model
  # Barabasi Albert Model is selected by typing BA model
  # Standard Caveman Model is selected by typing SC model
  # Dynamic Caveman Model is selected by typing DC model
  # Erdos and Renyi Random Model is selected by ER model

  "Network type":"DC model",

  # Total number of caves that will be present in the DC network model
  "DC_model number of caves":5,

  # Determine the number of nodes that will be present in each cave in the network
  # Negative binomial, every caves number of nodes will determined by a NEGATIVE BINOMIAL distribution.
  # Poisson, every caves number of nodes will determined by a POISSON distribution.
  "DC_model node distribution":"Negative binomial",

  # Assignment of parameter values; sucesses and probability for DC model with NEGATIVE BINOMIAL for per cave node distributions.
  "DC_model node Negative binomial sucesses":10,
  "DC_model node Negative binomial probability":"0.55",

  # Assignment of parameter values; mean for DC model with POISSON for per cave node distributions.
  "DC_model node Poisson mean":"1",

  # Determine the percentage of nodes that will take part in forming neighbouring node attachments and from them how many will form global attachments.
  "DC_model neighbouring nodes percentage":"0.25",
  "DC_model global nodes percentage":"0.55"
}
```

**Figure S - 12.** Example of configured network parameters file

##### 5.4. Configuration of the nodes master file and nodes profile

For these files along with the rest, sample files in line with the described case study are provided in the GitHub repo in the folder parameters_Apollo.

##### 5.5. Configuration of sequence master file

Here too the relevant parameter files are available under the parameters_Apollo folder.
